# Resource economics of tree communities control soil food web multifunctionality in European forests

**DOI:** 10.1101/2025.02.19.639063

**Authors:** Ludovic Henneron, David A. Wardle, Matty P. Berg, Stephan Hättenschwiler, Jürgen Bauhus, François Buscot, Sylvain Coq, Thibaud Decaëns, Nathalie Fromin, Pierre Ganault, Lauren M. Gillespie, Kezia Goldmann, Radim Matula, Alexandru Milcu, Bart Muys, Johanne Nahmani, Luis Daniel Prada-Salcedo, Michael Scherer-Lorenzen, Kris Verheyen, Janna Wambsganss, Paul Kardol

**Affiliations:** Department of Forest Ecology and Management, Swedish University of Agricultural Sciences, Umeå, Sweden; ECODIV USC 1499, Univ Rouen Normandie, INRAE, Rouen, France; Department of Ecology and Environmental Science, Umeå University, Umeå, Sweden; Amsterdam Institute for Life and Environment, Section Ecology and Evolution, Vrije Universiteit, Amsterdam, the Netherlands; GELIFES, Community Conservation Group, Groningen University, Groningen, The Netherlands; CEFE, Univ Montpellier, CNRS, EPHE, IRD, Montpellier, France; Chair of Silviculture, Faculty of Environment and Natural Resources, University of Freiburg, Freiburg, Germany; Department of Soil Ecology, UFZ - Helmholtz Centre for Environmental Research, Halle (Saale), Germany; German Centre for Integrative Biodiversity Research (IDiv), Leipzig, Germany; Department of Forest Ecology, Faculty of Forestry and Wood Sciences, Czech University of Life Sciences, Prague, Czech Republic; Montpellier European Ecotron, Univ Montpellier, CNRS, Campus Baillarguet, Montferrier-sur-Lez, France; Department of Earth & Environmental Sciences, KU Leuven, Leuven, Belgium; Geobotany, Faculty of Biology, University of Freiburg, Freiburg, Germany; Forest & Nature Lab, Department of Environment, Ghent University, Melle-Gontrode, Belgium; Research Institute for Forest Ecology and Forestry (FAWF), Landesforsten Rheinland-Pfalz, Trippstadt, Germany; Department of Forest Mycology and Plant Pathology, Swedish University of Agricultural Sciences, Uppsala, Sweden

## Abstract

Plants affect terrestrial ecosystem functioning by performing the primary production that energetically sustains heterotrophic organisms^1^, and by shaping the microenvironment^2^. However, the influence of plant diversity and community composition on ecosystem functioning through their effects on energy flow into food webs has been little studied^3,4^, especially for soil food webs that channel most of the plant-derived energy^1,5^. Applying a food web energetics approach^6,7^, we show that the resource economics of dominant tree species control soil food web multifunctionality across European forests. Specifically, tree communities dominated by resource acquisitive species promoted faster rates of multiple soil trophic functions simultaneously than did those dominated by resource conservative species. This was primarily driven by their production of plant litter with higher nutritional quality and their warmer forest microclimate, leading to a higher metabolic activity of soil organisms^8^. Tree species mixing had rather weak and negative effects on soil food web multifunctionality, mostly due to a shift in the resource-based energy channeling from living plant fine roots to litter and a cooling effect on the forest microclimate. Tree diversity effects were largely outweighed by community compositional effects, which were of similar magnitude to the effects of biogeographic differences among locations. Our findings emphasize the importance of plant functional traits related to resource economics as drivers of plant community effects on soil food web functioning^5,9^ and highlight the consequences that climate-driven shifts in tree community composition could have for forest soil functioning.

## Main text

Motivated by growing concerns that ongoing biodiversity change could compromise the provisioning of ecosystem services vital to humanity^10,11^, a large number of experiments have shown that local species diversity enhances multiple ecosystem processes across trophic levels and habitats, both individually^12^ and simultaneously^13^ (*i.e.* multifunctionality). However, most research linking biodiversity to ecosystem functioning (BEF) has focused on single trophic levels or simple food chains, despite the need for a multitrophic perspective to better understand BEF relationships and their relevance to real-world ecosystems^4,14^. In terrestrial ecosystems, primary productivity has commonly been found to be enhanced by plant diversity^3^, in addition to be affected by community composition^15^. Multitrophic BEF studies further demonstrated that plant diversity and composition can in turn greatly affect consumer communities across trophic levels, especially in aboveground producer-based (‘green’) food webs^16–18^. Nonetheless, the broader influence of plant diversity and composition on terrestrial ecosystem functioning through cascading effects across trophic levels remain much less explored, as few studies have explicitly incorporated a quantitative food web approach^3,4^.

Trophic interactions play a major role in underpinning ecosystem functioning because heterotrophic organisms regulate multiple ecosystem processes primarily through their food consumption^6^. Accordingly, quantifying energy flow along trophic links in food webs^19,20^ has been proposed as a powerful tool to mechanistically understand BEF relationships in complex multitrophic systems^4,6^. Energy fluxes in food webs reflect trophic functions such as herbivory, detritivory, microbivory, and predation (Fig. 1), and therefore provide a common currency for assessing ecosystem functioning across trophic levels^6^. These trophic functions can be combined to measure food web multifunctionality, which represents the simultaneous performance of multiple trophic functions carried out by the food web^6,7^. However, the application of food web energetics to BEF research remains in its infancy^21,22^, particularly for belowground food webs that play a critical role in ecosystem functioning^5,7,23^.

**Fig. 1.**
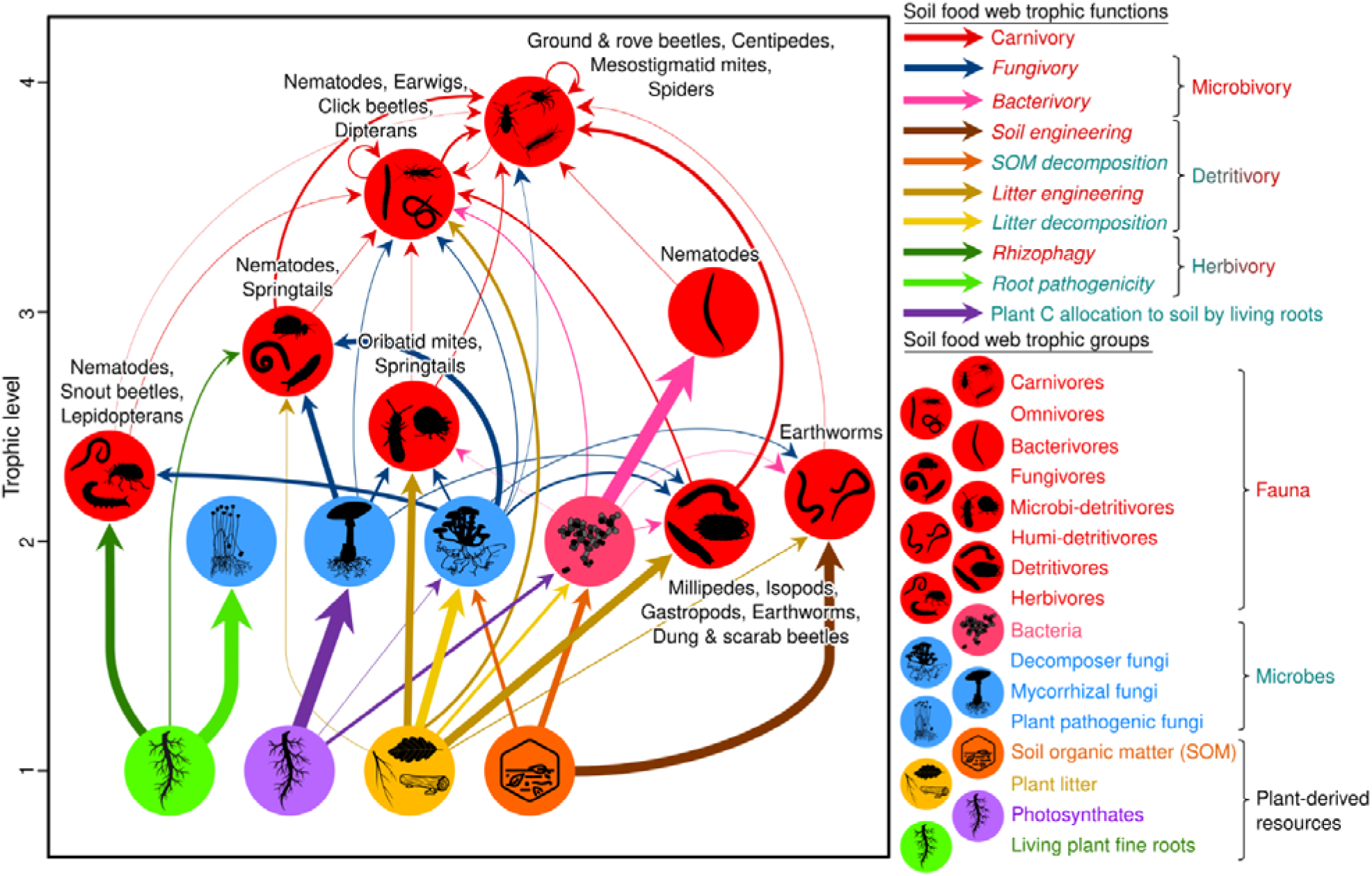
Graphical representation of the trophic functions and topography of the soil food web. Dominant taxonomic groups in faunal trophic groups are shown with text within the figure. Arrows indicate energy flows among trophic groups, with arrow widths corresponding to trophic interaction strengths. Arrow colours show how energy fluxes were aggregated by resource and consumer types to quantify trophic functions of the soil food web. These are related to five broad trophic functions: carnivory, *i.e.* the consumption of fauna; microbivory, *i.e.* the consumption of microbes (‘bacterivory’ and ‘fungivory’); plant C allocation to soil by living roots, *i.e.* the consumption of photosynthates and rhizodeposits from plant roots by mycorrhizal fungi and decomposer microbes; herbivory, *i.e.* the consumption of plant roots by microbes (‘root pathogenicity’) or fauna (‘rhizophagy’); and detritivory, *i.e.* the consumption of detritus (‘plant litter’ or ‘soil organic matter’, SOM) by microbes (‘decomposition’) or fauna (‘engineering’). Decomposition refers to the assimilation and mineralization of dead organic matter through respiration. Engineering refers to the physical modification, maintenance, or creation of habitats by feeding activity (Methods). Trophic interaction strengths shown here are based on biomass data averaged across all plots, but plot-specific biomass data was used to calculate trophic interaction strengths for each local food web. To simplify the representation, trophic guilds (finer level) were aggregated into trophic group (coarser level) by averaging their trophic interaction strengths, weighted by the relative biomasses of the corresponding guilds within each group. See Supplementary Fig. 4 for a more detailed illustration of the food web topology and trophic interaction strengths among trophic guilds. Silhouette are in the public domain under the Creative Commons Attribution License ‘CC BY’ (source: https://www.phylopic.org/).

In terrestrial ecosystems, most of the energy derived from primary production directly enters the soil as freshly dead organic matter (*i.e.* plant litter)^1^. Consequently, the dominant channel of energy flow is through the detritus-based (‘brown’) food web^5^, which underpins decomposition and nutrient cycling processes and ultimately controls the availability of growth-limiting nutrients to plant roots^24^. It also allows photosynthetically fixed carbon (C) to be stabilized in the soil or released back to the atmosphere by respiration, with major implications for ecosystem C cycling and sequestration^25^. Additionally, terrestrial ecosystems host an important belowground living root-based food web that involves root herbivory and plant C allocation to mycorrhizal fungi and rhizosphere microbes^5^, and which strongly influences the size and turnover of root biomass as well as the acquisition of belowground resources by plant roots^24^.

Plant community composition is well known to exert a strong control on ecosystem processes related to the soil food web^10^. This has traditionally been attributed to the functional traits of dominant plant species that are directly involved in the acquisition, processing, and conservation of resources (*i.e.* economic traits^9^), and which shape the quantity and nutritional quality of trophic resources such as litter^5^ and rhizodeposits^26^. However, empirical evidence that explicitly relates plant economic traits at the community-level to the functioning of soil food webs is still lacking. Plant diversity is generally thought to promote soil processes by increasing the quantity and diversity of trophic resources and by fostering favourable micro-environmental conditions^27^. Accordingly, plant diversity has recently been shown to foster multitrophic energy flow and storage across both above- and belowground food webs in a grassland biodiversity experiment^22^. Nevertheless, how plant diversity and composition affect soil food web functioning in real-world ecosystems has been seldom explored.

Here, we investigated how the diversity and composition of naturally assembled tree communities influence multiple soil trophic functions across a variety of environmental contexts in European forests. We hypothesized that: (1) tree diversity positively affects soil food web multifunctionality, and (2) tree composition exerts a strong effect on soil food web multifunctionality owing to the key role of the economic traits of dominant tree species. We tested these hypotheses using a pan-European network of 64 mature forest plots distributed across four geographic locations in different countries (Finland, Poland, Romania, and Italy), spanning boreal to Mediterranean climates and representing highly contrasting European forest types. Within each location, we followed a stratified sampling design by selecting three-species mixture stands with varying tree species compositions, along with corresponding monoculture stands (Extended Table 1, Methods). This plot selection was performed so as to minimize covariation with potential confounding factors (Supplementary Fig. 1a), and thereby increase our ability to infer relationships between the diversity and composition of tree communities and soil food web functioning.

To quantify energy fluxes in the belowground food web of each plot, we applied steady-state food web modelling based on ecosystem energetics^6^. This approach is grounded in the principle that, for a given trophic group in a food web, energy uptake by food consumption must balance energetic demands, including energy lost during food assimilation, and by metabolism (respiration^8^) and predation. We first quantified the biomass, metabolism and assimilation efficiency of major groups of soil organisms, including microbes and fauna, along with the biomass of plant-derived resources, *i.e.* living plant fine roots and associated photosynthates, plant litter, and soil organic matter (SOM). We then gathered soil organisms into trophic groups, established food web topology, and quantified trophic interaction strengths to allow the calculation of energy fluxes^6,7^ (Fig. 1, Methods). As commonly observed, the biomasses and energy fluxes into the soil food web were much higher at lower trophic levels^19^, as well as for microbes than for fauna^28^ (Fig. 2a, Extended Data Fig. 1). Energy fluxes were then aggregated by resource and consumer types to quantify ten trophic functions of the soil food web^6,7^, *i.e.* plant C allocation to soil by living roots, root pathogenicity, rhizophagy, litter decomposition, litter engineering, SOM decomposition, soil engineering, bacterivory, fungivory, and carnivory (Fig. 1). Decomposition refers here to the assimilation and mineralization of dead organic matter through microbial respiration, while engineering refers to the physical modification, maintenance, or creation of habitats^2^ by faunal detritivory (see Methods for further justification). These ten trophic functions are directly linked to various aspects of ecosystem functioning, including the cycling of nutrients and their supply to plants, soil carbon cycling and sequestration, soil structure maintenance, and biocontrol through predation^5,7,23^. For those calculated trophic functions for which direct measurements of corresponding ecosystem processes were also available, we found good agreement between them (Extended data Fig. 2, Supplementary Results). We then quantified soil food web multifunctionality^6,7^ by averaging the range-standardized values of these ten trophic functions.

**Fig. 2.**
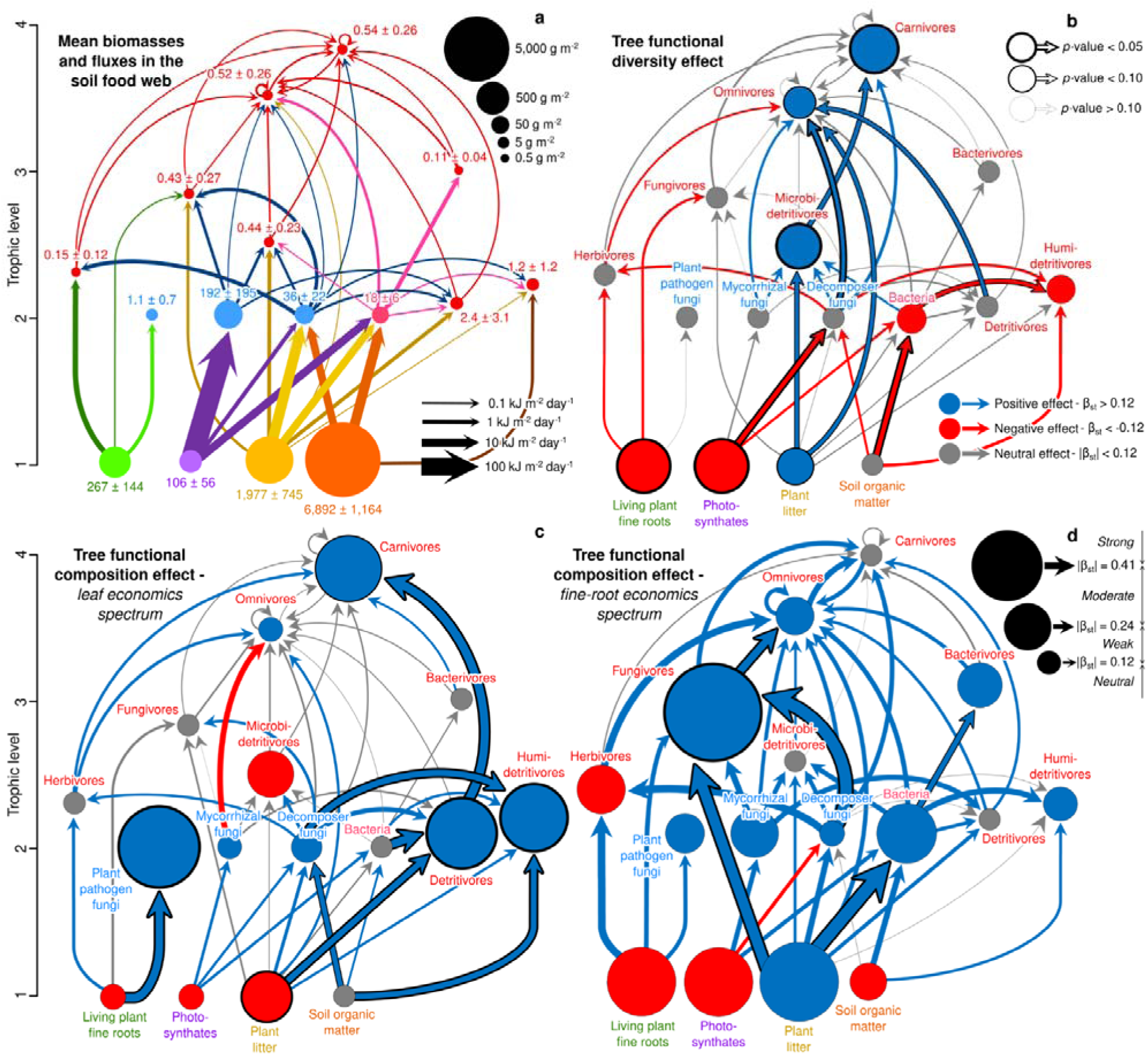
Effects of tree functional diversity and composition on energy fluxes and trophic group biomasses of the soil food web across European forests. **a**, Mean trophic group biomasses (circles, g dry weight m^-2^ ± 95% CI) and energy fluxes (arrows, kJ m^-2^ day^-1^) of the soil food web are shown as predicted for average conditions (mean values for each predictor). Circle areas and arrow widths are scaled by the cube root of biomasses and energy fluxes, respectively. Trophic group names are shown in other panels. The colours used for trophic functions and groups are the same as in Fig. 1. **b-d**, Effects of tree functional diversity (**b**) and composition (**c**, leaf economics spectrum; **d**, fine-root economics spectrum) on trophic group biomasses and energy fluxes of the soil food web. Blue, red and grey arrow indicate respectively positive, negative and neutral effects. Circle areas and arrows widths are scaled by effect sizes, that are partial slope regression coefficients standardized by standard deviation (β_st_), derived from Bayesian linear mixed-effect models (*n* = 64, Methods).

To test our hypotheses, we performed Bayesian multi-level modelling assessing how the diversity and composition of tree communities affect soil food web multifunctionality, while also statistically controlling for potential confounding factors (Methods). Since most BEF studies have focused on the effects of species diversity and composition, we first adopted a traditional taxonomic approach to evaluate the effect of tree species richness (mixing of three species) and quantify the relative importance of tree species richness, tree species composition (combination of species), and biogeography, *i.e.* variation in abiotic conditions and associated tree species turnover across locations (Supplementary Fig. 1a). To gain deeper mechanistic insights into BEF relationships^14,29^, we also adopted a functional (trait-based) approach assessing the effects of both functional diversity and functional composition of tree communities, along with biogeographic differences among locations. This enabled us to evaluate support for the ‘functional diversity’ hypothesis, which states that functionally diverse communities promote ecosystem functioning due to species complementarity mechanisms^29^, compared to the ‘mass-ratio’ hypothesis, which states that ecosystem functioning is primarily driven by the attributes (trait values) of dominant species and their relative abundances^30^. We characterized tree functional diversity and composition by using three leaf traits and six fine-root traits commonly used to describe ecological strategies related to the plant economics spectrum^9,31^, and which reflect the trade-off between resource acquisition, *i.e.* fast strategy, and resource conservation, *i.e.* slow strategy (Methods). Functional diversity was measured as the variability in trait values among tree species (functional dispersion), while functional composition was measured using the community-weighted mean (CWM) of trait values. We then reduced the complexity of tree functional composition to two dimensions (Extended data Fig. 3a). The first dimension corresponded to the leaf economics spectrum (LES), which ranges from slow/conservative to fast/acquisitive leaf attributes^9^, and was also aligned here with a fine-root gradient which ranges from low to high belowground resource foraging efficiency. The second dimension corresponded to the fine-root economics spectrum (RES), which ranges from slow/conservative to fast/acquisitive fine-root attributes, and was also aligned here with a fine-root gradient of soil exploration strategies which ranges from ‘do-it-yourself’ to ‘outsourcing’ attributes^31^. To investigate the mechanisms underlying the effects of tree diversity and composition on soil food web multifunctionality, we performed structural equation modelling that incorporates a wide range of environmental drivers such as tree and understorey vegetation, microclimate, tree leaf litter quality, and soil fertility (Extended data Fig. 4).

### Effects of tree diversity on soil food web multifunctionality

Contrary to our first hypothesis, we found soil food web functioning to be little affected by tree species mixing, *i.e.* one *versus* three tree species present (Fig. 3). Tree species richness had rather negative effects on many trophic functions of the soil food web, *i.e.* moderately negative for SOM decomposition, and weakly negative for plant C allocation to soil by living roots, rhizophagy, and soil engineering (Fig. 3a, Supplementary Fig. 2). Tree species richness also had a weakly positive effect on litter engineering, but neutral effects on the five remaining trophic functions. Similar results were observed for tree functional diversity (Fig. 2b, Fig. 3c), which is in contrast to what would be expected based on the ‘functional diversity’ hypothesis^29^. Accordingly, tree diversity had a weakly negative effect on soil food web multifunctionality (Fig. 3a,c), which was mediated through multiple mechanisms.

**Fig. 3.**
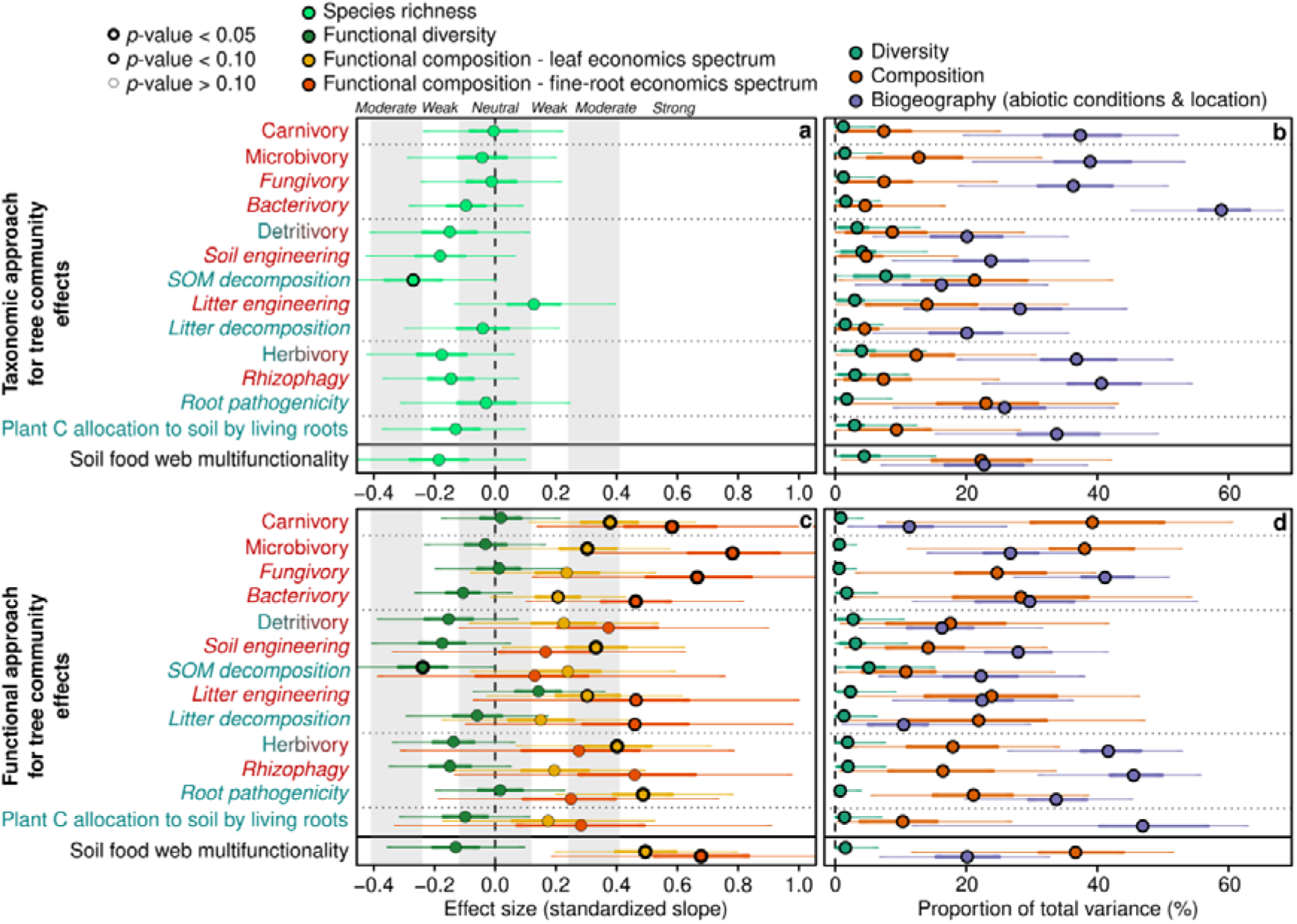
Effects of tree diversity and composition on trophic functions and multifunctionality of the soil food web across European forests. **a**,**c**, Effect sizes of tree diversity and composition calculated as partial slope regression coefficients, standardized by standard deviation. Circles represent the mean effect sizes, with thick and thin error bars representing 50% and 95% credible intervals, respectively. **b**,**d**, Decomposition of the total variance explained by tree diversity and composition, and biogeography (*i.e.* abiotic conditions and location), as determined by variance partitioning. Effects of diversity and composition of tree communities were estimated using both taxonomic (**a**,**b**) and functional (**c**,**d**) approaches based on Bayesian linear mixed-effect models (*n* = 64, Methods). Both the leaf and fine-root economics spectra representing tree functional composition ranged from slow/conservative to fast/acquisitive attributes (Extended Data Fig. 3a). Trophic functions are based on energy fluxes in the soil food web, aggregated by resource and consumer types (Fig. 1). Italicized function are subsets of broader functions (not italicized) between the dotted lines. Trophic functions in blue-green and dark red indicated those mediated by microbes and fauna, respectively. Soil food web multifunctionality is the average of standardized values of each of the ten trophic functions of the soil food web. For statistical results, see Extended Data Table 2.

First, tree diversity negatively affected soil food web multifunctionality by both decreasing root biomass and increasing tree litterfall (Fig. 4), which shifted the resource-based energy channeling in the soil food web from living plant fine roots to litter (Fig. 2b, Supplementary Fig. 2). Previous studies have emphasized the key importance of living plant roots for soil microbes^32,33^ and fauna^34,35^, primarily through the provision of readily available resources such as rhizodeposits and photosynthates^33^. This finding therefore suggests that the diversity-induced shift in tree resource allocation from below-ground to above-ground^36,37^ modifies the overall quality of trophic resources for soil organisms towards less readily available energy, thereby impairing soil food web multifunctionality.

**Fig. 4.**
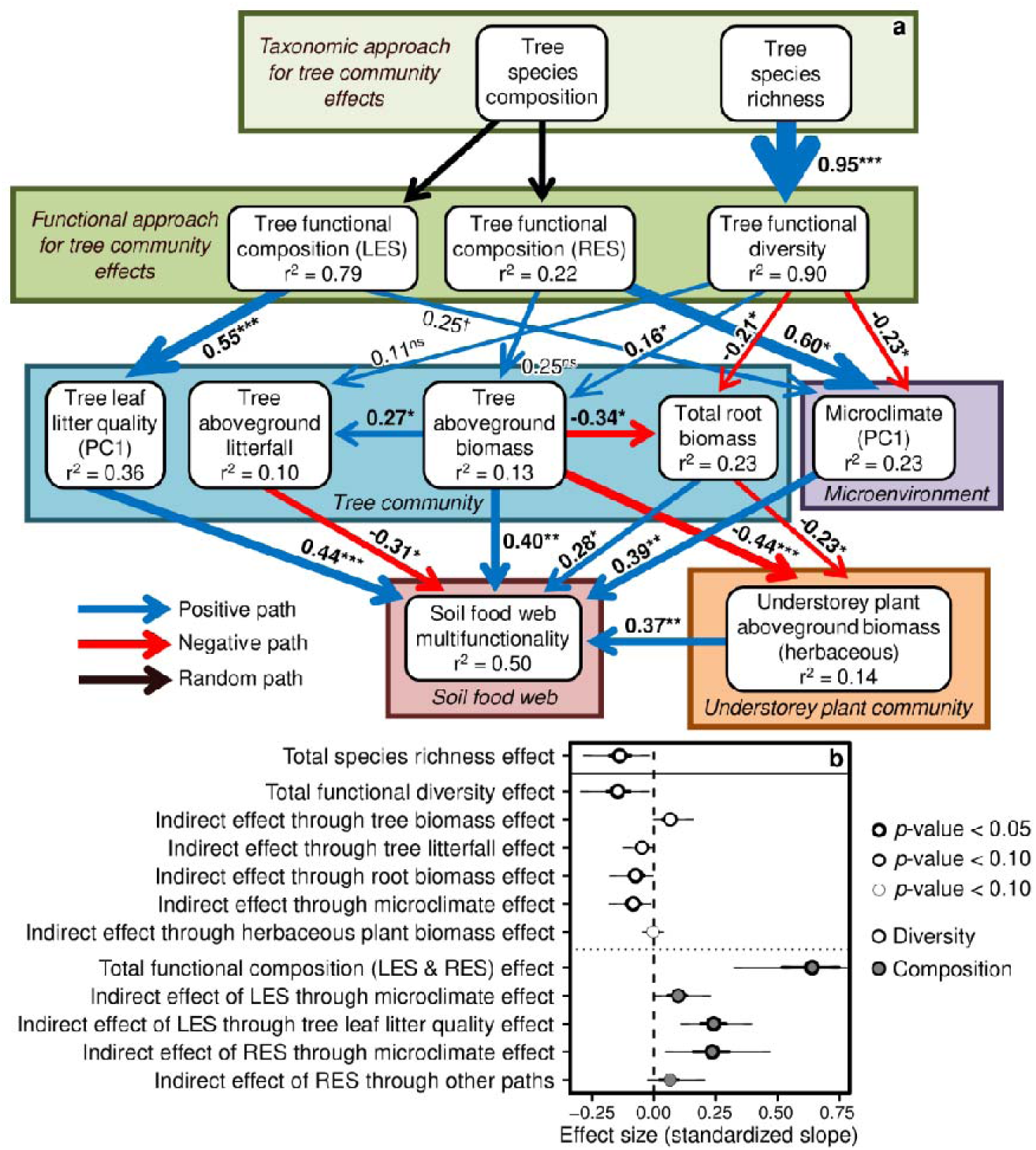
Mechanistic insights into the mediation of tree diversity and composition effects on soil food web multifunctionality across European forests. **a**, Best-supported Bayesian multi-level SEM model (*n* = 64). Arrows indicate the direction of causality, and their widths are proportional to the path coefficients (slopes standardized by standard deviation). The r^2^ values shown inside the boxes are the proportions of variance explained by explanatory variables present in this illustration. Both the leaf and fine-root economics spectra (LES and RES, respectively) representing tree functional composition ranged from slow/conservative to fast/acquisitive attributes (Extended Data Fig. 3a). ‘Microclimate’ ranged from colder and more humid to warmer and less humid (Extended data Fig. 3b). ‘Tree leaf litter quality’ ranged from low to high nutritional quality (Extended data Fig. 3c). The model was well supported by the data (Fisher’s C = 79.5, df = 96, *p*-value = 0.888). None of the independence claims implied by the model were statistically significant at α = 0.05 (Supplementary Table 7). Relationships with abiotic context variables, and correlated errors between variables, are not shown here for clarity. See 0additional statistical results. Significant effects (*p* < 0.05) are reported in bold. ^ns^, *p* > 0.10; ^†^, *p* < 0.10; *, *p* < 0.050; **, *p* < 0.010; ***, *p* < 0.001. **b**, Effect sizes of the indirect and total effects of tree diversity (white circles) and composition (grey circles) on soil food web multifunctionality, mediated by changes in tree and understorey plant communities, and microenvironment. Circles represent the mean effect sizes, with thick and thin error bars representing 50% and 95% credible intervals, respectively.

Second, tree diversity negatively affected soil food web multifunctionality by cooling the forest microclimate, leading to a decrease in mean soil and air temperature (Fig. 4, Extended data Fig. 3b), which could be due to enhanced canopy packing^38,39^. According to the ‘metabolic theory of ecology’^8^, metabolic rates of organisms are highly temperature-dependent. Consequently, the buffering of forest microclimate and associated decrease in mean temperature by tree diversity^40,41^ can lead to a decline in the energy demand of soil organisms, in turn reducing energy fluxes in the soil food web. This finding provides evidence for the emerging view that microclimate modulation can be an important, though often overlooked, mechanism contributing to BEF relationships in terrestrial ecosystems^38^.

Overall, our results sharply contrast with the results from most BEF experiments, which are typically based on randomly assembled communities and show positive effects of diversity on ecosystem functions^12,13^, especially for plant diversity^3,22^. However, in naturally assembled communities, the strength and direction of these diversity effects have been found to vary greatly^42^. While positive BEF relationships are often observed, neutral or even negative relationships are also common^42^. Our findings contribute to provide empirical evidence for the emerging theoretical view that positive BEF relationships should not necessarily be expected in real-world ecosystems because biotic interactions filter species during community assembly, which means that there can be a large effect of initial diversity on ecosystem functioning even with low observed local diversity^43^.

### Effects of tree composition on soil food web multifunctionality

Our data show that tree species composition had a much larger influence on soil food web multifunctionality than did tree species mixing *per se*. The effect of species composition was over five times stronger than that of species richness, accounting for 22.3% *versus* 4.4% of the total variance respectively (Fig. 3b). The effect of species composition within geographic locations was of similar magnitude as that of biogeography, which explained 22.7% of the variance. Tree species varied substantially in their effects, with species such as *Ostrya carpinifolia*, *Betula pendula*/*pubescens*, and *Carpinus betulus* having positive effects and *Pinus sylvestris* having negative effects on soil food web multifunctionality (Extended Data Fig. 5). Tree compositional effects were especially strong for trophic functions such as root pathogenicity, litter engineering, and SOM decomposition (Fig. 3b).

In line with the ‘mass-ratio’ hypothesis^30^, the functional composition of tree communities corresponding to community-level economic traits explained an even larger portion of the variance in soil food web multifunctionality than did their species composition alone (36.7% *versus* 22.3%, Fig. 2b,d). This is likely because functional composition takes into account the relative abundance of tree species and the variability in trait values across geographic locations, which species composition (combinations of species present) does not^29^. Functional composition also explained a larger portion of variance when compared to biogeography (20.1% of variance) and functional diversity (1.6%, Fig. 3d). This result emphasizes the importance of plant traits, in tandem with abiotic factors, for the functioning of real-world ecosystems^42^.

In accordance with our second hypothesis, both dimensions of functional composition, *i.e.* the leaf and fine-root economics spectra, had strong effects on soil food web multifunctionality. Specifically, we found that tree communities dominated by species with fast/acquisitive attributes promoted soil food web multifunctionality relative to those dominated by species with slow/conservative attributes (Fig. 2c). This finding provides empirical support for the theoretical framework which suggests that trait-driven plant strategies related to their resource economics at community-level control the functioning of soil food webs^5^. Positive effects of acquisitive attributes were observed across all trophic functions, particularly for lower-level trophic functions such as root pathogenicity and litter engineering, as well as for higher-level trophic functions such as bacterivory, fungivory, and carnivory (Fig. 3c,d). This contrasts sharply with earlier evidence from experimental grasslands, where plant diversity effects on consumer communities have been shown to weaken with increasing trophic level^16^.

The positive effects of leaf acquisitive attributes of tree communities were most pronounced for root pathogenicity, litter and soil engineering, and carnivory (Fig. 2c, Fig. 3c), and we observed that the effect of the leaf economics spectrum on soil food web multifunctionality was mostly mediated by the quality of tree leaf litter (Fig. 4, Extended data Fig. 3c). Specifically, tree communities with acquisitive leaf attributes produced leaf litter of higher nutritional quality, which in turn promoted greater soil food web multifunctionality. There is broad consensus that litter quality plays a key role as a major driver of soil food webs^5,44^, and our study provides empirical evidence that one of the primary ways that plant economic traits control soil food web functioning is through affecting the nutritional quality of the litter they produce^1,5^.

The positive effects of fine-root acquisitive attributes of tree communities on trophic functions were particularly strong for rhizophagy, litter decomposition, litter engineering, bacterivory, fungivory, and carnivory (Fig. 2c, Fig. 3c). Interestingly, we found that the effect of the fine-root economics spectrum on soil food web multifunctionality was mostly mediated by the forest microclimate, and this was also the case to a lower extent for the effects of leaf economics spectrum (Fig. 4, Extended data Fig. 3b). Specifically, tree communities with acquisitive attributes were associated with a warmer and less humid microclimate, *i.e.* higher mean soil and air temperature and lower mean soil moisture. This could potentially be explained by a lower interception of solar radiation by their canopies and a higher water uptake by their roots^9,45,46^, the latter leading to a lower cooling effect of water evaporation (as an endothermic process) into the soil^41^. A ‘warmer’ forest microclimate can in turn promote soil food web multifunctionality by increasing the metabolic activity of soil organisms^8^. This finding emphasizes that plant economic traits can also control soil food web functioning indirectly through their effects on micro-environmental conditions^2,38^. Through an examination of the effects of individual tree CWM traits, we found that soil food web multifunctionality was best explained by fine-root nitrogen content, which had a strongly positive effect that explained 31.4% of the total variance (Extended data Fig. 6). This result suggests that the positive effect of fine-root acquisitive attributes of tree communities may also be linked to the higher nutritional quality of root litter^1,5^. Additionally, fine roots of acquisitive plant species that have a higher nitrogen content and are more metabolically active are known to be associated with increased rhizodeposition rates, which in turn promotes greater stimulation of microbial activity in the rhizosphere^26,47^.

### Conclusions

Our study provides empirical evidence that tree communities control the functioning of belowground food webs across European forests, mainly through compositional effects related to the leaf and fine-root economic traits of dominant tree species. We found that tree communities dominated by species with fast/acquisitive attributes promoted soil food web multifunctionality compared to those dominated by species with slow/conservative attributes, mainly due to the higher nutritional quality of the trophic resources they provide and their warmer forest microclimate. Both the species and functional composition of tree communities had large effects on soil food web multifunctionality, comparable in magnitude to the effects of biogeographic differences among locations. In contrast, the effects of tree species mixing were much smaller and, generally, negative rather than positive, mostly due to a shift in the resource-based energy channeling from living plant fine roots to litter, as well as a cooling effect on forest microclimate. These findings emphasize the importance of tree species selection based on their economic traits rather than tree species mixing *per se* for the functioning of forest ecosystems.

Altogether, our findings highlight the value of the food web energetics and functional (trait-based) approaches to better understand and predict future shifts in terrestrial ecosystem functioning in the context of global change. For instance, climate change is expected to cause widespread tree mortality by intensifying drought and temperature stress in temperate forests^48,49^, particularly for tree species with acquisitive attributes^50^. In addition to the abrupt decline in ecosystem functioning following forest die-back until recovery, differential mortality of coexisting tree species can operate as a major driver of long-term shifts in tree community composition^49^. Specifically, drought-resistant conservative species could be expected to increasingly dominate tree communities in future scenarios that involve increasing drought intensity and forest management efforts to favour tree species better adapted to future climates^48^. Our study indicates that this shift in tree communities could slow those ecosystem processes that are driven by soil food webs in European forests.

## Methods

### Study sites and sampling design

We used a pan-European network of 64 mature, uneven-aged forest plots (30 × 30 m) consisting of three-species mixture stands (34 plots) and corresponding monospecific stands (30 plots, Extended Data Table 1). These plots are part of the FunDivEUROPE exploratory platform^51^, and were established across European forests over 2011-2012 to investigate the role of richness and composition of regionally common and economically important tree species on ecosystem functioning. The studied plots were distributed across four locations featuring different European forest types and spanning a large biogeographic gradient: North Karelia (Finland), Białowieża (Poland), Râşca (Romania), and Colline Metallifere (Italy), corresponding to typical boreal, hemi-boreal, mountainous beech and thermophilous deciduous (Mediterranean) forests, respectively (Supplementary Table 1, Supplementary Fig. 3a). In each location, plots were carefully selected based on tree species richness and composition while minimizing as much as possible covariation with potentially confounding environmental factors such as topography and soil conditions^51^ (Supplementary Fig. 1a). Plot selection was performed so as to include monocultures of all tree species from the local species pool and replicate the three-species mixture treatment with different tree species combinations while maximizing community evenness (Extended Data Table 1). This allowed strict avoidance of a dilution gradient, such as would occur in a design with monoculture stands of only one species combined with mixture stands including this monoculture species, along with a clear distinction between the effects of species diversity and composition. Our stratified plot selection procedure enabled us to mimic formal biodiversity experiments, given that such manipulative approaches are virtually impossible to undertake in mature forests due to the high longevity of tree species. Tree species diversity and composition in the studied plots were predominantly the result of natural community assembly from the regional species pool, combined with local forest management practices. The investigated levels of species richness, *i.e.* one *versus* three tree species, are typical for European forest ecosystems (https://forest.eea.europa.eu/topics/forest-biodiversity-and-ecosystems/forest-ecosystems). Overall, our sampling design encompassed a total pool of 13 tree species, including 12 ectomycorrhizal and one arbuscular mycorrhizal tree species, and the local species pool ranged from three to five tree species per location (Extended Data Table 1, Supplementary Table 1).

### Sampling, extraction and classification of soil organisms

In each plot, we assessed energy fluxes through the soil food web by measuring the biomass of major groups of organisms within this food web^28,52–54^, including both microbial (bacteria and fungi) and faunal (nematodes, microarthropods, and macroinvertebrates) groups. Biomass data for all soil organism groups were expressed per unit surface area (g dry weight m^-2^) at the plot-level. Further details on biomass calculation and the trophic classification of soil organisms are provided in Supplementary Methods.

#### Sampling

Soil organisms were sampled in all plots during the phenological spring of 2017 (Supplementary Table 1), a period of high soil biological activity. We sampled both the litter layer (unfragmented aboveground litter, OL horizon) and the soil layer (including both fragmented/humified organic matter and mineral soil, OF/OH/A horizons). In each plot, we selected five 10 × 10 m subplots, with samples taken equidistantly from three trees of either the same species in monospecific stands or of different species for three-species mixture stands (Supplementary Fig. 3b). For each subplot, soil samples for microbial analyses and nematode extraction were collected by taking five soil cores (10 cm depth, 5.3 cm diameter), spaced approximately 35 cm apart around the equidistant point between the three trees, weighted by tree individual size, *i.e.* individual diameter at breast height. The five cores were gently sieved through 6 mm mesh (to avoid damaging nematodes), homogenized, and pooled at the subplot level for nematode extraction. Pooled soil was then sieved through 2 mm mesh for microbial analyses. All subsamples were stored at 4°C until further processing. For microarthropod extraction, an intact core (10 cm depth, 10 cm diameter), including both the litter and soil layers, was collected from each of three subplots along a southwest-northeast transect and stored at 4°C until further processing. For the hand-sorting of soil macroinvertebrates, an intact monolith (25 cm depth, 25 × 25 cm surface), including both the litter and soil layers, was collected from each of the same three subplots. To express all data per unit surface area, an additional core was sampled, sieved through 2 mm mesh, dried at 105 °C for 48 h, and weighted to measure soil bulk density.

#### Microorganisms

The biomass of bacteria (gram-positive and gram-negative), arbuscular mycorrhizal (AM) fungi, and non-AM fungi was quantified at the plot-level using phospholipid fatty acid (PLFA) data^55^. Fungal community data based on metagenomic amplicon sequencing and bioinformatics^56^ were used to partition non-AM fungal biomass into five trophic guilds: ericoid mycorrhizal fungi, ectomycorrhizal fungi, general saprotrophic fungi, wood saprotrophic fungi, and plant pathogenic fungi (Supplementary Methods). The biomass of each fungal trophic guild was calculated by multiplying its relative abundance, *i.e.* number of reads divided by the total number of reads for the five trophic guilds, by the total non-AM fungal biomass.

#### Nematodes

Nematodes were extracted for each subplot within 72 h after sampling from approximately 100 g of fresh soil using a modified sugar flotation method^57^, before being heat-killed and fixed in 4% formaldehyde. Nematodes were then pooled and counted at the plot-level, and a subsample of approximately 160 randomly selected individuals were identified to family level. The biomass of nematode families was calculated based on body mass data retrieved from the Nemaplex database (http://nemaplex.ucdavis.edu/). Nematode families were assigned to five trophic guilds: herbivores, bacterivores, fungivores, omnivores, and carnivores^58^.

#### Microarthropods

Microarthropods were extracted from the intact core (including both the litter and soil layers) within 72 h after sampling for each subplot using the Berlese-Tullgren funnel method^59^, and were fixed in 70% ethanol. Microarthropods were then counted and identified to species level for collembola and to order level for mites. Microarthropod biomass was estimated based on an allometric model using body length data retrieved from the BETSI database (https://portail.betsi.cnrs.fr/) for collembola, and data from the literature for mites^52^. Microarthropod taxa were assigned to seven trophic guilds^54^ belonging to the following broad trophic groups: microbi-detritivores, fungivores, omnivores, and carnivores.

#### Macroinvertebrates

Macroinvertebrates were hand sorted in the field for each subplot and fixed in 70% ethanol. Macroinvertebrates were then counted and identified to species level for Lumbricidae, Isopoda, Diplopoda, Chilopoda, and Araneae; and order to family level for other taxa^60^. All macroinvertebrate individuals were weighed for body mass. Macroinvertebrates were assigned to 23 trophic guilds^54^ belonging to the following broad trophic groups: herbivores, detritivores, humi-detritivores, omnivores, and carnivores.

### Calculation of metabolic rates and assimilation efficiencies

We used soil microbial respiration and biomass data^61^ to calculate the metabolic rates of microbial groups, while we used a model based on individual body mass, environmental temperature (mean soil temperature during the growing season, see ‘microclimate’ section below), and phylogenetic grouping^62^ to calculate the metabolic rates of faunal groups (Supplementary Methods). Assimilation efficiencies (*i.e.* the proportion of consumed food assimilated by digestion) specific to each food type (plant-derived resource or prey) for faunal consumers were calculated using a model based on food N content^63^, and we also applied a temperature correction of assimilation efficiency^64^ (Supplementary Methods). Assimilation efficiencies of all food resources were set to 1 for microbial consumers, given their external digestion system.

### Plant-derived resources

In each plot, we quantified the biomass of the following plant-derived (basal) resources of the soil food web: living plant fine roots^37^ (absorptive roots belonging the first three root orders) and associated photosynthates/rhizodeposits^33^, plant litter (including dead leaves^61^, dead roots ^65^, and dead wood), and soil organic matter^66^ (SOM, Supplementary Methods). Photosynthates/rhizodeposits refer to organic compounds provided by roots to mycorrhizal fungi, or released directly by roots into the soil. To estimate the biomass of rhizodeposits, we used a mass ratio of 0.4 between net rhizodeposition (the portion of rhizodeposited C remaining in the soil after microbial utilization and respiration) and root biomass, based on the results of a meta-analysis of fixed C partitioning in plant–soil systems^33^. The biomass of rhizodeposits was calculated by multiplying the biomass of living plant fine roots by this factor.

### Reconstruction of the soil food web topology and interaction strengths

We constructed our soil food web from 47 network nodes, including six plant-derived (basal) resources and 41 trophic guilds of soil organisms (consumers) that were differentiated based on multiple traits^54^ (Supplementary Table 2). To establish the topology of the soil food web and quantify trophic interaction strengths (Supplementary Fig. 4), a food web interaction matrix was constructed based on basic food web principles, and *a priori* knowledge of soil organism biology and key traits of consumers following the approach of Potapov (2022)^7^, except that microbes were considered here as consumers rather than basal resources (Supplementary Methods). Briefly, the food web interaction matrix was calculated by multiplying five matrices representing different trait dimensions: phylogenetically-defined feeding preferences, density-dependence, predator-prey interactions related to body mass ratio and hunting strategy, prey defence mechanisms, and spatial niche overlap related to vertical stratification. The five matrices relied on the following assumptions, respectively^7^: (1) there are phylogenetically conserved differences in feeding preferences of consumers^67^; (2) food consumption is density (biomass) dependent, *i.e.* consumers will preferentially feed on food resources that are locally abundant due to a higher encounter rate; (3) the strength of predator-prey interactions is primarily defined by the optimum predator-prey mass ratio (PPMR), *i.e.* a predator is typically larger than its prey, but certain predator hunting traits can modify the optimum PPMR^68^; (4) the strength of predator-prey interactions can be weakened by prey defence traits, *i.e.* prey with efficient physical, chemical, or behavioural protection are consumed less; (5) the strength of trophic interactions between a consumer and its food resource is modulated by the overlap in their spatial niches related to vertical stratification, with greater overlap leading to a stronger interaction. Food-web reconstruction was carried out separately for each plot to account for plot-specific density-dependence.

### Calculation of soil food web energy fluxes, trophic functions and multifunctionality

For the calculation of energy fluxes, we assumed a steady state of the soil food web^6,69^. This means that the energy flowing into a given feeding guild of the food web through food consumption balances the energy lost by excretion, metabolism, and predation of that feeding guild. Energy fluxes to each feeding guild within the food web (kJ m^-2^ day^-1^) were then calculated based on the trophic interaction matrix using the following equation^6,69^:

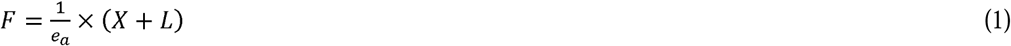

where F is the total flux of energy into the feeding guild, e_a_ is the diet-specific assimilation efficiency, X is the community metabolism of the feeding guild, and L is the energy loss to predation (see Supplementary Methods for further details). To simplify the representation of the food web, we aggregated biomass and energy flux matrices at broader trophic group levels by summing the rows and columns of trophic nodes belonging to the same trophic group (Fig. 1, Supplementary Table 2).

We then calculated five broad trophic functions of the soil food web (Fig. 1): carnivory, the sum of energy fluxes outgoing from fauna to their faunal consumers; microbivory, the sum of energy fluxes outgoing from microbes to their faunal consumers; herbivory, the sum of energy fluxes outgoing from living plant fine roots to their consumers; plant C allocation to soil by living roots, the sum of energy fluxes outgoing from photosynthates and rhizodeposits (via living plant fine roots) to their microbial consumers; and detritivory, the sum of energy fluxes outgoing from detritus (dead organic matter, including plant litter and SOM) to their consumers. Additionally, we calculated eight more specific trophic functions (Fig. 1): bacterivory and fungivory, the sum of energy fluxes outgoing from bacteria and fungi to their faunal consumers, respectively; root pathogenicity and rhizophagy, the sum of energy fluxes outgoing from living plant fine roots to their microbial and faunal consumers, respectively; litter decomposition and litter engineering, the sum of energy fluxes outgoing from plant litter to their microbial and faunal consumers, respectively; and SOM decomposition and soil engineering, the sum of energy fluxes outgoing from SOM to their microbial and faunal consumers, respectively. Decomposition refers to the assimilation and mineralization of dead organic matter through respiration, a process which is mediated mainly by microbes in soil^52,70^. Engineering refers to the physical modification, maintenance, or creation of habitats^2^, which is a major way through which soil faunal detritivory affects the decomposition, transformation, and translocation of dead organic matter^7,71,72^. Such inferences from energy fluxes about effects that are not purely trophic are especially justifiable in soil where habitat and food resources are tightly interlinked^7^. As such, the faunal consumption of litter results in the conversion of litter into faeces which in turn accelerates its decomposition through chemical and physical changes^73,74^, as a form of litter engineering. Similarly, the faunal consumption of SOM is linked to biopedturbation and soil structure maintenance^74^ in a manner that represents soil engineering.

We quantified soil food web multifunctionality based on ten trophic functions: plant C allocation to soil by living roots, root pathogenicity, rhizophagy, litter decomposition, litter engineering, SOM decomposition, soil engineering, bacterivory, fungivory, and carnivory. Defining high multifunctionality by fast rates of multiple functions simultaneously^75^, we calculated soil food web multifunctionality as the average of range-standardized values of each of the ten trophic functions (Supplementary Methods). This ‘averaging’ approach provides a straightforward and easily interpretable measure of multifunctionality but also has some limitations, such as its inability to distinguish between multiple functions performing at intermediate values from some performing at high values while others perform at low values due to trade-offs among functions^76^. To check that our inferences were robust to multiple alternative approaches, we also calculated soil food web multifunctionality using the ‘threshold’ approach^76^ by estimating the number of trophic functions whose value exceeded 10, 20, 30, 40, 50, 60, 70, 80, and 90% of the maximal value (Supplementary Methods). We found that the two methods provided fairly consistent results for the effects of tree diversity and composition with both the taxonomic and functional approaches (Supplementary Table 3). Additionally, we adopted the ‘single functions’ approach to help illuminate which individual functions drive trends in the effects of tree diversity and composition on soil food web multifunctionality^76^.

### Tree functional (trait-based) diversity and composition

Functional diversity and composition of tree communities were characterized using a set of nine plant functional traits known to be directly involved in resource economics^9,31^. These traits included three leaf traits: leaf nitrogen content (LNC), specific leaf area (SLA), and leaf dry matter content (LDMC); and six fine (absorptive) root traits^65^: root nitrogen content (RNC), root tissue density (RTD), specific root length (SRL), mean root diameter (D_m_), root length density (RLD), and ectomycorrhizal colonisation intensity (%Myc). Trait data of each tree species was mostly derived from plot-specific measurement in the field. See Supplementary Methods for further details and Supplementary Table 4 for trait values of each tree species and geographic location.

To quantify functional diversity, we calculated the functional dispersion (FDis) index corresponding to the mean distance of each species to the centroid of all species within the multidimensional trait space^77^. For functional composition, we calculated community-weighted means (CWMs) of each trait. Both FDis and CWM values were computed based on the relative basal area of tree species. We performed a Principal Component Analysis (PCA) on all CWM traits, which simplified functional composition into two dimensions^65,78^ (Extended data Fig. 3a, Supplementary Table 5): (1) a leaf economics spectrum (LES, 45.2 % of variation), which ranged from slow/conservative leaf attributes (N-poor and dry matter-rich leaves with low leaf area per unit mass) to fast/acquisitive leaf attributes (N-rich and dry matter-poor leaves with high leaf area per unit mass), and also aligned here with a fine-root gradient of belowground resource foraging strategies ranging from low foraging efficiency (thick fine roots with low length per unit mass) to high foraging efficiency (thin fine roots with high length per unit mass); (2) a fine-root economics spectrum (RES, 31.5 % of variation), which ranged from slow/conservative fine-root attributes (N-poor fine roots with low tissue density) to fast/acquisitive fine-root attributes (N-rich fine roots with high tissue density), and also aligned here with a fine-root gradient of soil exploration strategies ranging from ‘do-it-yourself’ attributes (high root length density and low ectomycorrhizal colonisation intensity) to ‘outsourcing’ attributes (low root length density and high ectomycorrhizal colonisation intensity). The mean values of FDis and the scores of the two first PCA axes of CWM trait ordination for each location are shown in Extended Table 1.

### Tree and understorey vegetation

We characterized forest vegetation using a set of eight variables measured in each plot: tree aboveground biomass (AGB), tree leaf area index, tree aboveground litterfall, total root biomass, aboveground biomass of both woody and herbaceous understorey plants, and aboveground C:N ratio and species richness of understorey plant communities (Supplementary Methods).

### Environmental drivers

#### Abiotic conditions

We characterized abiotic conditions in each plot using a set of six variables related to soil texture, macroclimate, and topography: the sand, silt, and clay content of mineral soil (A horizon, 10 cm depth), mean annual temperature and precipitation, and altitude. We performed a PCA to reduce the dimensionality of abiotic properties into two dimensions (Supplementary Fig. 1a): (1) a soil texture and macroclimate gradient ranging from coarse soil texture, and dry and cold macroclimate to fine soil texture, and wet and hot macroclimate (70.8 % of variation); (2) a temperature gradient ranging from cold macroclimate with high elevation to warm macroclimate with low elevation (19.1 % of variation).

#### Microclimate

We characterized microclimate using a set of six variables: soil temperature and moisture (measured at 8 cm soil depth), and air temperature (measured 50 cm above the ground) for both annual and growing season (daily mean temperatures > 5 °C) periods (Supplementary Methods). We performed a PCA to simplify microclimatic properties into a single dimension ranging from cold and wet to hot and dry (Extended data Fig. 3b, 63.1 % of variation).

#### Leaf litter quality

We characterized freshly fallen tree leaf litter quality using a set of 16 chemical and physical variables mainly related to elemental composition and carbon quality^79^: the concentrations of N, P, Ca, Mg, and K, and the C:N and C:P ratios, the proportion of carbon fractions (lignin, cellulose, hemicellulose, and water-soluble compounds), the concentrations of secondary metabolites (condensed tannins, total phenolics, and soluble phenolics), as well as the litter pH and water holding capacity. Leaf litter quality data of each tree species was derived from location-specific measurement (Supplementary Methods).

We then quantified the functional diversity and composition of tree leaf litter by calculating the FDis index and CWMs of each litter property based on the relative leaf litter mass of the component tree species. We performed a PCA to simplify tree leaf litter properties into one dimension ranging from low to high nutritional quality (Extended data Fig. 3c, 42.5 % of variation).

#### Soil fertility

We characterized soil fertility using a set of six variables measured in each plot^66^: the organic C content of mineral soil (A horizon, 10 cm depth), the mass of the forest floor (including unfragmented aboveground litter and fragmented/humified organic matter, OL/OF/OH horizons), as well as the pH and C:N ratios of both the forest floor and mineral soil (Supplementary Methods). We performed a PCA to simplify soil properties into a single dimension ranging from low to high fertility (Supplementary Fig. 1b, 40.1 % of variation).

### Measured ecosystem processes

To characterize *in situ* patterns of litter decomposition, we used field-based data of naturally occurring leaf litter decomposition measured in each plot^80^ (Supplementary Methods). To characterize SOM decomposition, we further used soil microbial respiration and biomass data^61^ (Supplementary Methods). To characterize soil and litter engineering, we used humus type and forest floor mass data^66^ as these two variables reflect the transformation and translocation of dead organic matter by faunal detritivory^81^.

### Statistical analyses

All analyses were performed using R v4.3.0^82^. To test how the diversity and composition of tree communities affect soil food web properties, we used Bayesian multi-level models accounting for the hierarchical design of the study, where plots were nested within four geographic locations across Europe. To ensure that tree diversity and compositional effects were not biased by confounding factors, we also explicitly included abiotic covariates into the models (first two PCA axes of abiotic conditions, Supplementary Fig. 1a), thereby statistically controlling for variation in abiotic conditions both among and within locations. We adopted both taxonomic and functional approaches. For the taxonomic approach, we used linear mixed-effect models (LMMs) including tree species richness (comparing three-species mixture stands to corresponding monospecific stands) and abiotic covariates (see above) as fixed factors, and tree species composition (a factor listing all tree species present, Extended Data Table 1) and location as random factors. For the functional approach, we used LMMs including tree functional diversity (FDis), tree functional composition (first two PCA axes of tree trait CWMs, Extended data Fig. 3a), and abiotic covariates (see above) as fixed factors, and location as a random factor. To further investigate the effects of functional composition, we also used LMMs including individual tree CWM trait and abiotic covariates as fixed factors and location as a random factor. Bayesian LMMs were fitted with Markov Chain Monte Carlo (MCMC) methods, using non-informative priors for all parameters (Supplementary Methods). We used random intercept models (varying intercepts but a common slope across locations) to explore general effects across European forests (Supplementary Methods).

For each model, we performed variance partitioning to quantify the proportion of variance in the dependent variable explained by three groups of variables: diversity, composition, and biogeography (including abiotic covariates and location)^83^. To minimize issues related to correlation between predictors within each group (especially between abiotic covariates and location for biogeography, Supplementary Fig. 1a), we performed variance partitioning at the level of these variable groups by summing over the variance terms of the covariates included in each group^83^. This allowed to incorporate the covariances between group covariates to the variance term of that group, thereby reducing the covariance between variance components at variable group level. The means of posterior distributions of variance components were standardized by the sum of all variance components, including the residuals. Our Bayesian modelling approach allowed to avoid singular fit issues, which are common when dealing with complex models with few random-factor levels, and can bias variance partitioning^83,84^. We calculated the effect size of fixed factors as the slope regression coefficient, standardized by the standard deviation (Supplementary Methods). We quantified the uncertainty of effect size based on credible intervals^85^. Standardized slope (β_st_) values of |β_st_| < 0.12, 0.12 ≤ |β_st_| < 0.24, 0.24 ≤ |β_st_| < 0.41, and |β_st_| ≥ 0.41 were interpreted as neutral, weak, moderate, and strong effects, respectively^86^. We used *p*-values of slope regression coefficients and Bayes factors as measures of evidence for fixed and random effects, respectively. The *p*-values were derived from posterior distributions using a two-tailed test^87^. Bayes factors were used to quantify the support for models including the random factor tested, compared to null models without the random factor. Bayes factors were calculated as the ratio of marginal likelihoods of the two models (Supplementary Methods).

To test *a priori* multivariate causal hypotheses regarding how tree diversity and composition effects on soil food web multifunctionality are mediated by changes in tree and understorey plant properties and microenvironment, we performed piecewise multi-level structural equation modelling (SEM). We first constructed a causal diagram based on ecological theory and our prior knowledge of the system (Extended data Fig. 4) and built an initial full SEM model (Supplementary Table 6). Following a local estimation approach^88^, we fitted component LMMs with random intercept across locations using Bayesian methods similar to those described above. We performed model selection using information_theoretic methods to select the most parsimonious SEM model. For each component LMM within the initial full SEM model, stepwise backward selection was performed using Pareto smoothed importance-sampling leave-one-out (PSIS-LOO) cross-validation^89^ (Supplementary Methods). To test the goodness_of_fit of the selected SEM model, we used the Shipley’s d_sep test^90^ (Supplementary Methods). To quantify and compare the magnitude of tree diversity and compositional effects on soil food web multifunctionality, indirect effects were calculated as the product of the standardized coefficients along each path. Total effects were calculated as the sum of indirect effects.

## Supporting information

Tree biodiversity effects on soil food web multifunctionality - Supplementary Information

## Code availability

Code for computation of food-web energy fluxes and statistical analyses will be made available upon acceptance for publication.

## Data availability

All data generated or analysed during this study will be made available upon acceptance for publication.

## Acknowledgements

This research was part of the SoilForEUROPE project funded through the 2015-2016 BiodivERsA COFUND call for research proposals, with the national funders Agence Nationale de la Recherche (ANR, France), Belgian Science Policy Office (BELSPO, Belgium), Deutsche Forschungsgemeinschaft (DFG, Germany), Research Foundation Flanders (FWO, Belgium), and the Swedish Research Council (FORMAS, Sweden). We greatly thank Jean-François David for his help in macroinvertebrate identification, and Jurgen van Hal for his help in microarthropod identification. We further thank the site managers Leena Finér, (Natural Resource Institute Finland, LUKE), Bogdan Jaroszewicz (University of Warsaw, Poland), Olivier Bouriaud (Forest Research and Management Institute, ICAS, Romania), Filippo Bussotti and Federico Selvi (University of Florence, Italy), Jakub Zaremba, Ewa Chećko, Iulian Dănilă, and Timo Domisch for their assistance with field sampling. Silhouette images used in Fig. 1 are in the public domain under the Creative Commons Attribution License ‘CC BY’ (source: https://www.phylopic.org/). We thank all authors of these images. The study was supported by the TRY initiative on plant traits (https://www.try-db.org/). The TRY database is hosted at the Max Planck Institute for Biogeochemistry (MPI BGC, Germany) and supported by Future Earth and the German Centre for Integrative Biodiversity Research (iDiv) Halle-Jena-Leipzig. We thank all scientists who contributed leaf trait data to the public domain under the ‘CC BY’ license, which was used in this study.

## Author Contributions

L.H., P.K., D.A.W., and M.B. conceived the ideas and designed methodology. S.H., along with D.A.W., M.B., J.B., F.B., S.C., T.D., N.F., A.M., B.M., J.N., K.V., and M.S.-L. acquired the funding and performed the plot selection and sampling design. P.K., S.H., F.B., J.B., F.B., S.C., J.-F.D., T.D., N.F., L.M.G., A.M., J.N., L.D.P.-S., M.S.-L., and J.W. performed the sampling. Most authors performed analyses for different subsets of the data: L.D.P.-S., K.G., and F.B. for microorganisms; L.H. and P.K. for nematodes; M.B. and L.H. for microarthropods; P.G., J.N., J.-F.D., and T.D. for macroinvertebrates; J.W., J.B., and M.S.-L. for roots; L.M.G., N.F., and S.H. for litter and soil respiration; M.R. for microclimate; K.V. for understorey plants; B.M. for soil texture. L.H. performed the data compilation, food web modelling, and statistical analyses. L.H. led the writing of the manuscript, and P.K., D.A.W., M.B., and S.H. contributed critically to the drafts. All authors revised the manuscript and gave final approval for publication.

---

**Extended Data Fig. 1.**
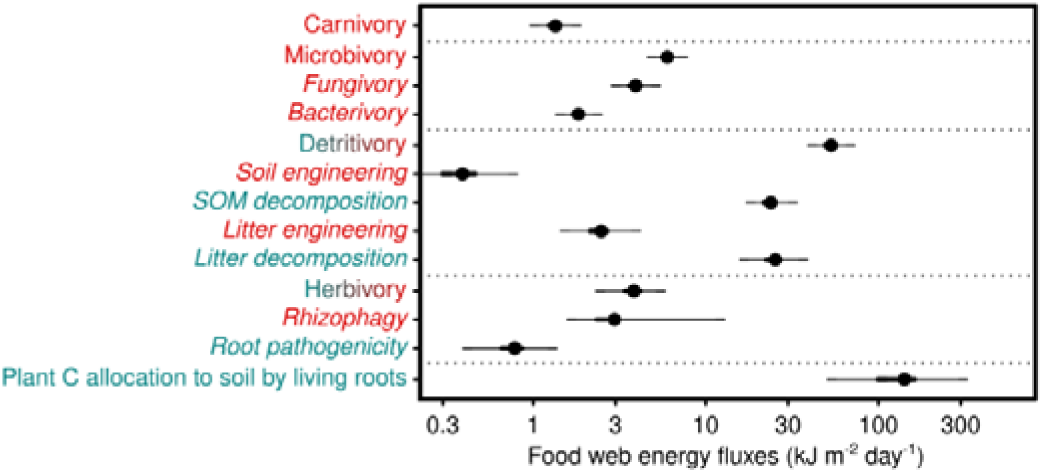
Soil food web trophic functions (aggregated energy fluxes per function) predicted by the model for average conditions (mean values for each predictor). Circles represent the mean values, with thick and thin error bars representing 50% and 95% credible intervals, respectively. The *x* axis is displayed on a logarithmic scale. Predicted values were derived from Bayesian linear mixed-effect models used for the functional approach (Methods).

**Extended Data Fig. 2.**
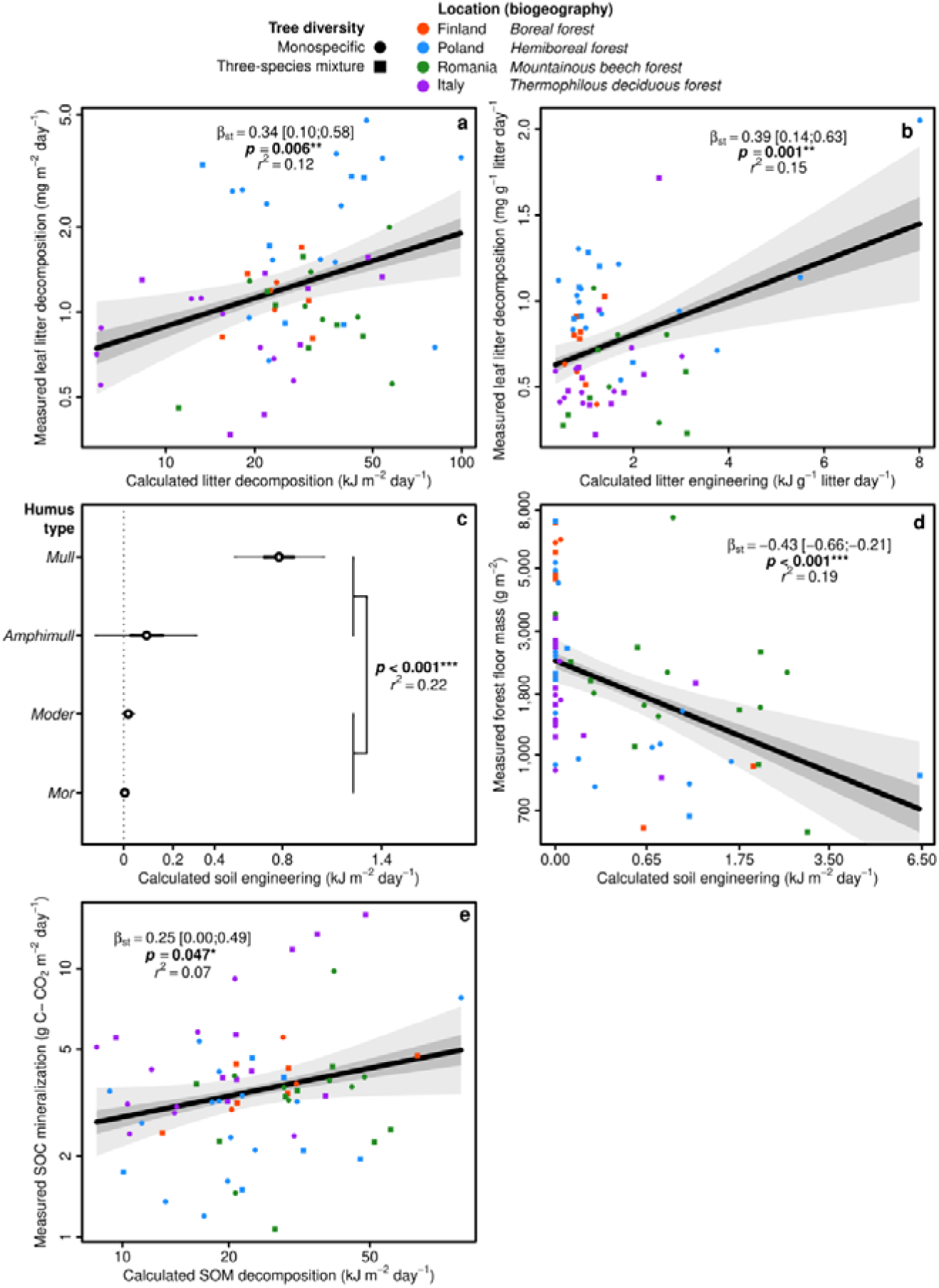
Relationships between calculated soil food web trophic functions and measured soil functions. Relationships were tested using Bayesian linear regression models. For all panels, calculated soil food web trophic functions and measured ecosystem processes or properties are displayed on the *x* and *y* axes, respectively. Thick and thin error bars, along with dark and light grey filled area, represent 50% and 95% credible intervals, respectively. β_st_ is the slope regression coefficient, standardized by standard deviation (effect size). Values in square brackets are the 95% credible intervals of the standardized slopes. For panels a, c, d, and e, the *x* and *y* axes are displayed on a logarithmic scale. *, *p* < 0.050; **, *p* < 0.010; ***, *p* < 0.001.

**Extended Data Fig. 3.**
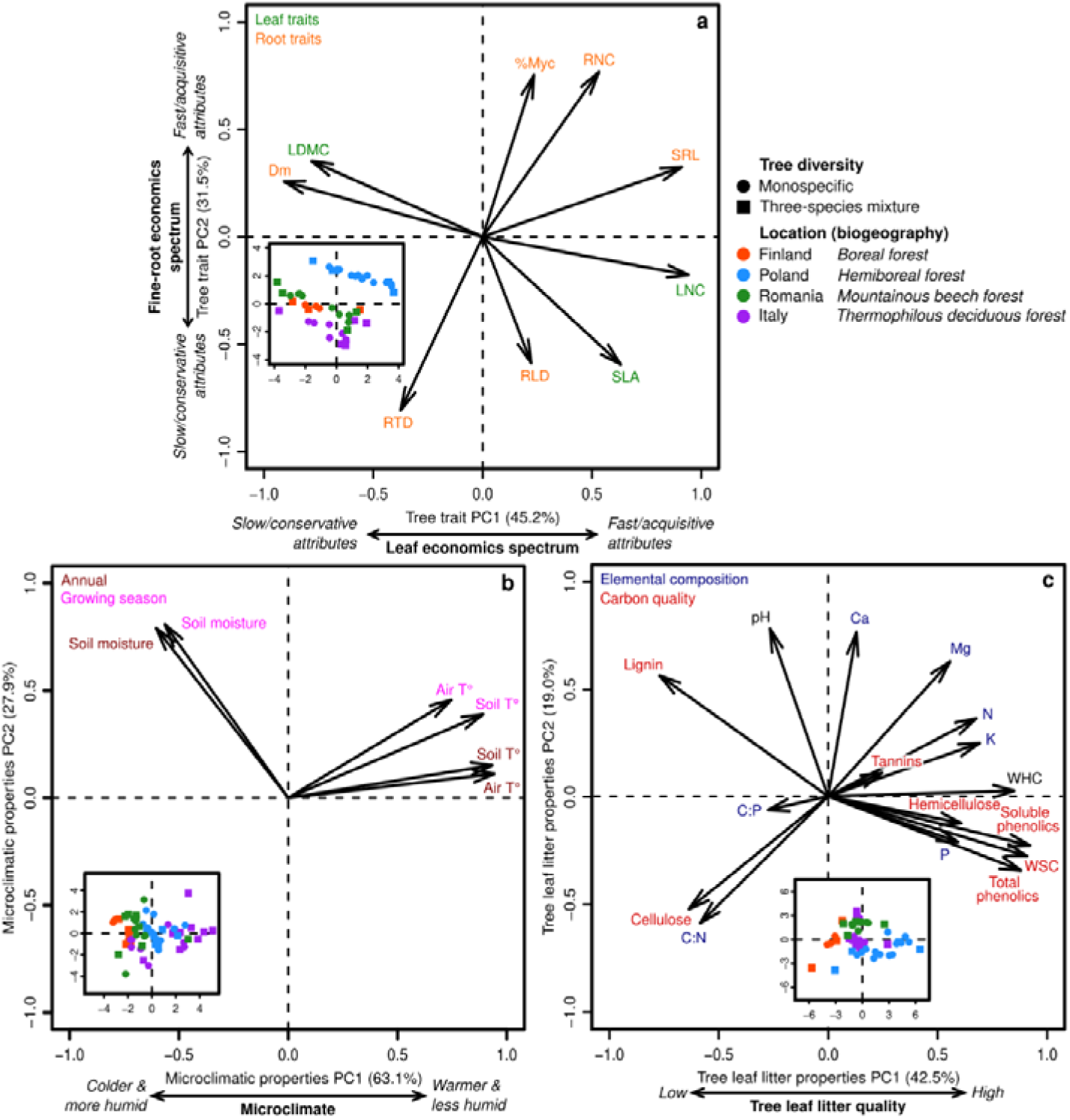
Principal Component Analysis (PCA) of tree functional composition based on community-level tree traits (a), microclimatic properties (b), and community-level tree leaf litter properties (c). The values for community-level tree traits and leaf litter properties are community-weighted means (CWMs), weighted by relative basal area for tree traits and by leaf litterfall mass per tree species for leaf litter properties. Variable loadings represent the correlations of variables with corresponding principal components. Insets show plot coordinates. Mean score values for the two first PCA axes of tree functional composition per species composition for each location are shown in Extended Data Table 1. Correlation and contribution of tree CWM traits to the principal components (panel **a**) are shown in Supplementary Table 6. LNC, leaf nitrogen content; SLA, specific leaf area; LDMC, leaf dry matter content; RNC, root nitrogen content; SRL, specific root length; RTD, root tissue density; Dm, mean root diameter; RLD, root length density; %Myc, ectomycorrhizal colonisation intensity; WSC, water soluble compounds; WHC, water holding capacity. Tannins are condensed tannins.

**Extended Data Fig. 4.**
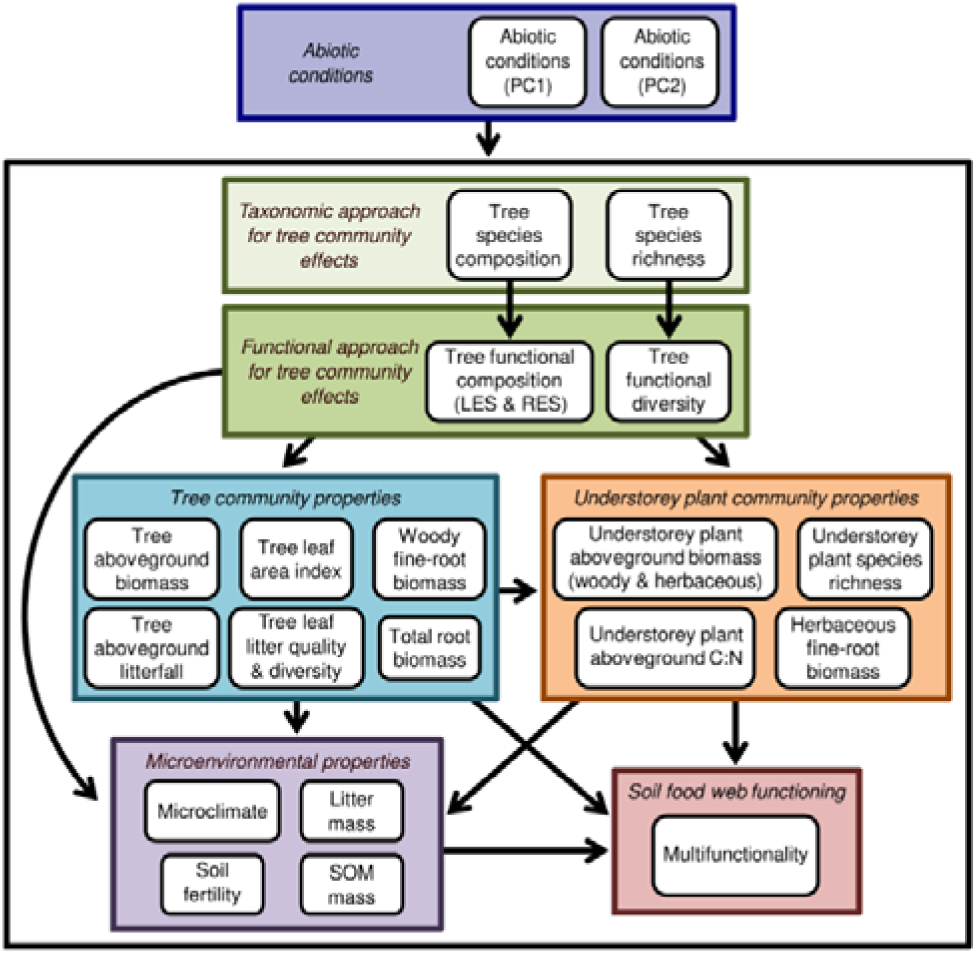
Conceptual causal model of the *a priori* hypotheses about how tree diversity and composition (in green) drives soil food web multifunctionality (red) through changes in tree community properties (blue), understorey plant community properties (orange), and micro-environmental properties (purple). Arrows represent the flow of causality. LES, leaf economics spectrum; RES, fine-root economics spectrum; SOM, soil organic matter.

**Extended Data Fig. 5.**
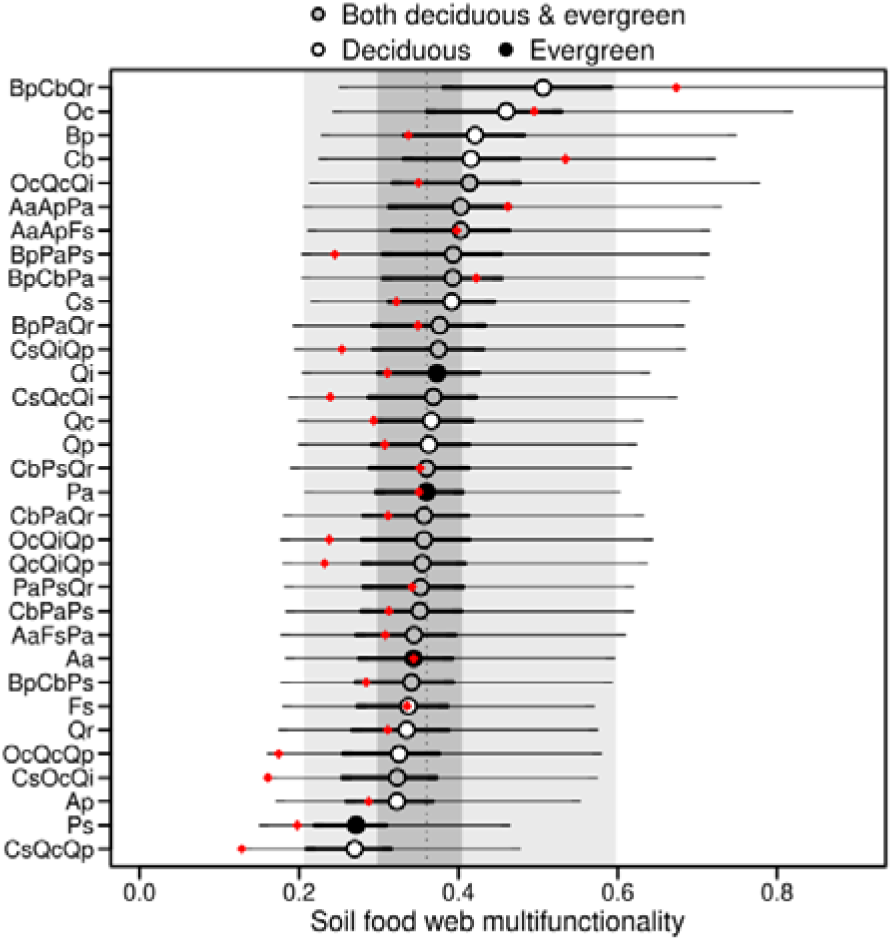
Random effects of tree species composition on soil food web multifunctionality. Circles represent the mean of predicted values derived from Bayesian linear mixed-effect models used for the taxonomic approach (Methods). The dotted vertical line represents the general intercept of the model. Thick and thin error bars, along with dark and light grey filled area, represent 50% and 95% credible intervals, respectively. Red diamonds represent the mean observed values of soil food web multifunctionality for each tree species composition. Aa, *Abies alba*; Ap, *Acer pseudoplatanus*; Bp, *Betula pendula*/*pubescens*, Cb, *Carpinus betulus*, Cs, *Castanea sativa*; Fs, *Fagus sylvatica*; Oc, *Ostrya carpinifolia*; Pa, *Picea abies*; Ps, *Pinus sylvestris*; Qc, *Quercus cerris*; Qi, *Q. ilex*; Qp, *Q. petraea*; Qr, *Q. robur*.

**Extended Data Fig. 6.**
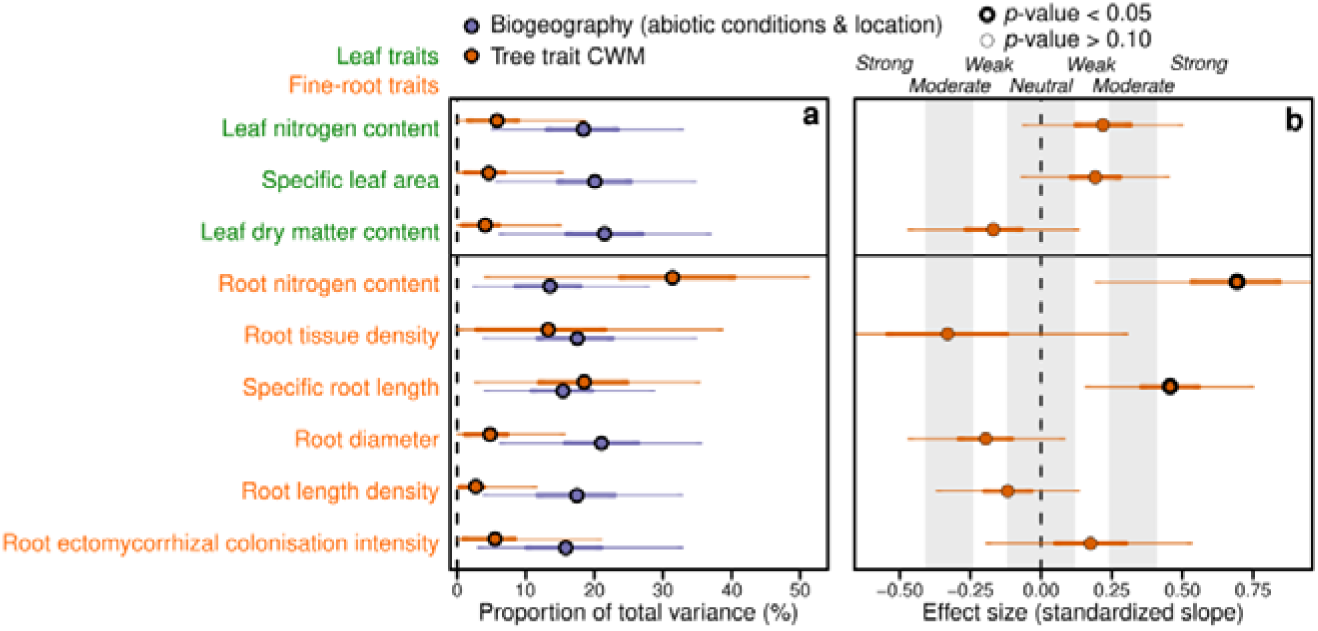
Community-weighted means of tree traits as drivers of soil food web multifunctionality. **a**, Decomposition of the total variance explained by community-weighed means of each tree trait, and biogeography (abiotic conditions and location), as determined by variance partitioning. **b**, Effect sizes calculated as slope regression coefficients standardized by standard deviation. Circles represent the mean effect sizes, with thick and thin error bars representing 50% and 95% credible intervals, respectively.

**Extended Data Table 1.**
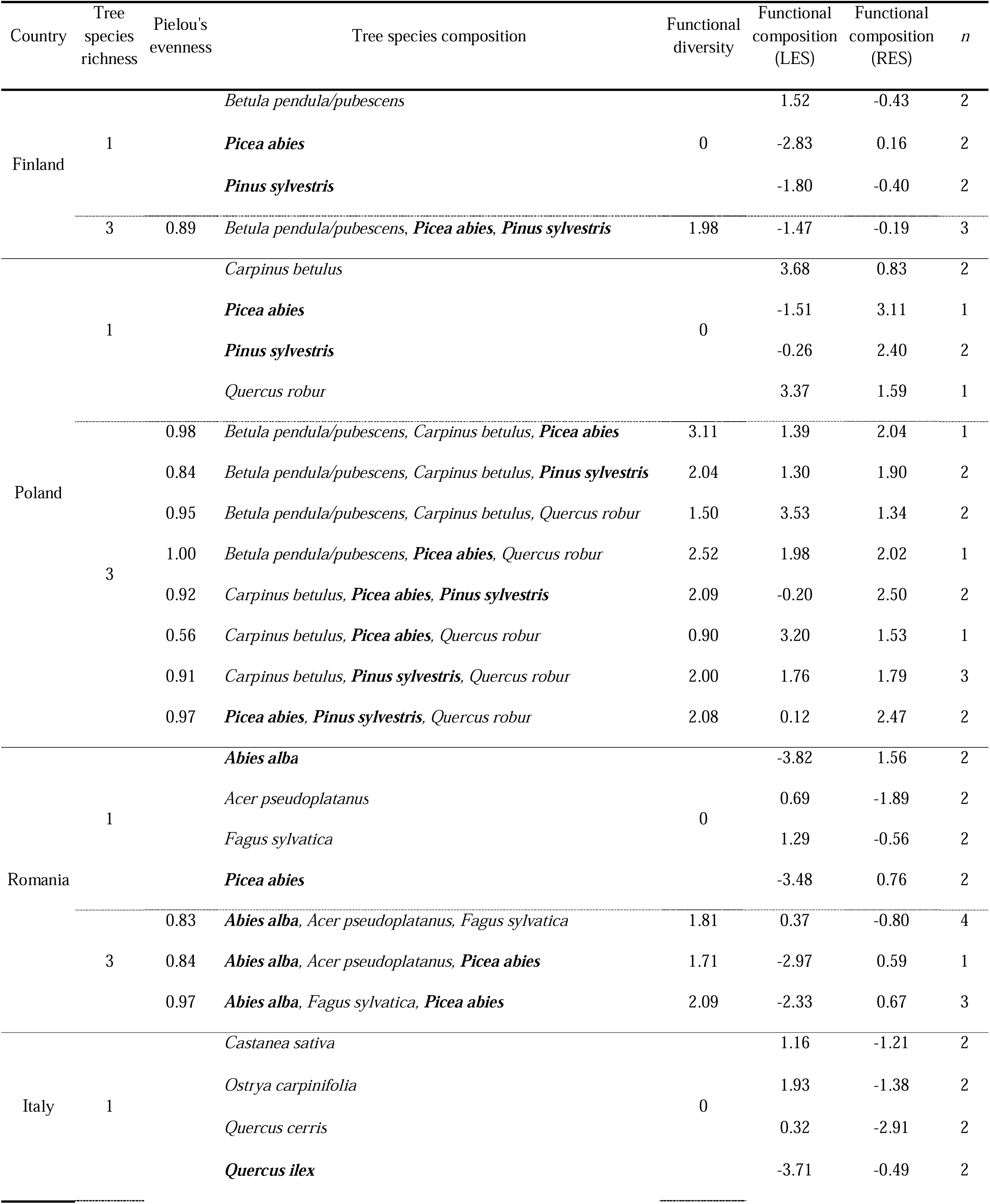
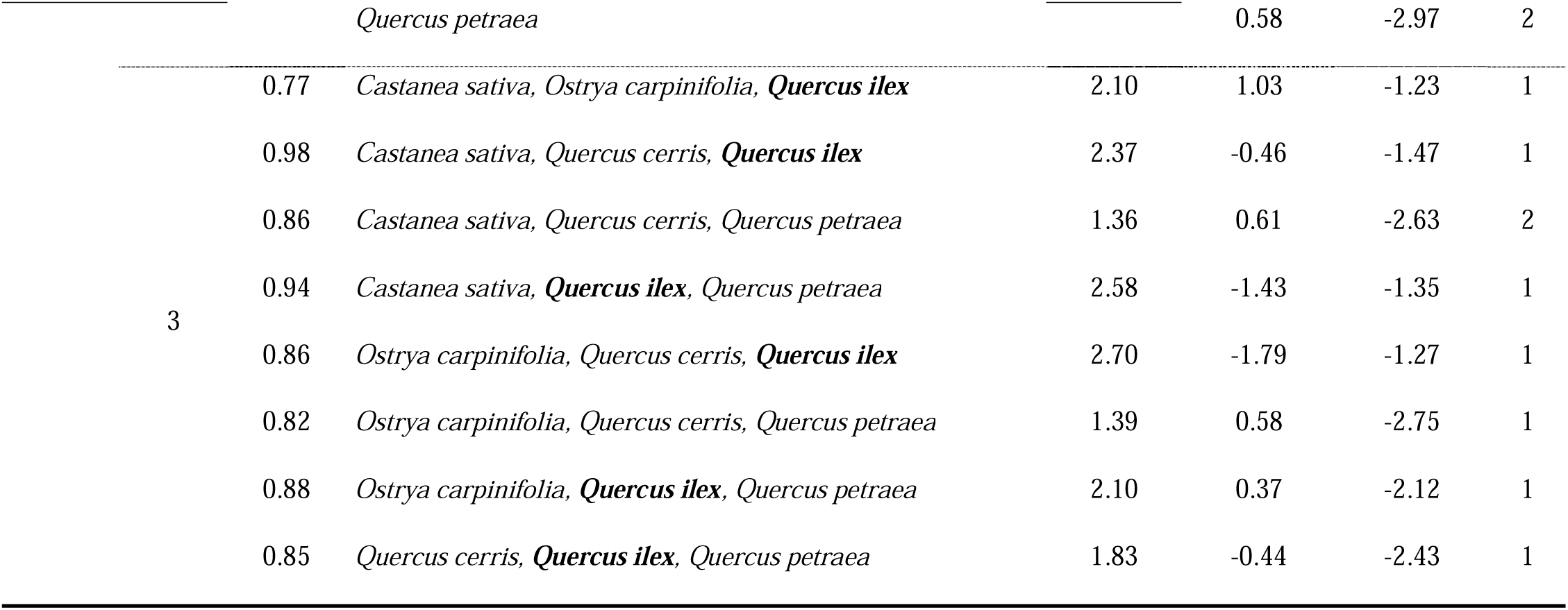
Sampling design with the species composition and diversity of tree communities. Mean values are provided when applicable. Evergreen species are shown in bold.

**Extended Data Table 2.**
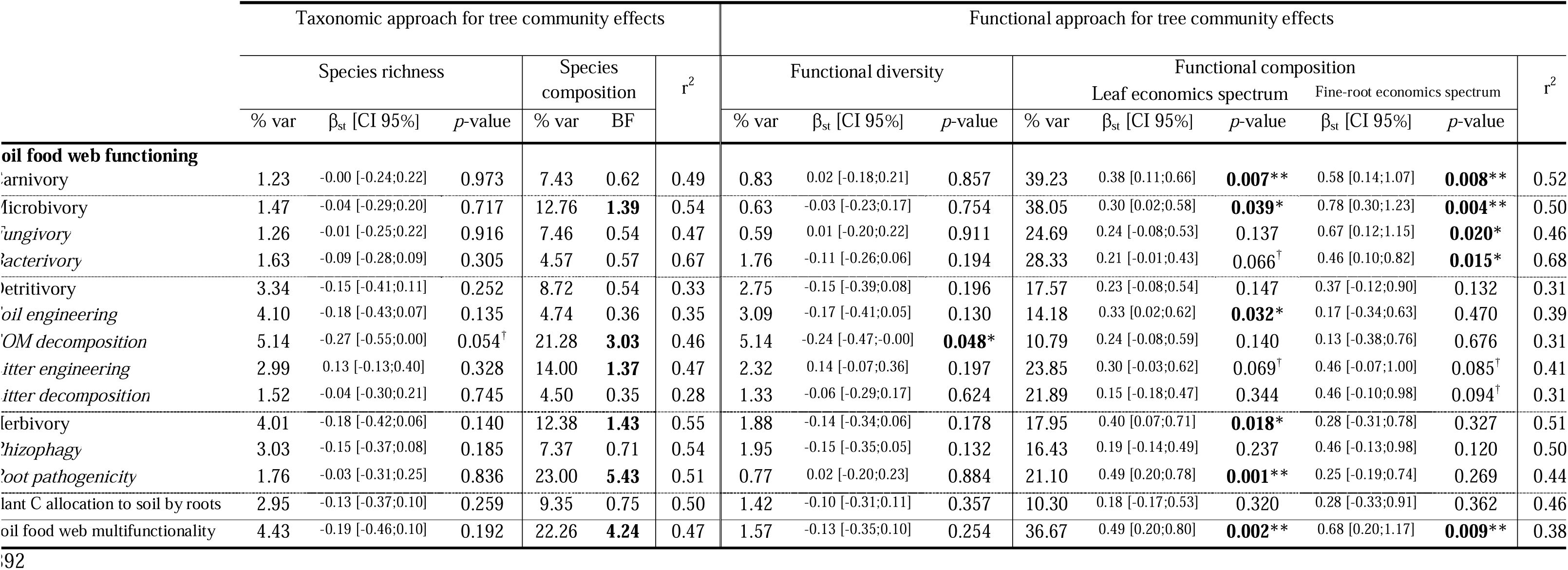
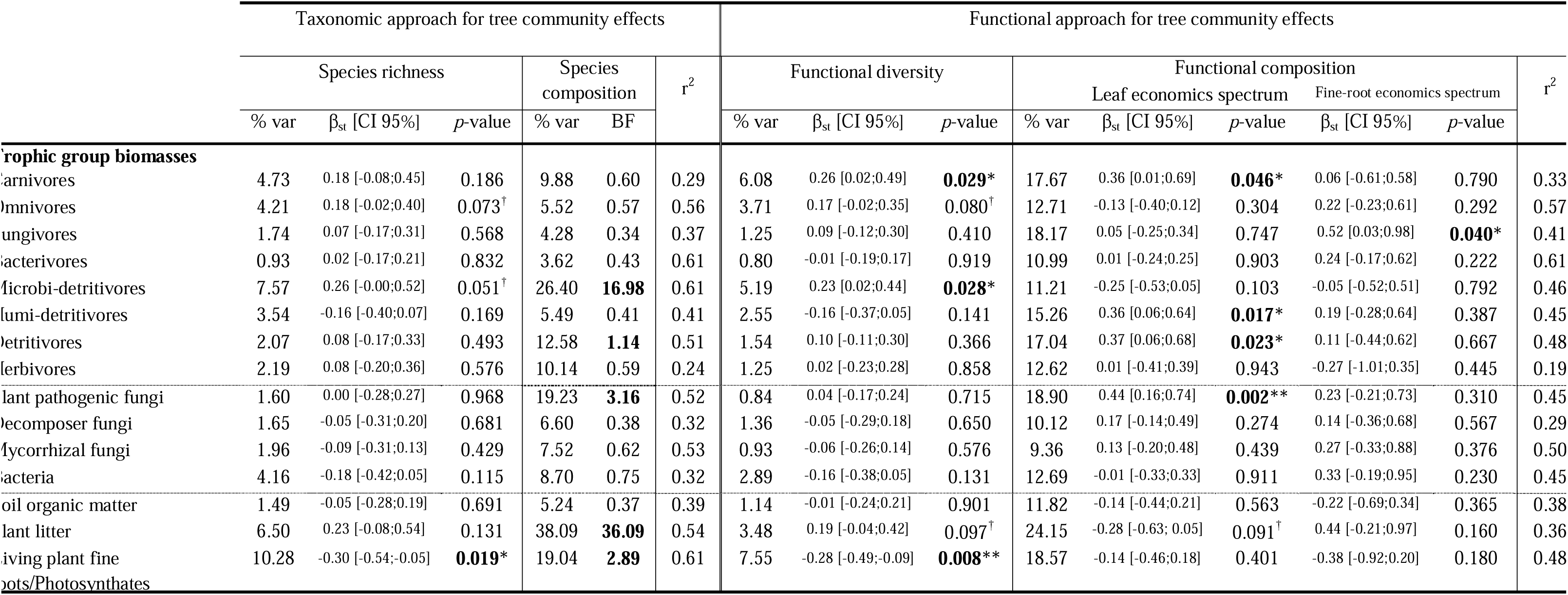
Results of Bayesian multi-level models testing tree community effects on trophic functions and trophic group biomasses of the soil food web. Both the leaf and fine-root economics spectra (LES & RES) variables representing tree functional composition ranged from slow/conservative to fast/acquisitive attributes (Extended Data Fig. 3a). % var, proportion of total variance. β_st_, slope regression coefficients standardised by standard deviation. CI 95%, 95% credible intervals. BF Bayes factor (Methods). Significant effects (*p* < 0.05 or BF > 1) are reported in bold. ^†^, *p* < 0.10; *, *p* < 0.050; **, *p* < 0.010; ***, *p* < 0.001. SOM, soil organic matter.

**Extended Data Table 3.** Results of the best-supported Bayesian multi-level SEM model testing the mediation of tree community effects on soil food web multifunctionality. β_st_, (partial) regression slope coefficients, standardized by standard deviation. CI 95%, 95% credible intervals. % of r^2^_m_, proportion of marginal r^2^ explained by the predictor variable. r^2^_m_ and r^2^, marginal and conditional r^2^, that are the proportion of variance explained by fixed factors only, and by both fixed factors and random factors, respectively. For LES and RES response variables however, r^2^_m_ and r^2^_c_ represent the proportion of variance explained by tree species composition only, and both tree species composition and location, respectively. VIF, variance inflation factor. ‘Tree species composition’ predictor variable was fitted as a random factor. LES, leaf economics spectrum. RES, fine-root economics spectrum. Significant effects (*p* < 0.05) are reported in bold. ^†^, *p* < 0.10; *, *p* < 0.050; **, *p* < 0.010; ***, *p* < 0.001; ‡, log(*x*)-transformed; L, log(*x*+1)-transformed.

| Response variable | Predictor variable | $\beta_{st}$ [CI 95%] | $p$-value | % of $r^2_m$ | $r^2_m$ | $r^2_c$ | VIF |
| --- | --- | --- | --- | --- | --- | --- | --- |
| Tree functional diversity | Tree species richness | 0.95 [0.87;1.03] | **<0.001\*\*\*** |  | 0.90 | 0.90 |  |
| Tree functional composition | Tree species composition |  |  |  | 0.79 | 0.97 |  |
| Tree functional composition | Tree species composition |  |  |  | 0.22 | 0.99 |  |
| Tree aboveground biomass$^\ddagger$ | Tree functional diversity | 0.16 [0.00;0.33] | **0.048\*** | 27.9 | 0.13 | 0.62 | 1.00 |
|  | Tree functional composition (RES) | 0.25 [-0.08;0.57] | 0.134 | 72.3 |  |  | 1.00 |
| Tree aboveground litterfall$^\ddagger$ | Tree functional diversity | 0.11 [-0.04;0.27] | 0.128 | 19.4 | 0.10 | 0.72 | 1.07 |
| | Tree aboveground biomass$^\ddagger$ | 0.27 [-0.04;0.50] | **0.022\*** | 80.6 | | | 1.07 |
| Tree leaf litter quality (PC1) | Tree functional composition (LES) | 0.55 [0.37;0.72] | **<0.001\*\*\*** |  | 0.36 | 0.60 |  |
| Total root biomass$^\ddagger$ | Tree functional diversity | -0.21 [-0.42;-0.00] | **0.050\*** | 29.2 | 0.23 | 0.32 | 1.03 |
| | Tree aboveground biomass$^\ddagger$ | -0.34 [-0.63;-0.02] | **0.033\*** | 70.8 | | | 1.03 |
| Microclimate (PC1) | Tree functional diversity | -0.22 [-0.40;-0.04] | **0.019\*** | 3.6 |  |  | 1.02 |
| | Tree functional composition (LES) | 0.25 [0.00;0.49] | 0.051$^\dagger$ | 4.7 | | | 1.56 |
|  | Tree functional composition (RES) | 0.60 [0.17;0.97] | **0.011\*** | 23.7 | 0.54 | 0.61 | 1.74 |
|  | Abiotic conditions (PC1) | 0.67 [0.23;1.14] | **0.007\*\*** | 28.3 |  |  | 1.27 |
|  | Abiotic conditions (PC2) | 0.79 [0.31;1.29] | **0.008\*\*** | 39.7 |  |  | 1.16 |
| Understorey plant aboveground biomass (herbaceous)□ | Tree aboveground biomass$^\ddagger$ | -0.44 [-0.70;-0.18] | **<0.001\*\*\*** | 76.1 | 0.14 | 0.62 | 1.07 |
| | Total root biomass$^\ddagger$ | -0.23 [-0.44;-0.02] | **0.029\*** | 24.2 | | | 1.07 |
| Soil food web multifunctionality$^\ddagger$ | Tree aboveground biomass$^\ddagger$ | 0.40 [0.15;0.66] | **0.003\*\*** | 19.5 | | | 1.34 |
| | Tree aboveground litterfall$^\ddagger$ | -0.31 [-0.56;-0.06] | **0.019\*** | 12.3 | | | 1.12 |
|  | Tree leaf litter quality (PC1) | 0.44 [0.22;0.66] | **<0.001\*\*\*** | 22.8 |  |  | 1.04 |
| | Total root biomass$^\ddagger$ | 0.28 [0.05;0.51] | **0.017\*** | 10.0 | 0.50 | 0.52 | 1.36 |
| | Understorey plant aboveground biomass (herbaceous)□ | 0.37 [0.14;0.60] | **0.002\*\*** | 16.7 | | | 1.21 |
|  | Microclimate (PC1) | 0.39 [0.16;0.63] | **0.001\*\*** | 18.6 |  |  | 1.13 |
| ~~ Tree species richness | ~~Tree aboveground litterfall$^\ddagger$ | 0.08 | 0.520 | | | | |
| ~~Tree functional composition (LES) | ~~Tree functional composition (RES) | -0.80 | **<0.001\*\*\*** |  |  |  |  |
| ~~Tree aboveground litterfall$^\ddagger$ | ~~Tree leaf litter quality (PC1) | -0.35 | **0.005\*\*** | | | | |
| ~~Tree aboveground biomass$^\ddagger$ | ~~Tree leaf litter quality (PC1) | -0.29 | **0.021\*** | | | | |

## References

1. Cebrian, J. Patterns in the fate of production in plant communities. Am. Nat. 154, 449–468 (1999).

2. Jones, C. G., Lawton, J. H. & Shachak, M. Positive and negative effects of organisms as physical ecosystem engineers. Ecology 78, 1946–1957 (1997).

3. Cardinale, B. J. et al. The functional role of producer diversity in ecosystems. Am. J. Bot. 98, 572–592 (2011).

4. Eisenhauer, N. et al. Chapter One - A multitrophic perspective on biodiversity–ecosystem functioning research. in Advances in Ecological Research (eds. Eisenhauer, N., Bohan, D. A. & Dumbrell, A. J.) vol. 61 1– 54 (Academic Press, 2019).

5. Wardle, D. A. et al. Ecological linkages between aboveground and belowground biota. Science 304, 1629– 1633 (2004).

6. Barnes, A. D. et al. Energy Flux: The Link between Multitrophic Biodiversity and Ecosystem Functioning. Trends Ecol. Evol. 33, 186–197 (2018).

7. Potapov, A. M. Multifunctionality of belowground food webs: resource, size and spatial energy channels. Biol. Rev. 97, 1691–1711 (2022).

8. Brown, J. H., Gillooly, J. F., Allen, A. P., Savage, V. M. & West, G. B. Toward a metabolic theory of ecology. Ecology 85, 1771–1789 (2004).

9. Reich, P. B. The world-wide ‘fast–slow’ plant economics spectrum: a traits manifesto. J. Ecol. 102, 275–301 (2014).

10. Hooper, D. U. et al. Effects of biodiversity on ecosystem functioning: A consensus of current knowledge. Ecol. Monogr. 75, 3–35 (2005).

11. Dornelas, M. et al. Assemblage Time Series Reveal Biodiversity Change but Not Systematic Loss. Science 344, 296–299 (2014).

12. Cardinale, B. J. et al. Effects of biodiversity on the functioning of trophic groups and ecosystems. Nature 443, 989–992 (2006).

13. Lefcheck, J. S. et al. Biodiversity enhances ecosystem multifunctionality across trophic levels and habitats. Nat Commun 6, (2015).

14. Reiss, J., Bridle, J. R., Montoya, J. M. & Woodward, G. Emerging horizons in biodiversity and ecosystem functioning research. Trends Ecol. Evol. 24, 505–514 (2009).

15. Hector, A. et al. BUGS in the Analysis of Biodiversity Experiments: Species Richness and Composition Are of Similar Importance for Grassland Productivity. PLoS ONE 6, e17434 (2011).

16. Scherber, C. et al. Bottom-up effects of plant diversity on multitrophic interactions in a biodiversity experiment. Nature 468, 553–556 (2010).

17. Schuldt, A. et al. Multiple plant diversity components drive consumer communities across ecosystems. Nat. Commun. 10, 1460 (2019).

18. Wan, N.-F. et al. Global synthesis of effects of plant species diversity on trophic groups and interactions. Nat. Plants 6, 503–510 (2020).

19. Lindeman, R. L. The Trophic-Dynamic Aspect of Ecology. Ecology 23, 399–417 (1942).

20. Moore, J. C. & de Ruiter, P. C. Energetic Food Webs: An Analysis of Real and Model Ecosystems. (OUP Oxford, 2012).

21. Barnes, A. D. et al. Biodiversity enhances the multitrophic control of arthropod herbivory. Sci. Adv. 6, eabb6603 (2020).

22. Buzhdygan, O. Y. et al. Biodiversity increases multitrophic energy use efficiency, flow and storage in grasslands. *Nat*. Ecol. Evol. 4, 393–405 (2020).

23. de Vries, F. T. et al. Soil food web properties explain ecosystem services across European land use systems. Proc. Natl. Acad. Sci. 110, 14296–14301 (2013).

24. Kulmatiski, A. et al. Most soil trophic guilds increase plant growth: a meta-analytical review. Oikos 123, 1409–1419 (2014).

25. Crowther, T. W. et al. The global soil community and its influence on biogeochemistry. Science 365, eaav0550 (2019).

26. Henneron, L., Cros, C., Picon-Cochard, C., Rahimian, V. & Fontaine, S. Plant economic strategies of grassland species control soil carbon dynamics through rhizodeposition. J. Ecol. 108, 528–545 (2020).

27. Wardle, D. A. & Van der Putten, W. H. Biodiversity, ecosystem functioning and above-ground-below-ground linkages. in Biodiversity and ecosystem functioning: synthesis and perspectives (eds. Loreau, M., Naeem, S. & Inchausti, P.) 155–168 (Oxford University Press, Oxford, 2002).

28. Fierer, N., Strickland, M. S., Liptzin, D., Bradford, M. A. & Cleveland, C. C. Global patterns in belowground communities. Ecol. Lett. 12, 1238–1249 (2009).

29. Díaz, S. et al. Incorporating plant functional diversity effects in ecosystem service assessments. Proc. Natl. Acad. Sci. 104, 20684–20689 (2007).

30. Grime, J. P. Benefits of plant diversity to ecosystems: immediate, filter and founder effects. J. Ecol. 86, 902– 910 (1998).

31. Weigelt, A. et al. An integrated framework of plant form and function: the belowground perspective. New Phytol. 232, 42–59 (2021).

32. Högberg, P. et al. High temporal resolution tracing of photosynthate carbon from the tree canopy to forest soil microorganisms. New Phytol. 177, 220–228 (2008).

33. Pausch, J. & Kuzyakov, Y. Carbon input by roots into the soil: Quantification of rhizodeposition from root to ecosystem scale. Glob. Change Biol. 24, 1–12 (2018).

34. Pollierer, M. M., Langel, R., Körner, C., Maraun, M. & Scheu, S. The underestimated importance of belowground carbon input for forest soil animal food webs. Ecol. Lett. 10, 729–736 (2007).

35. Zhou, Z. et al. Plant roots fuel tropical soil animal communities. Ecol. Lett. 26, 742–753 (2023).

36. Jucker, T., Bouriaud, O., Avacaritei, D. & Coomes, D. A. Stabilizing effects of diversity on aboveground wood production in forest ecosystems: linking patterns and processes. Ecol. Lett. 17, 1560–1569 (2014).

37. Wambsganss, J., Beyer, F., Freschet, G. T., Scherer-Lorenzen, M. & Bauhus, J. Tree species mixing reduces biomass but increases length of absorptive fine roots in European forests. J. Ecol. 109, 2678–2691 (2021).

38. Beugnon, R. et al. Microclimate modulation: An overlooked mechanism influencing the impact of plant diversity on ecosystem functioning. Glob. Change Biol. 30, e17214 (2024).

39. Jucker, T., Bouriaud, O. & Coomes, D. A. Crown plasticity enables trees to optimize canopy packing in mixed-species forests. Funct. Ecol. 29, 1078–1086 (2015).

40. De Frenne, P. et al. Global buffering of temperatures under forest canopies. *Nat*. Ecol. Evol. 3, 744–749 (2019).

41. Geiger, R., Aron, R. H. & Todhunter, P. The Climate near the Ground. (Rowman & Littlefield, 2009).

42. van der Plas, F. Biodiversity and ecosystem functioning in naturally assembled communities. Biol. Rev. 94, 1220–1245 (2019).

43. Hagan, J. G., Vanschoenwinkel, B. & Gamfeldt, L. We should not necessarily expect positive relationships between biodiversity and ecosystem functioning in observational field data. Ecol. Lett. 24, 2537–2548 (2021).

44. Wardle, D. A. & Lavelle, P. Linkages between soil biota, plant litter quality and decomposition. in Driven by Nature – Plant Litter Quality and Decomposition (eds. Cadisch, G. & Giller, K. E.) 107–127 (CAB International, Wallingford, 1997).

45. Martin-Guay, M.-O. et al. Tree identity and diversity directly affect soil moisture and temperature but not soil carbon ten years after planting. Ecol. Evol. 12, e8509 (2022).

46. Gillerot, L. et al. Forest structure and composition alleviate human thermal stress. Glob. Change Biol. 28, 7340–7352 (2022).

47. Han, M. et al. Linking rhizosphere soil microbial activity and plant resource acquisition strategy. J. Ecol. 111, 875–888 (2023).

48. Millar, C. I. & Stephenson, N. L. Temperate forest health in an era of emerging megadisturbance. Science 349, 823–826 (2015).

49. Anderegg, W. R. L., Kane, J. M. & Anderegg, L. D. L. Consequences of widespread tree mortality triggered by drought and temperature stress. Nat. Clim Change 3, 30–36 (2013).

50. Greenwood, S. et al. Tree mortality across biomes is promoted by drought intensity, lower wood density and higher specific leaf area. Ecol. Lett. 20, 539–553 (2017).

51. Baeten, L. et al. A novel comparative research platform designed to determine the functional significance of tree species diversity in European forests. Perspect. Plant Ecol. Evol. Syst. 15, 281–291 (2013).

52. Petersen, H. & Luxton, M. A comparative analysis of soil fauna populations and their role in decomposition processes. Oikos 39, 287–388 (1982).

53. Bar-On, Y. M., Phillips, R. & Milo, R. The biomass distribution on Earth. Proc. Natl. Acad. Sci. 115, 6506 (2018).

54. Potapov, A. M. et al. Feeding habits and multifunctional classification of soil-associated consumers from protists to vertebrates. Biol. Rev. 97, 1057–1117 (2022).

55. Prada-Salcedo, L. D., Wambsganss, J., Bauhus, J., Buscot, F. & Goldmann, K. Low root functional dispersion enhances functionality of plant growth by influencing bacterial activities in European forest soils. Environ. Microbiol. 23, 1889–1906 (2021).

56. Prada-Salcedo, L. D. et al. Fungal guilds and soil functionality respond to tree community traits rather than to tree diversity in European forests. Mol. Ecol. 30, 572–591 (2021).

57. Jenkins, W. R. B. A rapid centrifugal-flotation technique for separating nematodes from soil. Plant Dis. Report. 48, (1964).

58. Yeates, G. W., Bongers, T., Degoede, R. G. M., Freckman, D. W. & Georgieva, S. S. Feeding Habits in Soil Nematode Families and Genera—An Outline for Soil Ecologists. J. Nematol. 25, 315–331 (1993).

59. Coleman, D. C. & Crossley, D. A. J. Fundamentals of Soil Ecology. (Academic Press, San Diego, 2004).

60. Ganault, P. et al. Relative importance of tree species richness, tree functional type, and microenvironment for soil macrofauna communities in European forests. Oecologia 196, 455–468 (2021).

61. Gillespie, L. M. et al. Tree species mixing affects soil microbial functioning indirectly via root and litter traits and soil parameters in European forests. Funct. Ecol. 35, 2190–2204 (2021).

62. Ehnes, R. B., Rall, B. C. & Brose, U. Phylogenetic grouping, curvature and metabolic scaling in terrestrial invertebrates. Ecol. Lett. 14, 993–1000 (2011).

63. Jochum, M. et al. Decreasing Stoichiometric Resource Quality Drives Compensatory Feeding across Trophic Levels in Tropical Litter Invertebrate Communities. Am. Nat. 190, 131–143 (2017).

64. Lang, B., Ehnes, R. B., Brose, U. & Rall, B. C. Temperature and consumer type dependencies of energy flows in natural communities. Oikos 126, 1717–1725 (2017).

65. Wambsganss, J. et al. Tree species mixing causes a shift in fine-root soil exploitation strategies across European forests. Funct. Ecol. 35, 1886–1902 (2021).

66. Dawud, S. M. et al. Is Tree Species Diversity or Species Identity the More Important Driver of Soil Carbon Stocks, C/N Ratio, and pH? Ecosystems 19, 645–660 (2016).

67. Potapov, A. M., Scheu, S. & Tiunov, A. V. Trophic consistency of supraspecific taxa in below-ground invertebrate communities: Comparison across lineages and taxonomic ranks. Funct. Ecol. 33, 1172–1183 (2019).

68. Brose, U. et al. Predator traits determine food-web architecture across ecosystems. *Nat*. Ecol. Evol. 3, 919–927 (2019).

69. Gauzens, B. et al. fluxweb: An R package to easily estimate energy fluxes in food webs. Methods Ecol. Evol. 10, 270–279 (2019).

70. Schaefer, M. The soil fauna of a beech forest on limestone: trophic structure and energy budget. Oecologia 82, 128–136 (1990).

71. Scheu, S., Ruess, L. & Bonkowski, M. Interactions between microorganisms and soil micro- and mesofauna. in Microorganisms in soils: roles in genesis and functions (eds. Varma, A. & Buscot, F.) vol. 3 253–275 (Springer Berlin Heidelberg, 2005).

72. Angst, G. et al. Conceptualizing soil fauna effects on labile and stabilized soil organic matter. Nat. Commun. 15, 5005 (2024).

73. Joly, F.-X. et al. Detritivore conversion of litter into faeces accelerates organic matter turnover. Commun. Biol. 3, 660 (2020).

74. Lavelle, P. Faunal activities and soil processes: Adaptive strategies that determine ecosystem function. in Advances in Ecological Research, Vol 27 (eds. Begon, M. & Fitter, A. H.) vol. 27 93–132 (Academic Press Ltd-Elsevier Science Ltd, London, 1997).

75. Manning, P. et al. Redefining ecosystem multifunctionality. Nat. Ecol. Evol. 2, 427–436 (2018).

76. Byrnes, J. E. K. et al. Investigating the relationship between biodiversity and ecosystem multifunctionality: challenges and solutions. Methods Ecol. Evol. 5, 111–124 (2014).

77. Laliberté, E. & Legendre, P. A distance-based framework for measuring functional diversity from multiple traits. Ecology 91, 299–305 (2010).

78. Da, R., Fan, C., Zhang, C., Zhao, X. & von Gadow, K. Are absorptive root traits good predictors of ecosystem functioning? A test in a natural temperate forest. New Phytol. 239, 75–86 (2023).

79. Joly, F.-X. et al. Tree species diversity affects decomposition through modified micro-environmental conditions across European forests. New Phytol. 214, 1281–1293 (2017).

80. Joly, F.-X., Scherer-Lorenzen, M. & Hättenschwiler, S. Resolving the intricate role of climate in litter decomposition. Nat. Ecol. Evol. 7, 214–223 (2023).

81. Ponge, J. F. Humus forms in terrestrial ecosystems: a framework to biodiversity. Soil Biol. Biochem. 35, 935– 945 (2003).

82. R Core Team. R: A Language and Environment for Statistical Computing. (R Foundation for Statistical Computing, Vienna, Austria, 2022).

83 Schulz, T., Saastamoinen, M. & Vanhatalo, J. Doors and corners of variance partitioning in statistical ecology. bioRxiv 2021.10.17.464682 (2021) doi:10.1101/2021.10.17.464682.

84. Oberpriller, J., de Souza Leite, M. & Pichler, M. Fixed or random? On the reliability of mixed-effects models for a small number of levels in grouping variables. Ecol. Evol. 12, e9062 (2022).

85. Nakagawa, S. & Cuthill, I. C. Effect size, confidence interval and statistical significance: a practical guide for biologists. Biol. Rev. 82, 591–605 (2007).

86. Lovakov, A. & Agadullina, E. R. Empirically derived guidelines for effect size interpretation in social psychology. Eur. J. Soc. Psychol. 51, 485–504 (2021).

87. Shi, H. & Yin, G. Reconnecting p-Value and Posterior Probability Under One- and Two-Sided Tests. Am. Stat. 75, 265–275 (2021).

88. Lefcheck, J. S. piecewiseSEM: Piecewise structural equation modeling in R for ecology, evolution, and systematics. Methods Ecol. Evol. 7, 573–579 (2016).

89. Vehtari, A., Gelman, A. & Gabry, J. Practical Bayesian model evaluation using leave-one-out cross-validation and WAIC. Stat. Comput. 27, 1413–1432 (2017).

90. Shipley, B. Confirmatory path analysis in a generalized multilevel context. Ecology 90, 363–368 (2009).

