## Supplementary material for "Resource economics of tree communities control soil food web multifunctionality in European forests": Tree biodiversity effects on soil food web multifunctionality - Supplementary Information

### Supplementary Methods

#### Biomass calculation and trophic classification of soil organisms

*Microorganisms*

Bacterial and fungal dry biomass in each plot was measured using phospholipid fatty acid (PLFA) analysis[1] as described by Prada-Salcedo et al. (2021)[2]. Briefly, soil samples were stored at -20 °C within 72 h after sampling, and PLFAs were extracted from 2 g of freeze-dried soil pooled at the plot-level using the method of Bligh & Dyer (1959)[3]. Individual PLFAs were then identified and quantified by GC/MS (Agilent, HP DB5 column), and were converted to nmol PLFA per g of dry soil. The PLFAs i15:0, a15:0, 15:0, i16:0, 16:1ω7, 16:1ω9, i17:0, a17:0, 17:0, and 18:1ω7 were pooled to quantify bacterial biomass using a conversion factor of 363.6 nmol PLFA per 1 mg C[1]. We also separated between bacterial PLFAs specific to gram-negative (16:1ω7, and 18:1ω7), and gram-positive (i15:0, a15:0, 10Me15:0, i16:0, 10Me16:0, i17:0, a17, 10Me17:0, and 10Me18:0)[4]. The ratio of gram-positive to gram-negative bacterial PLFAs was then used to partition their respective biomass from bacterial biomass. The PLFA 16:1ω5 was used to quantify arbuscular mycorrhizal (AM) fungal biomass using a conversion factor of 38.0 nmol PLFA per 1 mg C[5]. The PLFAs 18:2ω6,9 and 18:1ω9 were pooled to quantify non-AM fungal biomass using a conversion factor of 20.06 nmol PLFA per 1 mg C[6]. Microbial C mass per unit surface area (g C $m^{-2}$) was then calculated by multiplying microbial C mass per unit soil mass (mg C $g^{-1}$ dry soil) with soil bulk density (g $cm^{-3}$) and soil volume per unit surface area ($cm^3$ $m^{-2}$). We finally converted microbial C mass into microbial dry mass assuming a C content of 46 and 41% of dry matter for bacteria and fungi, respectively[7]. Across all plots, the ratio of fungal to bacterial biomass C was found to be 9.9 ± 4.8 (median ± standard deviation), which is close to the previously reported value of 8.5 based a meta-analysis using PLFA data[8]. The median of total microbial biomass was found to be 93 ± 66 g C microbial $m^{-2}$, which is within the same range than previously reported values for forest ecoystems[9] (51 ± 6 for boreal forests, 89 ± 26 for temperate coniferous forests, and 82 ± 9 for temperate deciduous forests).

Plot-specific fungal community data based on metagenomic sequencing and bioinformatics analyses[10] performed according to Prada-Salcedo et al. (2021)[11] were used to partition non-AM fungal biomass into five trophic guilds: ericoid mycorrhizal fungi, ectomycorrhizal fungi, general saprotrophic fungi, wood saprotrophic fungi, and plant pathogenic fungi. Briefly, soil DNA was extracted from samples pooled at the plot-level, and the internal transcribed spacer (ITS) region 2 was amplified by PCR using the primers P5-5N-ITS4, P5-6N-ITS4, P7-3N-fITS7, and P7-4N-fITS7. Amplicons were sequenced using an Illumina MiSeq System according to standard protocols, and the sequences were processed using the metabarcoding amplicon pipeline DeltaMP, and clustered into operational taxonomic units (OTUs) according to the UNITE database[12]. The OTUs were then assigned to trophic guilds using the FUNGuild tool[13], and the biomasses of each fungal trophic guild were calculated by multiplying their relative abundance (number of reads of the trophic guild divided by the total number of reads for all five trophic guilds) by the total non-AM fungal biomass[14–16]. The OTUs that potentially switch between trophic modes during their life history and OTUs that lacked sufficient taxonomic classification for functional description were not considered for the calculation of relative abundance. The biomass of mycorrhizal fungi was calculated as the sum of ericoid mycorrhizal, ectomycorrhizal and arbuscular mycorrhizal fungal biomass, while the biomass of decomposer fungi was calculated as the sum of general and wood saprotrophic fungal biomass. On average across all plots, we found mycorrhizal fungi, followed by saprotrophic fungi, to be the dominant trophic guilds in term of contribution to fungal biomass, in accordance with previous studies on forest stands dominated by ectomycorrhizal tree species[14,16,17].

*Soil fauna*

For each plot, we measured the dry mass of each faunal guild. Because the calculation of faunal metabolic rates was based on a model using faunal fresh mass[18], we also calculated fresh mass of each faunal guild.

The fresh mass of nematode families per unit soil mass (µg $g^{-1}$ dry soil) was calculated by multiplying their density (number of individuals $g^{-1}$ dry soil) with individual fresh body mass values (µg $individual^{-1}$) retrieved from the Nemaplex database (http://nemaplex.ucdavis.edu, Supplementary Table 8). These individual fresh body mass values are the mean of all species values available for each family, which were calculated using the following allometric equation[19]:

$$M = \frac{L \times D^2}{1.6 \times 10^6} \qquad (1)$$

where $M$ is the fresh mass (in µg) per individual, $L$ is the nematode length (in µm) and $D$ is the greatest body diameter at the largest body part (in µm). Nematode biomass per unit surface area (µg $m^{-2}$) was computed based on soil bulk density as described above, and we converted nematode fresh mass to dry mass assuming a dry matter content of 25%[20]. Nematode families were assigned to five trophic guilds: herbivores, bacterivores, fungivores, omnivores, and carnivores[21] (Supplementary Table 2, Supplementary Table 8).

The dry mass per unit surface area of microarthropods was calculated by multiplying their density (individual number $m^{-2}$) with individual dry body mass values (µg $individual^{-1}$). For collembola species, the mean individual dry body mass ($M$, in µg) was calculated from the mean body length ($L$, in mm) retrieved from the BETSI database (https://portail.betsi.cnrs.fr/) using the following allometric equation (Supplementary Table 9):

$$M = aL^{\mathrm{b}} \qquad (2)$$

where $a$ is the normalisation coefficient and $b$ is the scaling exponent. Abdomen length of Symphypleona was used in the original equations and was assumed to be 0.83 of the total body length. Two sets of coefficients derived from two independent studies[22,23] were used for each species ($a_1$, $b_1$ and $a_2$, $b_2$) and the two estimates of dry body mass were averaged[24]. Collembola fresh body mass was calculated from the resulting average by dividing it by the proportion of the dry weight using values from the literature[23,24] (Supplementary Table 9). For mite taxa, mean individual dry body mass were estimated based on a global dataset of soil fauna[25]: 5.3, 7.7 and 1 µg for Oribatida, Mesostigmata, and Prostigmata/Astigmata, respectively. Mite fresh body mass was calculated as described above assuming a dry matter content of 43.1%[26]. Collembola species were classified in four ecological groups (Epedaphic, Hemiedaphic, Euedaphic and Predators) based on Potapov et al. (2022[7], Supplementary Table 9). Microarthropods taxa were then assigned to seven trophic guilds[7]: Acari: Oribatida (microbi-detritivores), Acari: Mesostigmata (carnivores), Acari: Prostigmata & Astigmata (omnivores), Collembola: Epedaphic (fungivores), Collembola: Hemiedaphic (microbi-detritivores), Collembola: Euedaphic (fungivores), and Collembola: Predators (carnivores, Supplementary Table 2).

The abundances and biomasses of soil macroinvertebrates were measured as described by Ganault et al. (2021)[27]. All macroinvertebrate individuals were gently wiped and weighed to the nearest 0.01 mg to measure their fresh body mass. Since heavy soil materials can represent a significant fraction of faunal fresh mass for soil feeders, we corrected the fresh mass of endogeic and anecic earthworms by multiplying it with a conversion factor of 0.67 g empty gut fresh mass per g full gut fresh mass[25]. We converted macroinvertebrate fresh body mass ($M$) to dry body mass ($DM$) using the following equation[28]:

$$DM = e^{\frac{log(M) - a}{b}} \qquad (3)$$

where the $a$ and $b$ coefficient values are 0.9282 and 1.0899 for earthworms, and 0.6111 and 1.0213 for all other taxa, respectively. Lumbricidae species were classified in three ecological groups (Epigeic, Endogeic, and Anecic) based on the DriloBASE database (http://taxo.drilobase.org, Supplementary Table 10). Macroinvertebrates taxa were then assigned to 23 trophic guilds[7]: Small Araneae (< 1.14 mg, carnivores), Large Araneae (> 1.14 mg, carnivores), Opiliones (carnivores), Diplopoda (detritivores), Chilopoda: Geophilomorpha (carnivores), Chilopoda: Scolopendromorpha (carnivores), Chilopoda: Lithobiomorpha (carnivores), Isopoda (detritivores), Dermaptera (omnivores), Blattodea (detritivores), Formicidae (omnivores), Coleoptera: Carabidae (carnivores), Coleoptera: Staphylinidae (carnivores), Coleoptera: Elateridae (omnivores), Coleoptera: Scarabaeidae/Geotrupidae (herbi-detritivores), Coleoptera: Curculionidae (herbivores), Lepidoptera (herbivores), Diptera (omnivores), Lumbricina: epigeic (detritivores), Lumbricina: anecic (humi-detritivores), Lumbricina: endogeic (humi-detritivores), Gastropoda: snails (detritivores), and Gastropoda: slugs (detritivores, Supplementary Table 2).

**Calculation of metabolic rates and assimilation efficiencies**

Plot-specific soil microbial respiration and biomass data from Gillespie et al. (2021)[29] was used to compute the metabolic rates of microbial groups. For heterotrophs, the metabolic rate is equal to the rate of respiration because heterotrophs obtain energy by oxidizing carbon compounds[30]. To measure soil microbial basal respiration, 10 g of fresh soil was incubated at 25°C and 80% of water holding capacity during 6 h, and $CO_2$ concentration was quantified after 2 and 6 h using a MicroGC (S-Series, SRA Intruments, Marcy l'Etoile, France). A similar incubation but including the addition of glucose (mass equivalent of 15 mg C) was used to estimate soil microbial biomass based on the substrate-induced respiration method using the following equation[31]:

$$C_{microbial} = MIRR \times 40.04 + 0.37 \quad (4)$$

where '$C_{microbial}$' is the active microbial biomass (µg $C_{microbial}$ $g^{-1}$ dry soil) and '$MIRR$' is the substrate-induced maximum initial respiration rate (µl C-$CO_2$ $g^{-1}$ dry soil $hour^{-1}$). The metabolic quotient of soil microbes ($qCO_2$, mg C-$CO_2$ $g^{-1}$ $C_{microbial}$ $hour^{-1}$) was calculated as basal respiration divided by microbial biomass, and was standardized to the plot-specific mean soil temperature using the following equation[32]:

$$qCO_{2_{field}} = qCO_{2_{incubation}} \times Q_{10}^{\frac{T_{field}-T_{incubation}}{10}} \quad (5)$$

where '$qCO_{2\ field}$' and '$qCO_{2\ incubation}$' are the $qCO_2$ for field and incubation conditions, respectively, '$T_{field}$' and '$T_{incubation}$' are the plot-specific mean soil temperature during the growing season (°C, see 'microclimate' section below) and the incubation temperature (25 °C), respectively. '$Q_{10}$' is the temperature sensitivity coefficient, and a $Q_{10}$ value of 2 was used[32]. Across all plots, we found $qCO_{2\ field}$ to be 1.90 ± 0.26 mg C-$CO_2$ $g_{-1}$ $C_{microbial}$ $h^{-1}$ (median ± standard deviation), which is close to the previously reported value of 1.81 based a previous meta-analysis[32].

Assuming $qCO_2$ to be two times lower for fungi than bacteria[33], the $qCO_2$ of fungi and bacteria were calculated using the following equations:

$$qCO_{2_{fungi}} = \frac{qCO_{2_{field}}}{2 \times f_{bacteria} + f_{fungi}} \quad (6)$$

$$qCO_{2_{bacteria}} = 2 \times qCO_{2_{fungi}} \quad (7)$$

where '$f_{bacteria}$' and '$f_{fungi}$' are the bacterial and fungal fraction of total microbial biomass based on PLFA analysis, respectively. Based on this method, we found that bacterial respiration represent 16.8 ± 6.4 % (median ± standard deviation across all plots) of total soil microbial respiration, which is within the same range than previously reported values (12.2 ± 4.2 %) for soils within the pH range (~4.0) from a previous study using the selective inhibition method[34]. The metabolic rates (g C-$CO_2$ $m^{-2}$ $day^{-1}$) of microbial trophic guilds were then calculated as their biomass multiplied by their metabolic quotient, and were converted to kJ $m^{-2}$ $day^{-1}$ using a conversion factor of 1 g C = 39 kJ based on the enthalpy of glucose combustion[35].

The metabolic rates of faunal groups were computed from individual body mass, environmental temperature and phylogenetic grouping using the following equation[30]:

$$I = e^{\ln i_0 + a \times \ln M - \frac{E}{kT}} \quad (8)$$

where $I$ is the metabolic rate (in J hour$^{-1}$), $i_0$ is a normalization factor, $a$ is the allometric exponent, $M$ is the fresh body mass (in mg), $E$ is the activation energy (eV), $k$ is Boltzmann's constant (8.62 * 10$^5$ eV Kelvin$^{-1}$), and $T$ is the environmental temperature (in Kelvin). Taxon-specific or otherwise general parameter values fitted from a large soil invertebrate respiration dataset[18] were used for $i_0$, $a$, and $E$ (Supplementary Table 11). Plot-specific values of growing season soil temperature were used for $T$ (see below). For macroinvertebrates, metabolic rates were calculated for each individual, and the community metabolism ($X$, in kJ m$^{-2}$ day$^{-1}$) of each trophic guild was calculated as the sum of all individual metabolic rates divided by the sampled area. For nematodes and microarthropods, metabolic rates were calculated for each taxon using the mean individual fresh body mass, and the community metabolism of each trophic guild was calculated as the sum of the products of taxon-specific metabolic rate and taxon density.

Assimilation efficiencies ($e_a$) specific to each food type (plant-derived resource or prey) for faunal consumers were calculated based on resource stoichiometry using the following equation[36] (Supplementary Table 12):

$$e_a = \frac{\mathrm{e}^{(0.471\times \text{Food N}-2.097)}}{1+\mathrm{e}^{(0.471\times \text{Food N}-2.097)}} \quad (9)$$

where 'Food N' is the N content (%) of the food resource. For plant-derived resources, we used plot-specific N content measured as described above. For microbial and faunal trophic guilds, we used generic N content data available from the literature[7].

A temperature correction of assimilation efficiency ($\varepsilon_{cor}$) was calculated using the following equation[37]:

$$\varepsilon_{cor} = \frac{e^{\acute{\varepsilon}_{0,j}} e^{E_{\varepsilon,j}\frac{T-T_0}{kTT_0}}}{1+e^{\acute{\varepsilon}_{0,j}} e^{E_{\varepsilon,j}\frac{T-T_0}{kTT_0}}} - \varepsilon_{0,j} \quad (10)$$

where $\acute{\varepsilon}_0$ is the normalization constant of the assimilation efficiency, $E_\varepsilon$ is the activation energy for assimilation efficiency (eV), $T$ is the environmental temperature (K), $T_0$ is the temperature normalized to 20°C (293.15 K), $k$ is the Boltzmann's constant (8.62 * 10$^5$ eV K$^{-1}$), and $\varepsilon_0$ is the assimilation efficiency at 20°C. The subscript $j$ refers to the different consumer types (herbivores, detritivores, or carnivores). Consumer type-specific parameter values from the literature[37] were used for $\acute{\varepsilon}_0$, $E_\varepsilon$, and $\varepsilon_0$ (Supplementary Table 13). Plot-specific values of soil temperature during the growing season converted to Kelvin were used for $T$ (see below). Assimilation efficiencies were then standardized to the plot-specific mean soil temperature during the growing season by adding the correction coefficient $\varepsilon_{cor}$ to $e_a$.

**Plant-derived resources**

The biomass and N content of living plant fine roots and dead roots in each plot was measured as described by Wambsganss et al. (2021a, c)[38,39]. Briefly, roots were sampled during the phenological spring 2017 by taking one core of the soil layer (10 cm depth, 5.3 cm diameter) in each of the same five subplots used for soil sampling (Supplementary Fig. 3b). Living fine roots (absorptive roots belonging the first three root orders)[40] of all plants (dominated by woody tree species, but also with some herbaceous plants) were washed and visually sorted into dead and live roots, before being dried for at least 72 h at 40 °C and weighed to estimate the biomass of living plant fine roots and dead roots per unit surface area. Living plant fine roots were then pooled at the plot level, ground and their N content was analyzed by dry combustion using an elemental analyzer (EA, Elementar Vario El Cube). We assumed equal N content of living plant fine roots and root litter based on a global synthesis of plant root data[41].

The biomass and N content of the litter layer in each plot was measured as described by Gillespie et al. (2021)[29]. Briefly, four 15 × 15 cm square samples of the litter layer predominantly composed of dead leaves were sampled during phenological spring 2017 in each of the same five subplots used for soil sampling (Supplementary Fig. 3b). Litter material was then dried at 60°C, weighed to estimate the biomass of dead leaves per unit surface area, before being ground, and analyzed by dry combustion to mesure its N content.

The biomass and N content of dead wood in each plot was retrieved from the FunDivEurope database (https://data.botanik.uni-halle.de/fundiveurope/). Briefly, the volume per unit surface area of woody debris, that are all standing dead trees and snags, and all stumps and other dead wood pieces lying on the forest floor, was measured during summer 2012 in two circular subplots (radius of 7 m) located in opposite corners of each plot. The fresh volume and dry mass of a woody debris subsample were also measured to estimate wood density, and the biomass of dead wood per unit surface area was calculated as woody debris volume per unit surface area divided by wood density. The woody debris subsample was then ground, and its N content was analyzed by dry combustion.

The biomass and N content of soil organic matter (SOM) in each plot was measured as described by Dawud et al. (2016)[42]. Briefly, mineral soil (A horizon) was sampled during summer 2012 by taking one soil core (10 cm deep, 3.6 cm diameter) in each of nine subplots, and was then sieved through 2 mm and pooled at the plot-level, dried at 60°C for at least 48 h, ground and its C and N contents were analyzed by dry combustion. The C stock of SOM was calculated based on soil C content and bulk density, and SOM dry mass was estimated assuming a C content of SOM of 50%.

**Reconstruction of the soil food web topology and interaction strengths**

To reconstruct the food web interaction matrix representing the topology and trophic interaction strengths of the food web, a set of five matrices with food resources in rows and consumers in columns were first generated, and then merged by multiplying them together (see Fig. 5 in Potapov 2022[43] for an illustration). Each matrix represents one trait dimension: phylogenetically defined feeding preferences, density(biomass)-dependence, predator–prey interactions related to body mass ratio and hunting strategies, prey protection mechanisms, and spatial niche overlap related to vertical stratification. To generate the matrices, we used multiple parameters mostly derived from Potapov *et al.* (2022)[7] corresponding to organism traits (Supplementary Table 3). Food-web reconstruction was carried out separately for each plot to account for plot-specific density(biomass)-dependence.

First, we generated a resource matrix based on phylogenetically inherited differences in feeding preferences of consumers for various food resources[7], including living plant fine roots roots (LivR), photosynthates & rhizodeposits (PR), dead leaves (LLit), dead roots (RLit), dead wood (WLit), soil organic matter (SOM), bacteria (Ba), mycorrhizal fungi (FuM), general saprotroph fungi (FuS), wood saprotroph fungi (FuW), plant pathogenic fungi (FuP), and animal prey (Fauna, Supplementary Table 3). For each trophic guild, feeding preferences of the corresponding consumers for different food resources (plant-derived resources and/or other consumers) were determined assuming that additional food resources (coded as 0.5) are five times less important than the main resources (coded as 1)[43]. For omnivores, *i.e.* organisms that have developed predation capabilities while also feeding on other food resources (both coded as 1), animal diet was assumed to represent 50% of the node budget[44]. Feeding preferences were then scaled so that preferences of a given consumer for all food resources summed to 1. Feeding preferences were defined based on Potapov *et al.* (2022)[7] for faunal trophic guilds, while feeding preferences of microbial trophic guilds were defined based on previous isotopic labeling studies[45]. Specifically, photosynthates and rhizodeposits were defined as the primary food resources for mycorrhizal fungi[46,47] and gram-negative bacteria[48–50]. Decomposer fungi were defined to feed primarily on litter[51–53] and rhizodeposits[49,53], but also secondarily on soil organic matter[53,54]. Gram-positive bacteria were defined to feed primarily on soil organic matter[48,51] and litter[51,52].

Second, we generated a biomass matrix corresponding to density dependent preferences of consumers for various food resources by filling each row with the biomass of the corresponding trophic guild.

Third, we generated an allometric matrix to modulate the strength of carnivore trophic interactions based on predator–prey mass ratio[55] (PPMR) and predator traits. The optimum

PPMR was first calculated for each faunal predator based on their mean body mass (in g, $log_{10}$-transformed values given in Supplementary Table 2 as 'Mass mean') using the following equation[55]:

$$log_{10}PPMR_{optimum} = -0.159 \times log_{10}\text{body mass} + 0.330 \quad (11)$$

For each faunal predator, the mean body mass of its optimum prey was then calculated by subtracting $PPMR_{optimum}$ from the predator mean body mass, while the body mass standard deviation of its optimum prey was calculated by multiplying $PPMR_{width}$ ('Mass SD' in Supplementary Table 2) with predator mean body mass. For most predators, the body mass standard deviation of its optimum prey was set to be the same as the body mass standard deviation of the predator ($PPMR_{width}$ = 1). However, the values of $PPMR_{optimum}$ and $PPMR_{width}$ was first corrected according to hunting strategies by multiplying them with the correction coefficients based on predator traits (Supplementary Table 2). The distributions of optimum and real prey body mass were then modelled using the mean and standard deviation of optimum and real body mass, assuming these distributions to follow a log-normal pattern. Overlaps of the size distributions of optimum and real prey were then calculated for all potential predator–prey interactions, and overlap proportion [0;1] was used as a proxy of interaction strength.

Fourth, we generated a protection matrix to modulate the strength of carnivore trophic interactions that they can be considerably reduced by prey protective traits. An empty matrix was filled with ones, and it was then multiplied by each of the correction coefficients based on protective traits (Supplementary Table 2). The lower the coefficient, the higher the protection.

Fifth, a spatial matrix was generated to modulate the strength of trophic interactions based on consumer-resource overlap in habitat niches. Pairwise Bray–Curtis dissimilarities for all nodes were calculated based on the vertical distribution of each node in soil (euedaphic, 'eu'), litter (hemiedaphic, 'hemi'), or surface (epiedaphic, 'epi'). Vertical distribution was coded as 0 (nearly absent), 0.5 (occasional) and 1 (present, Supplementary Table 2). Note that a given node can occur in multiple vertical layers, owing to organism mobility for instance. Reverse dissimilarity [0;1], as a measure of spatial overlap, was used as a proxy for interaction strength.

Finally, the five matrices were combined to generate the food web interaction matrix. All the matrices, except the resource matrix, were first multiplied together. The resulting matrix was then multiplied by the resource matrix separately for basal, microbial and faunal types of food resources, and then scaled so that trophic guild preferences for all food resources of a given type summed to the proportion of this given resource type in the trophic guild diet. This approach allows to avoid the overestimation of the contribution of basal and/or microbial resources to the diet due to their much larger biomass[43]. The full R script used to generate the trophic interaction matrix will be made available upon acceptance for publication.

**Calculation of soil food web energy fluxes, trophic functions and multifunctionality**

Energy fluxes to each feeding guild of the food web were calculated using the following equation[56]:

$$F = \frac{1}{e_a} \times (X + L) \quad (12)$$

where $F$ (kJ $m^{-2}$ $day^{-1}$) is the total flux of energy into the feeding guild, $e_a$ is the diet-specific assimilation efficiency, $X$ is the community metabolism of the feeding guild, and $L$ is the energy loss to predation.

More specifically, energy fluxes of individual trophic interactions were calculated using the following equation[57]:

$$\sum_j F_i W_{ji} e_{a\,ji} = X_i + \sum_j F_j W_{ij} \quad (13)$$

where $F_i$ and $F_j$ are the sum of all ingoing fluxes to feeding guild $i$ and $j$, respectively, '$e_{a\,ji}$' is the diet-specific assimilation efficiency of the food resource $j$ consumed by feeding guild $i$, $X_i$ is the energy lost through community metabolism of the feeding guild $i$, and $W_{ji}$ and $W_{ij}$ are the proportions of $F_i$ and $F_j$ that are obtained from feeding guilds $j$ and $i$, respectively. This equation is solved in two stages: first, the sum of ingoing fluxes for each trophic guild ($F_i$) is computed.

Second, individual fluxes for each pairwise consumer-food resource interaction ($F_{ji}$) are calculated using consumer preferences ($W_{ji}$) based on the trophic interaction matrix. The calculation of energy fluxes starts with the top predators and proceeds downward to the trophic guilds with the lowest trophic position. The loss to predation in these lower feeding guilds was synonymous with energy fluxes directed toward their consumer feeding guilds. Energy fluxes were computed using the '*fluxing*' function of the '*fluxweb*' package[57].

We quantified soil food web multifunctionality using the 'averaging' approach[58] by estimating the average standardized value of 10 trophic functions of the soil food web: mycorrhizal symbiotrophy, root pathogenicity, rhizophagy, litter decomposition, litter engineering, SOM decomposition, soil engineering, bacterivory, fungivory and carnivory. All of the ten trophic functions were first standardized using the following equation[58]:

$$F_i = \frac{rawF_i - \min(rawF)}{\max(rawF) - \min(rawF)} \quad (14)$$

Where $F_i$ and $rawF_i$ are the standardized and unstandardized function values of plot *i*, respectively, and min(rawF), and max(rawF) are the minimal and maximal values of the function across all plots, respectively. The averaged multifunctionality index was then calculated as the mean of the 10 standardized trophic functions. To check that the results were robust to multiple alternative approaches, we also quantified soil food web multifunctionality using the 'threshold' approach[58] by estimating the number of trophic functions whose value exceeded 30, 50, 70 and 90% of the maximal value. Calculations for the 'threshold' approach were performed using the '*getFuncMaxed*' function of the '*multifunc*' package[59]. For both approaches, the maximum value of the trophic function was calculated as the mean of the four highest values to reduce the influence of outliers[58].

**Tree functional (trait-based) diversity and composition**

Leaf trait data for each tree species in each plot was mostly derived from *in situ* measured values retrieved from the FunDivEurope database (https://data.botanik.uni-halle.de/fundiveurope/). Leaf nitrogen content (LNC) was measured for each tree species in all plots, while specific leaf area (SLA), and leaf dry matter content (LDMC) were measured for Finland and Romania plots only. The three leaf traits were measured on sun-exposed leaves of each tree species collected in spring 2015 using standard protocols[60]. Briefly, ten individual trees of each target species present were selected in each plot, and five leaves were sampled for each of these individual trees. Each sampled leaf was gently wiped after overnight immersion in tap water, weighed to measure leaf fresh mass, and scanned with a flat-bed scanner (resolution 800 dpi). Scans were then analysed with the WinFOLIA software (Regents Instruments, Québec, Canada, 2009) to obtain leaf area. Subsequently, each leaf was dried for at least 72 h at 40 °C, and weighed to measure leaf mass. SLA and LDMC were then calculated as leaf area divided by dry mass, and leaf fresh mass divided by dry mass, respectively. Leaf samples of each tree were then pooled, ground and their N content was analysed by dry combustion. Trait values were averaged at the plot level (one value per tree species and plot). For Italy and Poland, SLA and LDMC values of each of the seven tree species of these two countries were retrieved using the TRY plant trait database[61] (https://www.try-db.org/, accession date October 2019, request n° 6303), and we then used the average of trait values for each individual trait and tree species.

Root trait data was derived from *in situ* values measured in each plot[39]. Briefly, the six root traits were measured on tree fine roots (absorptive roots belonging to the first three root orders[40]) of each species using standard protocols as described by Wambsganss et al. (2021)[39]. Root tips colonised by ectomycorrhizal fungi (EcM) were first visually identified and counted on representative subsamples based on the presence or absence of a fungal sheath for the 12 species known to associate with EcM (*Acer pseudoplatanus* occurring in Romania associate with arbuscular mycorrhizal fungi). Thereafter, roots were scanned in water with a flat-bed scanner (resolution 800 dpi), and scans were analysed with the WinRhizo software (Regents Instruments, Québec, Canada, 2009) to obtain root length, volume and diameter. Subsequently, all root samples were dried for at least 72 h at 40 °C and weighed to obtain root dry mass. Samples were then ground, and their N content was analysed as described above to obtain root N content (RNC). Root tissue density (RTD) was calculated as root dry mass divided by root volume. Specific root length (SRL) was calculated as root length divided by root dry mass. Measured root diameter values were averaged to obtain mean root diameter ($D_m$). Root length density (RLD) was calculated as the length of all fine roots recovered in the sampled soil core divided by the soil core volume. Ectomycorrhizal colonisation intensity (%Myc) was calculated as the number of root tips colonised by ectomycorrhizal fungi (EcM) divided the length of roots examined.

For each plot, we quantified tree functional diversity based on these nine traits using the functional dispersion (FDis) index, which is the mean distance of each species to the centroid of all species in the multifunctional trait space[62]. For monocultures, the computation of the FDis index always yielded a value equal to zero because of the absence of functional dispersion when only one tree species is present. We also calculated community-weighted mean (CWM) of each trait to characterize tree functional composition[63]. FDis and CWM values were computed based on the relative basal area of tree species using the '*dbFD*' function of the '*FD*' package[62]. We also performed a Principal Component Analysis (PCA) on all CWM traits using the '*principal*' function of the '*psych*' package[64]. This allowed to simplify the tree functional composition into two dimensions[39,65] (Extended data Fig. 2a): (1) a leaf economics spectrum (LES, 45.2 % of variation), which ranged from slow/conservative leaf attributes (N-poor and dry matter-rich leaves with low leaf area per unit mass) to fast/acquisitive leaf attributes (N-rich and dry matter-poor leaves with high leaf area per unit mass), and which was also aligned here with a fine-root gradient of belowground resource foraging strategies ranging from low foraging efficiency (thick fine roots with low length per unit mass) to high foraging efficiency (thin fine roots with high length per unit mass); (2) a fine-root economics spectrum (RES, 31.5 % of variation), which ranged from slow/conservative fine-root attributes (N-poor fine roots with low tissue density) to fast/acquisitive fine-root attributes (N-rich fine roots with high tissue density), and which was also aligned here with a fine-root gradient of soil exploration strategies ranging from 'do-it-yourself' attributes (high root length density and low ectomycorrhizal colonisation intensity) to 'outsourcing' attributes (low root length density and high ectomycorrhizal colonisation intensity). The LES axis was positively indeed related to LNC, SLA and SRL, and negatively to LDMC and $D_m$, while the RES axis was positively related to RNC and %Myc, and negatively to RTD and RLD (Supplementary Table 5).

**Tree and understorey vegetation**

For each plot, all tree stems ≥ 7.5 cm in diameter at breast height (DBH) were identified to species, and their height and DBH were measured in spring 2017 for Finland, Romania and Italy, and in spring 2018 for Poland. Height and DBH measurements were used to estimate tree aboveground biomass (AGB) based on published allometric equations[66], as well as the basal area of each tree. Tree canopy density was estimated using the leaf area index (LAI), with five replicated measurements of LAI performed in each plot using a Plant Canopy Analyzer LAI-2000 (Li-Cor Inc., Lincoln, NE, USA)[67] during spring 2012 for Finland and Italy, and spring 2013 for Poland and Romania. Aboveground litter production of tree communities was quantified by regularly collecting canopy tree litterfall (including leaf, wood and reproductive parts) over one entire year from October 2011 to November 2012 using five litter traps (0.5 $m^2$, 1 m above the ground) per plot[68]. Collected litter was pooled by plot, sorted by species and litter type (*i.e.*, leaves, twigs, and reproductive parts), and weighed after drying at 65 °C. The percentage cover of understorey plant species (< 1.3 m tall individuals) was characterized in three 5 m × 5 m quadrats in each plot between May and August 2012[69]. At the same time, the aboveground biomass and C:N ratio of woody and herbaceous vegetation were determined from samples cut in a smaller area (0.5 m × 0.5 m) within each of these three quadrats. Understorey plant diversity was quantified using the species richness calculated at the plot-level (combined across the three sampling quadrats). Total root biomass was quantified as the sum of absorptive (first three orders) and transport (higher orders) roots of both herbaceous and woody plant species[38].

#### Environmental drivers

*Abiotic conditions*

In each plot, altitude was recorded during plot selection. Plot-specific mean annual temperature and precipitation were retrieved using the WorldClim database[70] (https://www.worldclim.org/). The texture of mineral soil texture was measured in each plot using the laser diffraction method[71].

We performed a PCA as described above to reduce the dimensionality of abiotic properties into two dimensions[39,65] (Supplementary Fig. 1a): (1) a soil texture gradient ranging from coarse to fine, coordinated with a macroclimate gradient ranging from dry and cold to wet and hot (PC1, 70.8% of variation), (2) a gradient of temperature and altitude ranging from cold and high elevation to warm and low elevation (PC2, 19.1% of variation).

*Microclimate*

Microclimate was measured in the center of each plot at all four countries during an entire year from spring 2018 to spring 2019. Soil temperature and moisture (0-14 cm soil depth), and air temperature (50 cm above the ground) were measured every 15 min using TMS-4 data loggers (TOMST, Prague, Czech Republic)[72], with a special version allowing temperature measurements 50 cm above the ground. Specifically, the bottom part of the TMS logger constitutes a probe measuring volumetric soil moisture from the soil surface until a depth of 14 cm, and also includes a soil temperature probe located in the middle at 8 cm depth. Mean annual values and mean growing season (daily mean air temperatures >5 °C) values were calculated for soil and air temperature, and soil moisture. Daily minimum values instead of means were used in the calculation for soil moisture. We did this because water tends to accumulate around the sensors after rain events, thus leading to an overestimation of soil moisture when using all values measured per day[72].

The microclimatic dataset was incomplete due to missing values following wild boar damage on TMS-4 data loggers in six plots (five in Poland, and one in Italy). These missing values were replaced based on a regularized iterative multiple correspondence analysis[73] using the '*imputePCA*' function of the '*missMDA*' package[74].

We performed a PCA as described above to reduce the dimensionality of microclimatic properties into a single dimension (PC1, 63.6% of variation, Extended data Fig. 2b). This microclimate axis ranged from 'cold and wet' to 'hot and dry', and was positively related to soil and air temperature, and negatively to soil moisture, both for annual and growing season values.

*Leaf litter quality*

Litter properties of each tree species were measured for each location using standard protocols as described by Joly et al. (2017)[68] on freshly fallen leaf litter of each species. Litter material was collected at tree species-specific peak leaf litter fall between October 2011 and November 2012, in close vicinity of the experimental plots, and was processed as described above (see 'Tree and understorey vegetation' section). Briefly, total C and N concentrations were measured with a flash CHN Elemental Analyser (Flash EA1112 Series, ThermoFinnigan, Milan, Italy). After a mineralization step, phosphorus (P) concentration was measured colorimetrically with an autoanalyzer (SmartChem 200, Alliance Instruments, Rome, Italy). Water soluble compounds (WSC), cellulose, hemicellulose and lignin fractions were determined using a fiber analyzer (Fibersac 24, Ankom, Macedon, NJ, USA) according to the Van Soest extraction protocol[75]. Calcium (Ca) concentration was measured using an atom absorption spectrometer (AAS, iCE 3000 series, Thermo-Scientific, China). Concentrations of condensed tannins were measured spectrophotometrically using the butanol-HCl method[76]. For total phenolics, samples of 0.5 g were soaked in 30 mL of 50% methanol, shaken for 2 h, filtered (filter number 112, Durieux, Torcy, France), and the extracts were analyzed spectrophotometrically with the FolineCiocalteau reagent by using gallic acid as a standard. Leaf litter pH was measured according to Cornelissen et al. (2006)[77]. Water holding capacity (WHC) was measured by placing intact leaves in non-hermetically lidded large plastic containers and spraying them three times a day with deionized water. The amount of water retained by the leaves was measured after 48 h. Except for pH and physical traits, which were measured on intact leaf litter, all other chemical analyses were performed on subsamples ground to obtain a uniform particle size of 1 mm (Cyclotec Sample Mill, Tecator, Höganäs, Sweden).

We then quantified the functional diversity and composition of tree leaf litter for each plot using the FDis index and CWMs of each litter property as described above for tree functional traits, except that the computation was based on the relative leaf litter mass of the different tree species collected in litter traps[68], rather than the relative basal area of trees. We also performed a PCA similar to that described above for tree functional traits to reduce the dimensionality of litter CWM properties into one dimension: a spectrum of leaf litter nutritional qualities[78] (PC1, 42.5% of variation%, Extended data Fig. 2c). The leaf litter quality axis ranged from low to high nutritional quality, and was related positively to WHC and the concentrations of N, P, K, Mg, WSC, hemicellulose, total phenolics and soluble phenolics, and negatively to the C:N ratio and concentrations of lignin and cellulose.

*Soil fertility*

For each plot, soil properties were measured using standard protocols on forest floor (including both unfragmented aboveground litter and fragmented/humified organic matter, OL/OF/OH horizons) and mineral soil (A horizon, 10 cm depth) samples (n = 9 samples per plot) collected between spring and summer 2012 as described by Dawud et al. (2016)[42]. Briefly, the forest floors was sampled using a 25 cm by 25 cm wooden frame, while mineral soil was sampled with a soil corer (3.6 cm diameter). The nine samples were then homogenized and pooled at the plot level, dried at 55 °C to constant weight, and forest floor dry mass per unit surface area was measured. The pH was determined using a 827 pH lab sensor (Metrohm AG, Herisau, Switzerland) with 0.01 M $CaCl_2$ solution at a ratio of 1:10 and 1:2.5 for forest floor and mineral soil, respectively. Both forest floor and mineral soil were then ground, and their C and N contents were measured by dry combustion using a FLASH 2000 Soil CN Analyzer (Thermo Fisher Scientific, Milan, Italy). The soil fertility dataset had missing values for forest floor pH and C:N in one Finland plot. These missing values were replaced based on a regularized iterative multiple correspondence analysis[73] as described above.

We performed a PCA as described above to reduce the dimensionality of soil properties (including both forest floor and mineral soil) into a single dimension (PC1, 34.8% of variation, Supplementary Fig. 1b). This soil fertility axis ranged from low to high fertility, and was related positively to forest floor pH and lignin concentrations, as well as mineral soil pH, SOC, silt and clay contents, and negatively to forest floor C:N ratio, total phenolics and condensed tannins concentrations, as well as mineral soil C:N ratio.

**Measured ecosystem processes**

To characterize *in situ* patterns of litter decomposition in each plot, we used field-based data[79] of naturally occurring leaf litter decomposition. Briefly, litterbags were filled with 10 g of air-dried leaf litter consisting of a litter mixture with equal proportions of litter from all tree species present in the plot. Freshly senesced leaf litter from each tree species was collected at tree species-specific peak leaf litter fall between October 2011 and November 2012, in close vicinity of the experimental plots. Three replicate litterbags filled with the plot-specific leaf litter were placed on the bare soil, after the natural litter layer had been locally removed, within a 1 $m^2$ homogeneous area of each plot. Litterbags were retrieved when the most rapidly decomposing species within each region reached 40-50% mass loss. Collected decomposed materials were dried at 65 °C, cleaned of pieces of wood, stones or other foreign material that occasionally got into the litterbags and weighed. To correct for potential soil contamination during decomposition in the field, litter samples were ground with a Cyclotec Sample Mill (Tecator), their ash content determined and their mass loss rates expressed based on ash-free litter mass. To account for the differences in the duration of field exposure among the different locations, litter decomposition rate (mg $g^{-1}$ litter $day^{-1}$) was calculated as the ratio of mass lost per amount of initial mass per day of incubation. Litter decomposition per unit surface area (mg $m^{-2}$ $day^{-1}$) was calculated as the litter decomposition rates multiplied by the biomass of the litter layer.

Soil organic carbon (SOC) mineralization per unit surface area was estimated based on plot-specific soil microbial respiration and biomass data[29] decribed above to characterize SOM decomposition. SOC mineralization per unit surface area (mg C-$CO_2$ $m^{-2}$ $day^{-1}$) was calculated as the metabolic quotient of soil microbes standardized to the plot-specific soil temperature ($qCO_{2\ field}$, mg C-$CO_2$ $g^{-1}$ $C_{\text{microbial}}$ $day^{-1}$) and multiplied by the soil microbial biomass per unit surface area ($C_{microbial}$, g $C_{\text{microbial}}$ $m^{-2}$).

**Statistical analyses**

Bayesian linear mixed-effect models (LMMs) were fitted based on Markov Chain Monte Carlo (MCMC) methods using the '*rstanarm*' package[80]. We relied on the default settings of the '*rstanarm*' package for the priors, which are intended to be weakly informative, providing moderate regularization and helping stabilize computation (https://cran.r-project.org/web/packages/rstanarm/vignettes/priors.html#default-weakly-informative-prior-distributions). Five chains with different starting values were each ran over 2,000 iterations of both warm-up and sampling phases, resulting in a posterior sample size of 10,000. The target average acceptance probability ('adapt_delta') was set to 0.9999 to prevent divergent transitions and ensure reliable inference. We tested for model convergence by examining trace plots and evaluating the R-hat (*i.e.*, the ratio of between-chain variance to within-chain variance) and $N_{\text{eff}}$ (*i.e.,* the ratio of the effective sample size to the overall number of iterations) statistics. All models were checked for assumptions of normality of residuals, homogeneity of variances, linearity of relationships, and normality of random effects. We also performed graphical posterior prediction checks simulating replicated data under the fitted model and then comparing these to the observed data to ensure the lack discrepancies between real and simulated data[81,81]. For multiple regressions, multicollinearity issues were also checked based on variance inflation factors with a threshold value of 3, and collinear variables were omitted from the model when necessary[82]. All these model assumptions were graphically checked using the '*check_model*' function of the '*performance*' package[83]. To account for outliers and improve normality of the residuals, all response variables were log-transformed, except for 'soil engineering', 'detritivore biomass', and 'humi-detritivore biomass' which were log($x$+1)-transformed[84].

We used random intercept models (with varying intercepts but a common slope across locations) to explore general effects across European forests. The use of random slope models (with varying intercepts and slopes across locations for each predictor) instead of random intercept models has been recommended to control for inflated Type I (false positive) error[85,86], but such a confidence improvement comes at the cost of a significant loss of statistical power[87] (1 - Type II [false negative] error), *i.e.* a reduced ability to detect a particular effect that is actually present. To check that our inferences were robust to the approach employed, we therefore performed the same analyses using random slope models. Both approach provided very similar results, and the inclusion of random slopes never substantially improved model goodness-of-fit (Extended Table 2, Supplementary Table 14). To avoid model overfitting, we only present and discuss results from the simplest approach based on random intercept models in the main text.

To calculate regression slopes standardized by the standard deviation ($\beta_{st}$), raw slopes ($\beta_{unst}$) were scaled by the ratio of the standard deviation ($sd$) of the predictor variable ($x$) over the standard deviation of the response variable ($y$) using the following equation[88]:

$$\beta_{st} = \beta_{unst} \times \frac{sd_x}{sd_y} \tag{15}$$

Bayes factors (BFs) were calculated as the ratio of marginal likelihoods of two models: an alternative model ($m_1$) including the random factor tested over a null model ($m_0$) not including the random factor. A value of BF > 1 indicates that $m_1$ is more strongly supported by the data than $m_0$. The marginal likelihood of each model was calculated using the '*bridge_sampler*' function of the '*bridgesampling*' package[89]. We computed predicted values of the dependent variables for the average values of predictors and for different levels of the tree species richness predictor using the '*posterior_linpred*' function of the '*rstanarm*' package[80].

For each component LMM of the initial full SEM model (Supplementary Table 6), we performed stepwise backward model selection based on Pareto smoothed importance-sampling leave-one-out (PSIS-LOO) cross-validation[90] using the '*loo*' function of the '*loo*' package[91]. We fitted the full component LMM, computed the *p*-values associated with each fixed factor[92], and then removed the term with the highest *p*-value. To check that it improved model goodness-of-fit, the nested component LMMs were compared based on the LOO information criterion, *i.e.* the expected log predictive density (ELPD) multiplied by -2, with lower values indicating better model goodness-of-fit. We started by performing model selection on the component LMM predicting soil food web multifunctionality, and then performed selection on component LMMs predicting each of the variables retained as predictors of soil food web multifunctionality. To account for outliers in the SEM model, the variables 'tree aboveground biomass', 'tree aboveground litterfall', 'total root biomass' and 'soil food web multifunctionality' were log-transformed, while 'understory plant aboveground biomass (herbaceous)' was log($x$+1)-transformed.

The SEM model goodness-of-fit was assessed by the Shipley's d-sep test[93] combining the *p*-values of conditional independence claims in a test statistic, *i.e.* the Fisher's C, using the following equation:

$$C = -2\sum_{i=1}^{k} \ln(p_i) \quad (16)$$

where $p_i$ is the $i^{th}$ conditional independence claim in a basis set consisting of $k$ conditional independence claims associated with the hypothesized causal diagram. Fisher's C can then be compared to a $\chi^2$ distribution with $2k$ degrees of freedom. The hypothesized relationships of the SEM model are considered to be consistent with the data when there is weak support for the sum of the conditional independence claims, *i.e.* where the collection of such missing relationships represented by the Fisher's C could have easily occurred by chance, in which case the *p*-value for the $\chi^2$ test is greater than the chosen significance threshold (here $\alpha = 0.05$)[94]. For some conditional independence claims that were not verified (missing relationships unlikely to have occurred by chance), we fitted these relationships as correlated errors as these variables were assumed not to be causaly linked (*i.e.* to be caused by a shared underlying, unmeasured driver).

The full R script used for statistical analysis will be made available upon acceptance for publication.

### Supplementary Results

To test the theoretical assumption that the computed trophic functions based on aggregated energy fluxes through the soil food web provides a good description of independently measured ecosystem functions, we performed simple regressions or analysis of variance where a given measured ecosystem function was predicted based on Bayesian linear models including the corresponding calculated trophic function as fixed factor. Overall, we found that this theoretical assumption was well supported (Extended data Fig. 1). Indeed, the calculated decomposition of litter and soil organic matter (consumption of litter or soil organic matter by microbes) per unit surface area were positively related to measured litter mass loss and soil organic matter mineralization per unit surface area, respectively (Extended data Fig. 1a,e). Similarly, the calculated litter engineering rate (consumption of litter by fauna per unit litter mass) was positively related to measured litter decomposition rate (litter mass loss per unit litter mass, Extended data Fig. 1b), reflecting the stimulation of litter decomposition rate by soil faunal detritivory[95]. We also found that the calculated soil engineering (consumption of soil organic matter by fauna) per unit surface area was much higher for Mull/Amphimull humus types, characterized by the homogenization of humified organic matter with mineral particles within macro-aggregates, than for Moder/Mor humus types, characterized by the presence of humified organic matter (OH) separated from the mineral soil[96,97] (Extended data Fig. 1c). This shows that the calculated soil engineering is well representing the bioturbation of mineral soil by burrowing fauna such as endogeic and anecic earthworms. Accordingly, the calculated soil engineering per unit surface area was negatively related to the measured mass of forest floor (both unfragmented aboveground litter and fragmented/humified organic matter, OL/OF/OH, Extended data Fig. 1d), reflecting the rapid disappearance of litter and humified organic matter from the surface due to the bioturbation and translocation of dead organic matter from the surface to mineral soil by anecic earthworms. Altogether, these results provide empirical evidence for good agreement between calculated trophic functions and measured soil processes, confirming that our quantification of aggregated energy fluxes in soil food webs are reflective of soil functioning[98,99].

### Supplementary Figures and Tables

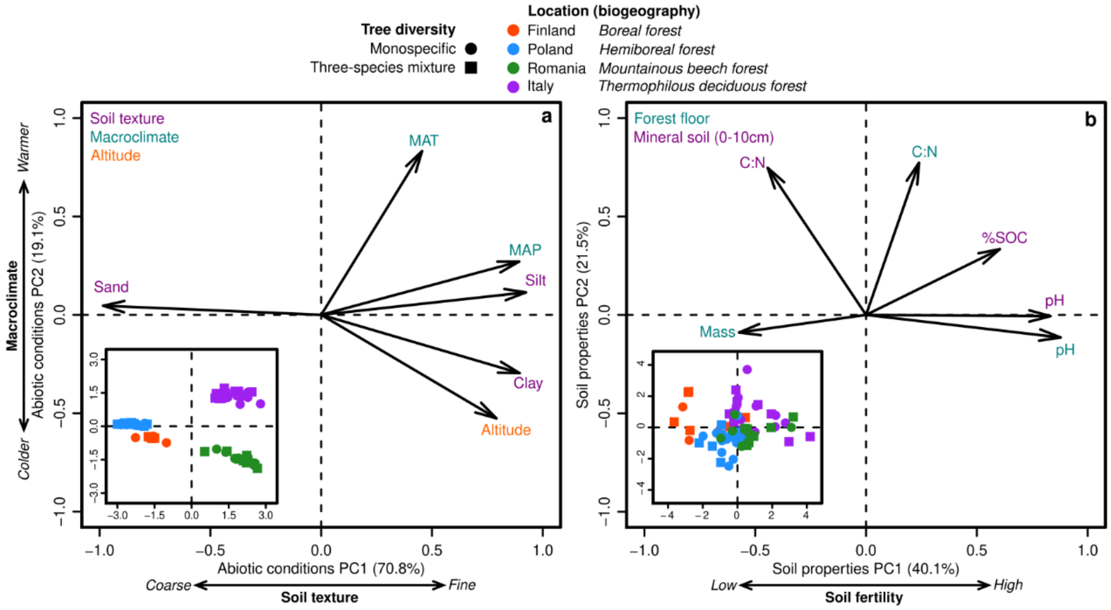

**Supplementary Fig. 1. Ordination by Principal Component Analysis (PCA) of abiotic conditions (a), and soil properties (b).** Variable loadings indicate the correlation of the variables with the corresponding principal components. MAT, mean annual temperature; MAP, mean annual precipitation; %SOC, soil organic carbon content.

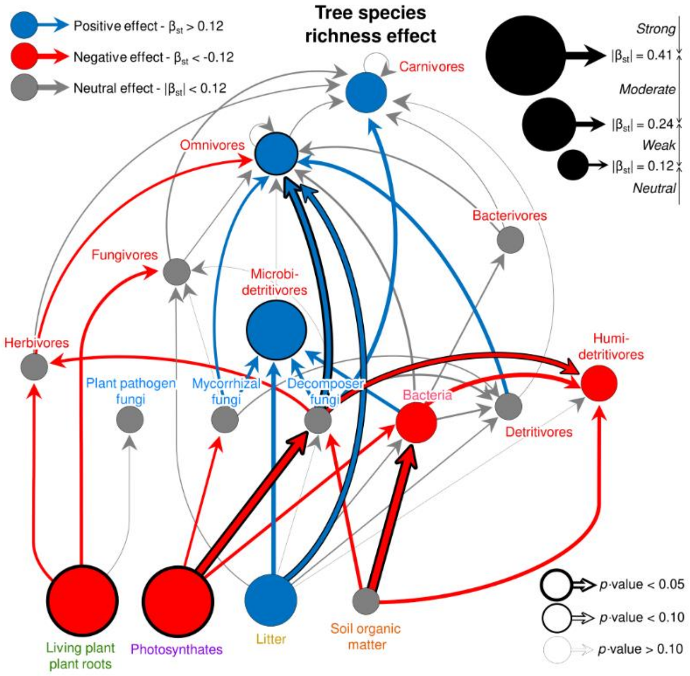

**Supplementary Fig. 2. Effects of tree species mixing on energy fluxes and trophic group biomasses of the soil food web across European forests.** Circle areas and arrows widths are scaled by the effect size of tree species richness, which was quantified using slope regression coefficients standardized by standard deviation ($\beta_{st}$) derived from Bayesian linear mixed-effect models used for the taxonomic approach (Methods).

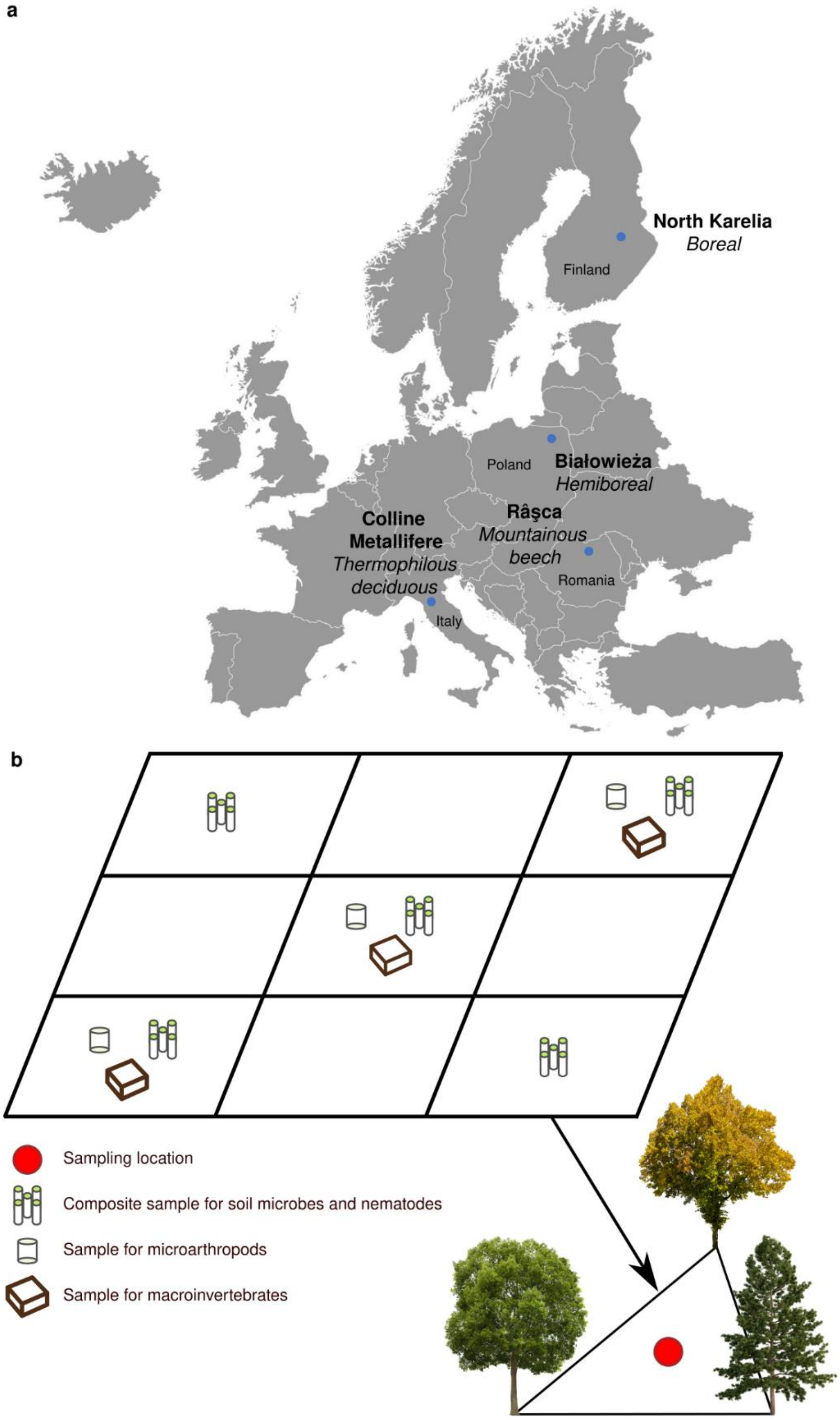

**Supplementary Fig. 3. Map of study site location across Europe (a), and illustration of the sampling design for each plot (b).**

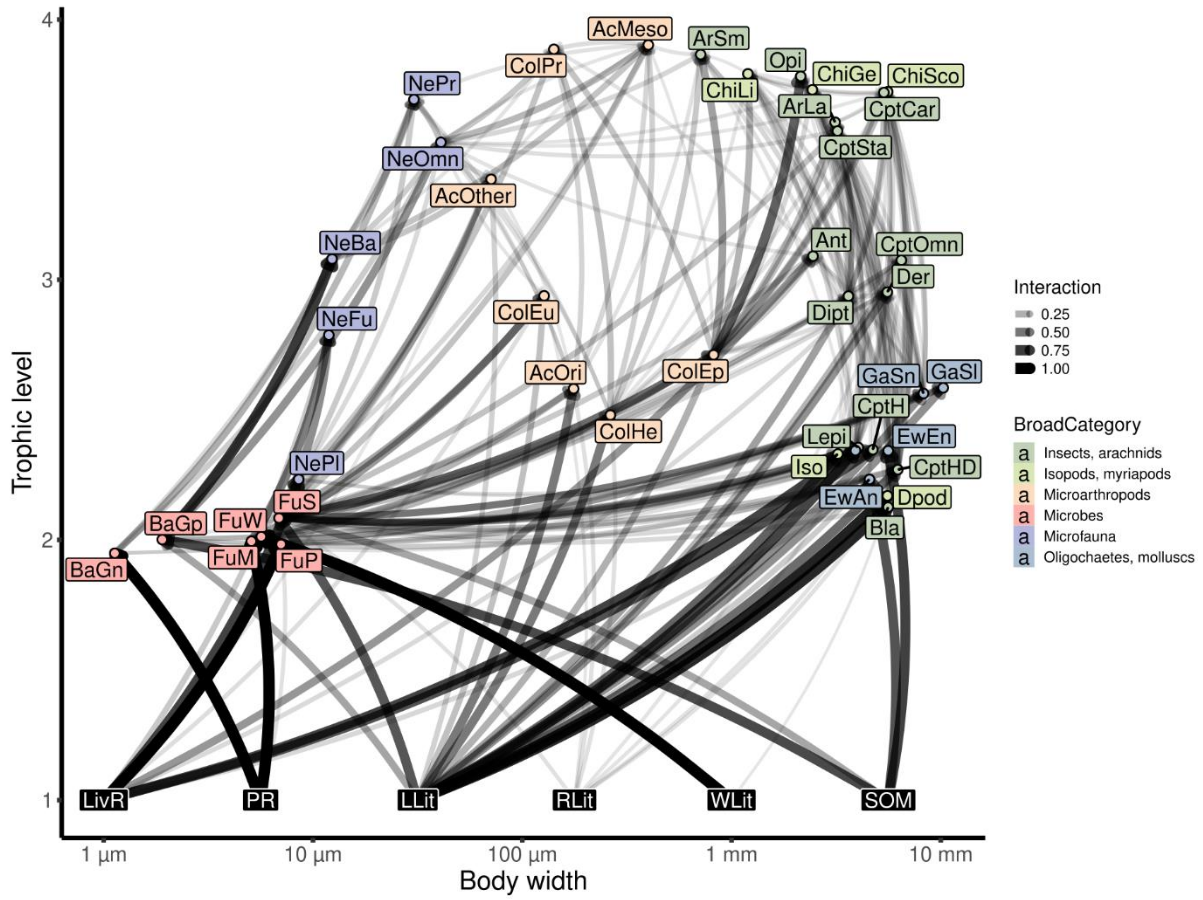

**Supplementary Fig. 4. Graphical illustration of the food web topology and trophic interaction strengths among trophic guilds for the studied European forest plots**. Arrows indicate energy flows among trophic groups, with arrow widths corresponding to trophic interaction strengths. The position of the trophic guilds are based on their body width and trophic level within the food web. The horizontal position of basal ressources at the food web bottom is arbitrary. Biomass data averaged across all plots was used to calculate trophic interaction strengths shown in this illustration. LivR, Living plant fine roots; PR, Photosynthates; LLit, Dead leaves; RLit, Dead roots; WLit, Dead wood; SOM, Soil organic matter; BaGn, Bacteria: Gram-; BaGp: Gram+; FuM, Mycorrhizal fungi; FuS, General saprotroph fungi; FuW, Wood saprotroph fungi; FuP, Plant pathogenic fungi; NePl, Nematoda: Plant feeders; NeBa, Nematoda: Bacterivores; NeFu, Nematoda: Fungivores; NeOmn, Nematoda: Omnivores; NePr, Nematoda: Predators; AcOri, Acari: Oribatida; AcMeso, Acari: Mesostigmata; AcOther, Acari: Prostigmata & Astigmata; ColEp, Collembola: Epedaphic; ColHe, Collembola: Hemiedaphic; ColEu, Collembola: Euedaphic; ColPr, Collembola: Predators; ArSm, Araneae: Small; ArLa, Araneae: Large; Opi, Opiliones; Dpod, Diplopoda; ChiGe, Chilopoda: Geophilomorpha; ChiSco, Chilopoda: Scolopendromorpha; ChiLi, Chilopoda: Lithobiomorpha; Iso, Isopoda; Der, Dermaptera; Bla, Blattodea; Ant, Formicidae; CptCar, Coleoptera: Carabidae; CptSta, Coleoptera: Staphylinidae; CptOmn, Coleoptera: Omnivores; CptHD, Coleoptera: Herbidetritivores; CptH, Coleoptera: Root feeders; Lepi, Lepidoptera; Dipt, Diptera; EwEp, Lumbricina: Epigeic; EwAn, Lumbricina: Anecic; EwEn, Lumbricina: Endogeic; GaSn, Gastropoda: Snails; GaSl, Gastropoda: Slugs. The interaction strength values shown here are provided in Supplementary Table 15.

**Supplementary Table 1**. **Study site characteristics**. Where applicable, mean ± standard error are given for each location. Mineral soil data are given for the 0-10 cm depth.

| | **North Karelia** | **Białowieża** | **Râşca** | **Colline Metallifere** |
|---|---|---|---|---|
| Country | Finland | Poland | Romania | Italy |
| Latitude/Longitude (°) | 62.9, 29.9 | 52.8, 23.9 | 47.3, 26.0 | 43.2, 11.2 |
| Elevation range (m) | 80–200 | 135–185 | 600–1,000 | 260–525 |
| Mean annual air temperature (°C) | 4.1 ± 0.1 | 9.8 ± 0.5 | 8.2 ± 0.5 | 12.5 ± 0.8 |
| Mean annual precipitation (mm) | 632 | 597 | 692 | 738 |
| Soil type | Podzsol | Cambisol/Luvisol | Eutric Cambisol | Cambisol |
| Soil texture class | Sandy loam | Sandy loam | Silty clay loam | Silt loam |
| Sand, silt, clay (%) | 48, 47, 5 | 65, 29, 6 | 13, 60, 27 | 17, 65, 18 |
| Soil bulk density (g $cm^{-3}$) | 1.03 ± 0.03 | 1.02 ± 0.02 | 0.93 ± 0.02 | 0.88 ± 0.02 |
| Mean annual soil temperature (°C) | 5.5 ± 0.2 | 9.4 ± 0.4 | 8.2 ± 0.5 | 12.2 ± 0.5 |
| Mean annual soil moisture (%) | 32.8 ± 1.7 | 24.6 ± 0.8 | 31.0 ± 2.1 | 21.3 ± 2.1 |
| Dominant humus type | Mull/Mor | Mull/Moder | Mull | Amphimull/Mull |
| Forest floor mass (kg $m^{-2}$) | 4.27 ± 0.81 | 2.28 ± 0.41 | 2.23 ± 0.39 | 1.81 ± 0.15 |
| Forest floor pH | 3.7 ± 0.2 | 4.8 ± 0.1 | 4.8 ± 0.1 | 5.4 ± 0.1 |
| Forest floor C:N | 27.5 ± 1.6 | 27.1 ± 0.9 | 30.5 ± 0.8 | 32.4 ± 0.9 |
| Mineral soil C (g $kg^{-1}$) | 37.8 ± 3.8 | 28.4 ± 1.2 | 49.2 ± 4.0 | 50.4 ± 3.1 |
| Mineral soil pH | 3.9 ± 0.1 | 3.8 ± 0.1 | 4.6 ± 0.2 | 4.6 ± 0.2 |
| Mineral soil C:N | 23.8 ± 1.3 | 16.7 ± 0.5 | 13.9 ± 0.3 | 19.9 ± 0.8 |
| Forest type | Boreal | Hemiboreal, nemoral coniferous, mixed broadleaved-coniferous | Mountainous mixed beech | Thermophilous deciduous |
| Stand age (year) | 45 ± 2 | 101 ± 7 | 86 ± 5 | 65 ± 2 |
| Basal area ($m^2$ $ha^{-1}$) | 28.4 ± 2.1 | 42.4 ± 2.4 | 51.0 ± 3.5 | 28.9 ± 0.8 |
| Tree aboveground biomass (kg $m^{-2}$) | 14.4 ± 1.0 | 28.7 ± 1.8 | 38.0 ± 2.4 | 21.4 ± 0.9 |
| Leaf area index ($m^2$ $m^{-2}$) | 2.91 ± 0.27 | 5.10 ± 0.19 | 5.66 ± 0.18 | 3.73 ± 0.23 |
| Target tree species | *Betula pendula/pubescens*⁋, *Picea abies*⁋, *Pinus sylvestris*⁋ | *Betula pendula*⁋, *Carpinus betulus*⁋, *Picea abies*⁋, *Pinus sylvestris*⁋, *Quercus robur*⁋ | *Abies alba*⁋, *Acer pseudoplatanus*†, *Fagus sylvatica*⁋, *Picea abies*⁋ | *Castanea sativa*, *Ostrya carpinifolia*, *Quercus cerris*, *Quercus ilex*, *Quercus petraea* |
| Number of monospecific/mixed stands | 6/3 | 6/14 | 8/8 | 10/9 |
| Soil sampling date (yyyy/mm/dd) | 2017/06/12-16 | 2017/05/05-10 | 2017/05/22-28 | 2017/04/10-16 |

⁋ Ectomycorrhizal tree species

† Arbuscular mycorrhizal tree species

**Supplementary Table 2**. **List of plant-derived (basal) resources and trophic guilds with trait values used for soil food web reconstruction**.

| | | | Feeding preferences | | | | | | | | | | | Body size | | Predator traits | | | Protection traits | | | | Vertical stratification | | |
|---|---|---|---|---|---|---|---|---|---|---|---|---|---|---|---|---|---|---|---|---|---|---|---|---|---|
| Trophic guild | Trophic group | Abbreviation | LivR | PR | LLit & RLit | WLit | SOM | Ba | FuM | FuS | FuW | FuP | Fauna | Mass mean | Mass SD | Trait | PPMR optimum | PPMR width | Carnivore | Agility | Physical protection | Metabolites | epi | hemi | eu |
| Living plant fine roots | Living plant fine roots | LivR | | | | | | | | | | | | | | | 1 | 1 | 1 | 1 | 1 | 1 | | 1 | 1 |
| Photosynthates & rhizodeposits | Photosynthates & rhizodeposits | PR | | | | | | | | | | | | | | | 1 | 1 | 1 | 1 | 1 | 1 | | 1 | 1 |
| Dead leaves | Plant litter | LLit | | | | | | | | | | | | | | | 1 | 1 | 1 | 1 | 1 | 1 | 1 | 1 | |
| Dead roots | Plant litter | RLit | | | | | | | | | | | | | | | 1 | 1 | 1 | 1 | 1 | 1 | | 1 | 1 |
| Dead wood | Plant litter | WLit | | | | | | | | | | | | | | | 1 | 1 | 1 | 1 | 1 | 1 | 1 | 1 | |
| Soil organic matter | Soil organic matter | SOM | | | | | | | | | | | | | | | 1 | 1 | 1 | 1 | 1 | 1 | | | 1 |
| Bacteria: Gram- | Bacteria | BaGn | | 1 | | | | | | | | | | | | | 1 | 1 | 1 | 1 | 1 | 1 | 1 | 1 | 1 |
| Bacteria: Gram+ & Actinomycetes | Bacteria | BaGp | | | 1 | | 1 | | | | | | | | | | 1 | 1 | 1 | 1 | 1 | 1 | | 1 | 1 |
| Mycorrhizal fungi | Mycorrhizal fungi | FuM | | 1 | | | | | | | | | | | | | 1 | 1 | 1 | 1 | 1 | 1 | | 1 | 1 |
| General saprotroph fungi | Decomposer fungi | FuS | | 1 | 1 | | 0.5 | | | | | | | | | | 1 | 1 | 1 | 1 | 1 | 1 | 1 | 1 | 1 |
| Wood saprotroph fungi | Decomposer fungi | FuW | | | | 1 | | | | | | | | | | | 1 | 1 | 1 | 1 | 1 | 1 | 1 | 1 | 1 |
| Plant pathogenic fungi | Plant pathogenic fungi | FuP | 1 | | | | | | | | | | | | | | 1 | 1 | 1 | 1 | 1 | 1 | 1 | 1 | 1 |
| Nematoda: Plant feeders | Herbivores | NePl | 1 | | | | | | | 0.5 | | 0.5 | | -6.62 | 0.5 | | 1 | 1 | 1 | 1 | 1 | 1 | | 1 | 1 |
| Nematoda: Bacterivores | Bacterivores | NeBa | | | | | | 1 | | | | | | -6.14 | 0.5 | | 1 | 1 | 1 | 1 | 1 | 1 | | 1 | 1 |
| Nematoda: Fungivores | Fungivores | NeFu | 0.5 | | | | | | 0.5 | 1 | | | | -6.44 | 0.5 | | 1 | 1 | 1 | 1 | 1 | 1 | | 1 | 1 |
| Nematoda: Omnivores | Omnivores | NeOmn | | | | | | 1 | | | | | 1 | -5.05 | 1 | | 1 | 1 | 1 | 1 | 1 | 1 | | 1 | 1 |
| Nematoda: Predators | Carnivores | NePr | | | | | | 0.5 | | | | | 1 | -5.47 | 1 | | 1 | 1 | 0.7 | 1 | 1 | 1 | | 1 | 1 |
| Acari: Oribatida | Microbi-detritivores | AcOri | | | 1 | | | 0.5 | 0.5 | 1 | | | 0.5 | -4.91 | 0.6 | | 1 | 1 | 1 | 1 | 0.4 | 0.7 | | 1 | 0.5 |
| Acari: Mesostigmata | Carnivores | AcMeso | | | | | | | | 0.5 | | | 1 | -4.75 | 1 | | 1 | 1 | 0.7 | 1 | 0.7 | 1 | | 1 | 0.5 |
| Acari: Prostigmata & Astigmata | Omnivores | AcOther | 0.5 | | 0.5 | | | 0.5 | 0.5 | 1 | | | 1 | -5.63 | 0.6 | | 1 | 1 | 0.7 | 1 | 0.7 | 1 | | 1 | 0.5 |
| Collembola: Epedaphic | Fungivores | ColEp | | | 0.5 | | | | 0.5 | 1 | | | | -3.16 | 0.5 | | 1 | 1 | 1 | 0.7 | 1 | 1 | 1 | 0.5 | |
| Collembola: Hemiedaphic | Microbi-detritivores | ColHe | | | 1 | | | 0.5 | 0.5 | 1 | | | 0.5 | -4.29 | 0.6 | | 1 | 1 | 1 | 0.7 | 1 | 1 | | 1 | 0.5 |
| Collembola: Euedaphic | Fungivores | ColEu | 0.5 | | 0.5 | | | 0.5 | 0.5 | 1 | | | 0.5 | -4.54 | 0.6 | | 1 | 1 | 1 | 1 | 1 | 0.4 | | 1 | 1 |
| Collembola: Predators | Carnivores | ColPr | | | | | | | | 0.5 | | | 1 | -3.74 | 0.6 | | 1 | 1 | 1 | 1 | 1 | 0.4 | 1 | 1 | |

| | | | Feeding preferences | | | | | | | | | | | Body size | | Predator traits | | | Protection traits | | | | Vertical stratification | | |
|---|---|---|---|---|---|---|---|---|---|---|---|---|---|---|---|---|---|---|---|---|---|---|---|---|---|
| Trophic guild | Trophic group | Abbreviation | LivR | PR | LLit & RLit | WLit | S | Ba | FuM | FuS | FuW | FuP | Fauna | Mass mean | Mass SD | Trait | PPMR optimum | PPMR width | Carnivore | Agility | Physical protection | Metabolites | epi | hemi | eu |
| Araneae: Small | Carnivores | ArSm | | | | | | | | | | | 1 | -3.26 | 0.5 | venom and web | 0.64 | 1.44 | 0.7 | 1 | 1 | 1 | 0.5 | 1 | |
| Araneae: Large | Carnivores | ArLa | | | | | | | | | | | 1 | -2.06 | 1 | | 0.64 | 1.44 | 0.7 | 1 | 1 | 1 | 1 | 0.5 | |
| Opiliones | Carnivores | Opi | | | | | | | | | | | 1 | -2.31 | 0.5 | | 1 | 1 | 0.7 | 1 | 1 | 1 | 1 | | |
| Diplopoda | Detritivores | Dpod | 0.5 | | 1 | 0.5 | | | | 0.5 | | | | -1.33 | 0.6 | | 1 | 1 | 1 | 1 | 0.4 | 0.7 | 1 | 1 | 0.5 |
| Chilopoda: Geophilomorpha | Carnivores | ChiGe | | | | | | | | | | | 1 | -1.96 | 0.60 | venom | 0.8 | 1.2 | 0.7 | 1 | 1 | 1 | | 1 | 1 |
| Chilopoda: Scolopendromorpha | Carnivores | ChiSco | | | | | | | | | | | 1 | -2.23 | 0.60 | | 0.8 | 1.2 | 0.7 | 1 | 1 | 1 | 0.5 | 1 | 0.5 |
| Chilopoda: Lithobiomorpha | Carnivores | ChiLi | | | | | | | | | | | 1 | -2.16 | 0.60 | | 0.8 | 1.2 | 0.7 | 1 | 1 | 1 | 0.5 | 1 | |
| Isopoda | Detritivores | Iso | 0.5 | | 1 | 0.5 | | 0.5 | 0.5 | 1 | | | | -1.70 | 1 | | 1 | 1 | 1 | 1 | 0.4 | 1 | 1 | 0.5 | |
| Dermaptera | Omnivores | Der | 0.5 | | 1 | | | 0.5 | | 0.5 | | | 1 | -1.42 | 1 | | 1 | 1 | 0.7 | 1 | 1 | 0.4 | 1 | 1 | |
| Blattodea | Detritivores | Bla | 0.5 | | 1 | 0.5 | | | | 0.5 | | | 0.5 | -1.99 | 1 | | 1 | 1 | 1 | 1 | 0.7 | 0.7 | 1 | 0.5 | |
| Formicidae | Omnivores | Ant | 1 | | | | | | | 0.5 | | 0.5 | 1 | -3.30 | 0.6 | pack hunting | 0.6 | 1.6 | 0.7 | 1 | 1 | 0.7 | 1 | 0.5 | 0.5 |
| Coleoptera: Carabidae | Carnivores | CptCar | | | | | | | | | | | 1 | -1.60 | 1 | | 1 | 1 | 0.7 | 1 | 0.4 | 0.4 | 1 | | |
| Coleoptera: Staphylinidae | Carnivores | CptSta | | | | | | | | 0.5 | | | 1 | -2.53 | 1 | | 1 | 1 | 0.7 | 1 | 0.4 | 0.4 | 1 | 1 | 1 |
| Coleoptera: Elateridae (Omnivores) | Omnivores | CptOmn | 1 | | 1 | 1 | | 0.5 | 0.5 | 1 | 0.5 | 0.5 | 1 | -1.93 | 1 | | 1 | 1 | 1 | 1 | 0.4 | 1 | | 1 | 1 |
| Coleoptera: Scarabaeidae / Geotrupidae (Herbi-detritivores) | Detritivores | CptHD | 1 | | 1 | 1 | | 0.5 | 0.5 | 1 | 0.5 | 0.5 | | -0.51 | 1 | | 1 | 1 | 1 | 1 | 0.4 | 1 | 1 | 1 | 1 |
| Coleoptera: Curculionidae (Root feeders) | Herbivores | CptH | 1 | | | 0.5 | | | | 0.5 | 0.5 | 0.5 | | -1.72 | 1 | | 1 | 1 | 1 | 1 | 0.4 | 1 | | 1 | 1 |
| Lepidoptera | Herbivores | Lepi | 1 | | | | | | | 0.5 | | 0.5 | | -1.55 | 1 | | 1 | 1 | 1 | 1 | 1 | 1 | 1 | 1 | |
| Diptera | Omnivores | Dipt | 0.5 | | 1 | 0.5 | | | 0.5 | 1 | | | 1 | -1.44 | 1 | | 1 | 1 | 1 | 1 | 1 | 1 | 1 | 1 | |
| Lumbricina: Epigeic | Detritivores | EwEp | 0.5 | | 1 | 0.5 | | 0.5 | | 0.5 | | | 0.5 | -0.78 | 0.6 | | 4.5 | 0.75 | 1 | 1 | 1 | 0.4 | 0.5 | 1 | |
| Lumbricina: Anecic | Humi-detritivores | EwAn | 0.5 | | 1 | | 1 | 0.5 | | 0.5 | | | 0.5 | -0.56 | 0.7 | filtering the environment | 4.5 | 0.75 | 1 | 1 | 1 | 0.4 | | 1 | 1 |
| Lumbricina: Endogeic | Humi-detritivores | EwEn | 0.5 | | | | 1 | 0.5 | | 0.5 | | | 0.5 | -0.65 | 0.7 | | 4.5 | 0.75 | 1 | 1 | 1 | 0.4 | | | 1 |
| Gastropoda: Snails | Detritivores | GaSn | 0.5 | | 1 | 0.5 | | 1 | 0.5 | 1 | | | 0.5 | -1.51 | 1 | | 1 | 1 | 1 | 1 | 0.4 | 0.4 | 1 | 1 | |
| Gastropoda: Slugs | Detritivores | GaSl | 0.5 | | 1 | 0.5 | | 1 | 0.5 | 1 | | | 0.5 | -1.01 | 0.6 | | 1 | 1 | 1 | 1 | 0.7 | 0.4 | 1 | 1 | 0.5 |

Herbivores, fauna mostly feeding on living plant fine roots; detritivores, fauna mostly feeding on litter; humi-detritivores, fauna feeding on both litter and soil organic matter; microbi-detritivores, fauna feeding on both microbes and litter; bacterivores, fauna mostly feeding on bacteria; fungivores, fauna mostly feeding on fungi; omnivores; fauna feeding on both fauna and other resources; carnivores, fauna mostly feeding on fauna.

**Supplementary Table 3. Results of Bayesian multi-level models testing tree community effects on soil food web multifunctionality using both the ‘average’ and ‘threshold’ approaches.**

| | Taxonomic approach for tree community effects | | | | | | Functional approach for tree community effects | | | | | | | | |
|---|---|---|---|---|---|---|---|---|---|---|---|---|---|---|---|
| | Species richness | | | Species composition | | Model $r^2$ | Functional diversity | | | Functional composition | | | | | Model $r^2$ |
| | | | | | | | | | | | Leaf economics spectrum | | Fine-root economics spectrum | | |
| | % var | $\beta_{st}$ [CI 95%] | *p*-value | % var | BF | | % var | $\beta_{st}$ [CI 95%] | *p*-value | % var | $\beta_{st}$ [CI 95%] | *p*-value | $\beta_{st}$ [CI 95%] | *p*-value | |
| Average | 4.43 | -0.19 [-0.46;0.10] | 0.192 | 22.26 | **4.24** | 0.47 | 1.57 | -0.13 [-0.35;0.10] | 0.254 | 36.67 | 0.49 [0.20;0.80] | **0.002**** | 0.68 [0.20;1.17] | **0.009**** | 0.38 |
| 10% threshold | 4.97 | -0.20 [-0.47;0.07] | 0.148 | 10.48 | 0.64 | 0.32 | 3.88 | -0.20 [-0.44;0.03] | 0.087† | 15.84 | 0.31 [0.00;0.64] | 0.053† | 0.26 [-0.27;0.79] | 0.306 | 0.30 |
| 20% threshold | 4.81 | -0.19 [-0.50;0.11] | 0.211 | 29.23 | **8.64** | 0.47 | 1.81 | -0.13 [-0.37;0.11] | 0.281 | 27.57 | 0.35 [0.02;0.72] | **0.036*** | 0.62 [0.10;1.25] | **0.022*** | 0.28 |
| 30% threshold | 1.87 | -0.07 [-0.34;0.19] | 0.558 | 9.27 | 0.71 | 0.39 | 0.66 | -0.00 [-0.22;0.22] | 0.989 | 37.30 | 0.47 [0.15;0.75] | **0.008**** | 0.72 [0.21;1.21] | **0.008**** | 0.41 |
| 40% threshold | 4.10 | -0.18 [-0.45;0.08] | 0.169 | 6.62 | 0.48 | 0.29 | 1.93 | -0.14 [-0.37;0.09] | 0.231 | 29.75 | 0.45 [0.11;0.76] | **0.010*** | 0.54 [0.00;1.03] | 0.050† | 0.33 |
| 50% threshold | 8.59 | -0.29 [-0.56;-0.02] | **0.037*** | 10.72 | 0.70 | 0.32 | 4.78 | -0.26 [-0.50;-0.03] | **0.025*** | 28.98 | 0.48 [0.18;0.80] | **0.003**** | 0.52 [0.01;1.01] | **0.044*** | 0.33 |
| 60% threshold | 2.08 | -0.07 [-0.35;0.22] | 0.647 | 9.00 | 0.48 | 0.18 | 1.65 | -0.08 [-0.34;0.18] | 0.527 | 14.16 | 0.21 [-0.13;0.58] | 0.220 | 0.32 [-0.24;0.90] | 0.243 | 0.15 |
| 70% threshold | 1.70 | 0.02 [-0.26;0.29] | 0.906 | 9.22 | 0.56 | 0.24 | 1.37 | 0.05 [-0.21;0.30] | 0.709 | 10.38 | 0.17 [-0.18;0.50] | 0.310 | 0.16 [-0.39;0.68] | 0.512 | 0.20 |
| 80% threshold | 2.60 | 0.10 [-0.19;0.38] | 0.505 | 12.88 | 0.69 | 0.27 | 1.46 | 0.06 [-0.20;0.31] | 0.650 | 16.67 | 0.27 [-0.10;0.60] | 0.139 | 0.28 [-0.33;0.82] | 0.313 | 0.22 |
| 90% threshold | 3.68 | 0.14 [-0.15;0.43] | 0.317 | 20.8 | **1.44** | 0.39 | 1.65 | 0.08 [-0.16;0.32] | 0.497 | 22.10 | 0.34 [0.02;0.66] | **0.040*** | 0.35 [-0.17;0.84] | 0.166 | 0.28 |

Both the leaf and fine-root economics spectra variables representing tree functional composition ranged from slow/conservative to fast/acquisitive economic strategies (two first PCA axes of CWM tree traits, Extended Data Fig. 2a). % var, proportion of total variance. $\beta_{st}$, slope regression coefficients standardised by standard deviation. CI 95%, 95% credible intervals. BF, Bayes factor calculated as the ratio of marginal likelihoods of two models: an alternative model ($m_1$) including the random factor tested over a null model ($m_0$) not including the random factor. Values of BF > 1 and > 10 respectively indicates moderate and strong support in favour of the random factor effect. Significant effects ($p < 0.05$ or BF > 1) are reported in bold. †, $p < 0.10$; *, $p < 0.050$; **, $p < 0.010$; ***, $p < 0.001$.

**Supplementary Table 4. Functional trait values of tree species for each location.** LNC, leaf nitrogen content; SLA, specific leaf area; LDMC, leaf dry matter content; RNC, root nitrogen content; SRL, specific root length; RTD, root tissue density; $D_m$, mean root diameter; RLD, root length density;%Myc, ectomycorrhizal colonisation intensity; ECM, Ectomycorrhizal tree species; AM, Arbuscular mycorrhizal tree species.

| Country | Species | Mycorrhizal type | Leaf phenology | LNC (% dry mass) | SLA ($mm^2$ $mg^{-1}$) | LDMC (mg $g^{-1}$) | RNC (% dry mass) | SRL (m $g^{-1}$) | RTD (g $cm^{-3}$) | $D_m$ (mm) | RLD (cm $cm^{-3}$) | %Myc (number $cm^{-1}$) |
|---|---|---|---|---|---|---|---|---|---|---|---|---|
| Finland | *Betula pendula/pubescens* | ECM | Deciduous | 2.37 | 14.6 | 385 | 1.21 | 42.7 | 0.333 | 0.321 | 0.88 | 2.3 |
| | *Picea abies* | ECM | Evergreen | 1.10 | 3.2 | 431 | 1.09 | 16.7 | 0.362 | 0.469 | 1.23 | 2.3 |
| | *Pinus sylvestris* | ECM | Evergreen | 1.49 | 3.6 | 422 | 1.07 | 22.8 | 0.387 | 0.394 | 1.16 | 1.9 |
| Poland | *Betula pendula* | ECM | Deciduous | 2.56 | 14.6 | 385 | 1.96 | 65.4 | 0.282 | 0.303 | 0.97 | 3.2 |
| | *Carpinus betulus* | ECM | Deciduous | 2.74 | 11.6 | 259 | 1.93 | 52.6 | 0.333 | 0.291 | 1.08 | 3.5 |
| | *Picea abies* | ECM | Evergreen | 1.30 | 3.7 | 447 | 1.62 | 25.7 | 0.311 | 0.433 | 0.39 | 4.6 |
| | *Pinus sylvestris* | ECM | Evergreen | 2.03 | 3.6 | 422 | 1.81 | 30.2 | 0.300 | 0.409 | 0.54 | 3.3 |
| | *Quercus robur* | ECM | Deciduous | 2.50 | 14.3 | 277 | 1.96 | 50.7 | 0.297 | 0.314 | 0.72 | 4.0 |
| Romania | *Abies alba* | ECM | Evergreen | 1.28 | 4.6 | 494 | 1.36 | 11.4 | 0.356 | 0.583 | 0.37 | 2.4 |
| | *Acer pseudoplatanus* | ECM | Deciduous | 2.16 | 12.5 | 366 | 1.53 | 35.2 | 0.391 | 0.355 | 1.41 | 0.0 |
| | *Fagus sylvatica* | ECM | Deciduous | 2.11 | 19.3 | 449 | 1.46 | 34.1 | 0.373 | 0.324 | 1.45 | 3.4 |
| | *Picea abies* | ECM | Evergreen | 1.31 | 4.2 | 463 | 1.29 | 12.3 | 0.438 | 0.542 | 0.65 | 2.8 |
| Italy | *Castanea sativa* | ECM | Deciduous | 2.62 | 13.6 | 265 | 1.01 | 27.7 | 0.454 | 0.337 | 0.37 | 2.9 |
| | *Ostrya carpinifolia* | AM | Deciduous | 2.32 | 13.7 | 242 | 1.26 | 40.2 | 0.407 | 0.301 | 0.76 | 1.8 |
| | *Quercus cerris* | ECM | Deciduous | 2.26 | 17.9 | 306 | 0.92 | 20.8 | 0.509 | 0.354 | 0.94 | 1.9 |
| | *Quercus ilex* | ECM | Evergreen | 1.32 | 6.4 | 530 | 0.89 | 12.8 | 0.553 | 0.446 | 0.35 | 2.7 |
| | *Quercus petraea* | ECM | Deciduous | 2.19 | 12.5 | 281 | 1.06 | 22.5 | 0.475 | 0.359 | 2.09 | 2.2 |

**Supplementary Table 5. Principal component analysis of tree community-weighted mean (CWM) traits.** For each tree CWM trait, Pearson's r with the highest absolute value among the two PCA axes is shown in bold.

| Tree CWM trait | Correlation with principal components (Pearson's r) | | Contribution to principal components (%) | |
|---|---|---|---|---|
| | PC1 (45.25% of variance) | PC2 (31.48% of variance) | PC1 (45.25% of variance) | PC2 (31.48% of variance) |
| **Leaf traits** | | | | |
| Leaf nitrogen content (LNC) | **0.94** | -0.18 | 21.7 | 1.1 |
| Specific leaf area (SLA) | **0.63** | -0.59 | 9.8 | 12.5 |
| Leaf dry matter content (LDMC) | **-0.78** | 0.35 | 15.0 | 4.4 |
| **Fine-root traits** | | | | |
| Root nitrogen content (RNC) | 0.53 | **0.77** | 6.9 | 20.8 |
| Root tissue density (RTD) | -0.37 | **-0.81** | 3.5 | 23.0 |
| Specific root length (SRL) | **0.91** | 0.32 | 20.4 | 3.7 |
| Root diameter (Dm) | **-0.91** | 0.26 | 20.2 | 2.3 |
| Root length density (RLD) | 0.22 | **-0.59** | 1.2 | 12.2 |
| Root ectomycorrhizal colonization intensity (%Myc) | 0.24 | **0.75** | 1.4 | 20.0 |

**Supplementary Table 6. Initial full SEM model, with component linear mixed-effect models in each row.** abiotic.PC1, abiotic conditions (PC1); abiotic.PC2, abiotic conditions (PC2), SR, tree species richness; compo, tree species composition; fundiv, tree functional diversity; composition, tree species composition; LES, tree functional composition (leaf economics spectrum); RES, tree functional composition (root economics spectrum); AGB, tree aboveground biomass; LAI, leaf area index; litterfall, tree aboveground litterfall; litter.quality, tree leaf litter quality (PC1); litter.quality.diversity, functional diversity of tree leaf litter; root, total root biomass; woody.root, woody fine-root biomass; herbaceous.root, herbaceous fine-root biomass; under, total understorey aboveground plant biomass; woody.under, woody understorey aboveground plant biomass; herbaceous.under, herbaceous understorey aboveground plant biomass; CN.under, C:N ratio of understorey aboveground plant biomass; SR.under, species richness of understorey aboveground plants; microclim, microclimate (PC1); fertility, soil fertility (PC1); litter.mass, mass of dead leaves, roots and wood; SOM.mass, mass of soil organic matter; multifun, soil food web multifunctionality.

| Model formula |
|---|
| fundiv ~ SR + abiotic.PC1 + abiotic.PC2 + (1\|location) |
| LES ~ abiotic.PC1 + abiotic.PC2 + (1\|compo) + (1\|location) |
| RES ~ abiotic.PC1 + abiotic.PC2 + (1\|compo) + (1\|location) |
| ABG ~ fundiv + LES + RES + abiotic.PC1 + abiotic.PC2 + (1\|location) |
| LAI ~ fundiv + LES + RES + AGB + abiotic.PC1 + abiotic.PC2 + (1\|location) |
| litterfall ~ fundiv + LES + RES + AGB + LAI + abiotic.PC1 + abiotic.PC2 + (1\|location) |
| litter.quality (PC1) ~ fundiv + LES + RES + abiotic.PC1 + abiotic.PC2 + (1\|location) |
| litter.quality.diversity ~ fundiv + LES + RES + abiotic.PC1 + abiotic.PC2 + (1\|location) |
| root ~ fundiv + LES + RES + AGB + under + abiotic.PC1 + abiotic.PC2 + (1\|location) |
| woody.root ~ fundiv + LES + RES + AGB + woody.under + abiotic.PC1 + abiotic.PC2 + (1\|location) |
| herbaceous.root ~ fundiv + LES + RES + AGB + herbaceous.under + abiotic.PC1 + abiotic.PC2 + (1\|location) |
| under ~ fundiv + LES + RES + AGB + LAI + root + litterfall + litter.quality (PC1) + abiotic.PC1 + abiotic.PC2 + (1\|location) |
| woody.under ~ fundiv + LES + RES + AGB + LAI + root + litterfall + litter.quality (PC1) + abiotic.PC1 + abiotic.PC2 + (1\|location) |
| herbaceous.under ~ fundiv + LES + RES + AGB + LAI + root + litterfall + litter.quality (PC1) + abiotic.PC1 + abiotic.PC2 + (1\|location) |
| CN.under ~ fundiv + LES + RES + AGB + LAI + root + litterfall + litter.quality (PC1) + abiotic.PC1 + abiotic.PC2 + (1\|location) |
| SR.under ~ fundiv + LES + RES + AGB + LAI + root + litterfall + litter.quality (PC1) + abiotic.PC1 + abiotic.PC2 + (1\|location) |
| microclim ~ fundiv + LES + RES + AGB + LAI + under + litter.mass + SOM.mass + abiotic.PC1 + abiotic.PC2 + (1\|location) |
| fertility ~ fundiv + LES + RES + AGB + litterfall + litter.quality (PC1) + root + under + CN.under + abiotic.PC1 + abiotic.PC2 + (1\|location) |
| litter.mass ~ fundiv + LES + RES + AGB + litterfall + litter.quality (PC1) + root + under + CN.under + abiotic.PC1 + abiotic.PC2 + (1\|location) |
| SOM.mass ~ fundiv + LES + RES + AGB + litterfall + litter.quality (PC1) + root + under + CN.under + abiotic.PC1 + abiotic.PC2 + (1\|location) |
| multifun ~ fundiv + LES + RES + AGB + LAI + litterfall + litter.quality (PC1) + litter.quality.diversity + root + woody.root + herbaceous.root + under + woody.under + herbaceous.under + CN.under + SR.under + microclim + fertility + litter.mass + SOM.mass + abiotic.PC1 + abiotic.PC2 + (1\|location) |

**Supplementary Table 7. Basis set of models testing all conditional independence claims implied by the final SEM model.** abiotic.PC1, abiotic conditions (PC1); abiotic.PC2, abiotic conditions (PC2), SR, tree species richness; compo, tree species composition ;fundiv, tree functional diversity; LES, tree functional composition (leaf economics spectrum); RES, tree functional composition (fine-root economics spectrum); AGB, tree aboveground biomass; litterfall, tree aboveground litterfall; litter.quality, tree leaf litter quality (PC1); root, total root biomass; microclim, microclimate (PC1); herbaceous, understorey plant aboveground biomass (herbaceous); multifun, soil food web multifunctionality.

| Conditional independance claim | Model formula | $H_0$ | *p*-value |
|---|---|---|---|
| (SR, AGB) \| {fundiv, RES} | ABG~SR+fundiv+RES+(1\|location) | SR = 0 | 0.721 |
| (SR, litter.quality) \| {LES} | litter.quality~SR+LES+(1\|location) | SR = 0 | 0.619 |
| (SR, root) \| {fundiv, AGB} | root~SR+fundiv+AGB+(1\|location) | SR = 0 | 0.584 |
| (SR, microclim) \| {fundiv, LES, RES, abiotic.PC1, abiotic.PC2} | microclim~SR+fundiv+LES+RES+abiotic.PC1+abiotic.PC2+(1\|location) | SR = 0 | 0.572 |
| (SR, herbaceous) \| {AGB, root} | herbaceous~SR+AGB+root+(1\|location) | SR = 0 | 0.736 |
| (SR, multifun) \| {AGB, litterfall, litter.quality, root, herbaceous, microclim} | multifun~SR+AGB+litterfall+litter.quality+root +herbaceous+microclim+(1\|location) | SR = 0 | 0.989 |
| (LES, fundiv) \| {SR} | fundiv~LES+SR+(1\|location) | LES = 0 | 0.123 |
| (LES, AGB) \| {fundiv, RES} | AGB~LES+fundiv+RES+(1\|location) | LES = 0 | 0.561 |
| (LES, litterfall) \| {fundiv, AGB} | litterfall~LES+fundiv +AGB+(1\|location) | LES = 0 | 0.091 |
| (LES, root) \| {fundiv, AGB} | root~LES +fundiv+AGB+(1\|location) | LES = 0 | 0.338 |
| (LES, herbaceous) \| {AGB, root} | herbaceous~LES+AGB+root+(1\|location) | LES = 0 | 0.634 |
| (LES, multifun) \| {AGB, litterfall, litter.quality, root, herbaceous, microclim} | multifun~LES+AGB+litterfall+litter.quality+root +herbaceous+microclim+(1\|location) | LES = 0 | 0.745 |
| (RES, fundiv) \| {SR} | fundiv ~RES+SR+(1\|location) | RES = 0 | 0.133 |
| (RES, litterfall) \| {fundiv, AGB} | litterfall~RES+fundiv +AGB+(1\|location) | RES = 0 | 0.103 |
| (RES, litter.quality) \| {LES} | litter.quality~RES+LES+(1\|location) | RES = 0 | 0.540 |
| (RES, root) \| {fundiv, AGB} | root~RES+fundiv+AGB+(1\|location) | RES = 0 | 0.267 |
| (RES, herbaceous) \| {AGB, root} | herbaceous~RES+AGB+root+(1\|location) | RES = 0 | 0.829 |
| (RES, multifun) \| {AGB, litterfall, litter.quality, root, herbaceous, microclim} | multifun~RES+AGB+litterfall+litter.quality+root +herbaceous+microclim+(1\|location) | RES = 0 | 0.323 |
| (abiotic.PC1, fundiv) \| {SR} | fundiv~abiotic.PC1+SR+(1\|location) | abiotic.PC1 = 0 | 0.751 |
| (abiotic.PC1, LES) \| {compo} | LES~abiotic.PC1+(1\|compo)+(1\|location) | abiotic.PC1 = 0 | 0.764 |
| (abiotic.PC1, RES) \| {compo} | RES~abiotic.PC1+(1\|compo)+(1\|location) | abiotic.PC1 = 0 | 0.917 |
| (abiotic.PC1, ABG) \| {fundiv, RES} | AGB~abiotic.PC1+fundiv+RES+(1\|location) | abiotic.PC1 = 0 | 0.123 |
| (abiotic.PC1, litterfall) \| {fundiv, AGB} | litterfall~abiotic.PC1+fundiv+AGB+(1\|location) | abiotic.PC1 = 0 | 0.633 |
| (abiotic.PC1, litter.quality) \| {fundiv, LES} | litter.quality~abiotic.PC1+LES+(1\|location) | abiotic.PC1 = 0 | 0.651 |
| (abiotic.PC1, root) \| {fundiv, AGB} | root~abiotic.PC1+fundiv+AGB+(1\|location) | abiotic.PC1 = 0 | 0.339 |
| (abiotic.PC1, herbaceous) \| {AGB, root} | herbaceous ~abiotic.PC1+AGB+root+(1\|location) | abiotic.PC1 = 0 | 0.253 |
| (abiotic.PC1, multifun) \| {AGB, litterfall, litter.quality, root, herbaceous, microclim} | multifun~ abiotic.PC1+AGB+litterfall+litter.quality+root +herbaceous+microclim+(1\|location) | abiotic.PC1 = 0 | 0.761 |
| (abiotic.PC2, fundiv) \| {SR} | fundiv~abiotic.PC2+SR+(1\|location) | abiotic.PC2 = 0 | 0.993 |
| (abiotic.PC2, LES) \| {compo} | LES~abiotic.PC2+(1\|compo)+(1\|location) | abiotic.PC2 = 0 | 0.230 |
| (abiotic.PC2, RES) \| {compo} | RES~abiotic.PC2+(1\|compo)+(1\|location) | abiotic.PC2 = 0 | 0.073 |
| (abiotic.PC2, ABG) \| {fundiv, RES} | AGB~abiotic.PC2+fundiv+RES+(1\|location) | abiotic.PC2 = 0 | 0.917 |
| (abiotic.PC2, litterfall) \| {fundiv, AGB} | litterfall~abiotic.PC2+fundiv+AGB+(1\|location) | abiotic.PC2 = 0 | 0.075 |
| Conditional independance claim | Model formula | $H_0$ | *p*-value |

| | | | |
|---|---|---|---|
| (abiotic.PC2, litter.quality) \| {fundiv, LES} | litter.quality~abiotic.PC2+LES+(1\|location) | abiotic.PC2 = 0 | 0.711 |
| (abiotic.PC2, root) \| {fundiv, AGB} | root~abiotic.PC2+fundiv+AGB+(1\|location) | abiotic.PC2 = 0 | 0.739 |
| (abiotic.PC2, herbaceous) \| {AGB, root} | herbaceous ~abiotic.PC2+AGB+root+(1\|location) | abiotic.PC2 = 0 | 0.767 |
| (abiotic.PC2, multifun) \| {AGB, litterfall, litter.quality, root, herbaceous, microclim} | multifun~ abiotic.PC2+AGB+litterfall+litter.quality+root +herbaceous+microclim+(1\|location) | abiotic.PC2 = 0 | 0.920 |
| (fundiv, litter.quality) \| {SR, LES} | litter.quality~fundiv+SR+LES+(1\|location) | fundiv = 0 | 0.455 |
| (fundiv, herbaceous) \| {SR, AGB, root} | herbaceous~fundiv+SR+AGB+root+(1\|location) | fundiv = 0 | 0.343 |
| (fundiv, multifun) \| {SR, AGB, litterfall, litter.quality, root, herbaceous, microclim} | multifun~fundiv+SR+AGB+litterfall+litter.quality+root +herbaceous+microclim+(1\|location) | fundiv = 0 | 0.638 |
| (AGB, microclim) \| {fundiv, LES, RES, abiotic.PC1, abiotic.PC2 } | microclim~AGB+fundiv+LES +RES+abiotic.PC1 +abiotic.PC2 +(1\|location) | AGB = 0 | 0.132 |
| (litterfall, root) \| {fundiv, AGB} | root~litterfall+fundiv+AGB+(1\|location) | litterfall = 0 | 0.902 |
| (litterfall, microclim) \| {fundiv, AGB, abiotic.PC1, abiotic.PC2, LES, RES} | microclim~litterfall+fundiv+AGB+abiotic.PC1 +abiotic.PC2+LES+RES+(1\|location) | litterfall = 0 | 0.491 |
| (litterfall, herbaceous) \| {fundiv, AGB, root} | herbaceous~litterfall+fundiv+AGB+root+(1\|location) | litterfall = 0 | 0.737 |
| (litter.quality, root) \| {fundiv, LES, AGB} | root~litter.quality+fundiv+LES+AGB+(1\|location) | litter.quality = 0 | 0.607 |
| (litter.quality, microclim) \| {fundiv, LES, RES, abiotic.PC1, abiotic.PC2} | microclim~litter.quality+fundiv+LES+RES +abiotic.PC1 +abiotic.PC2 +(1\|location) | litter.quality = 0 | 0.830 |
| (litter.quality, herbaceous) \| {LES, AGB, root} | herbaceous~litter.quality+LES+AGB+root+(1\|location) | litter.quality = 0 | 0.504 |
| (root, microclim) \| {fundiv, AGB, LES, RES, abiotic.PC1, abiotic.PC2 } | microclim~root+fundiv+AGB+LES+RES+abiotic.PC1 +abiotic.PC2+(1\|location) | root = 0 | 0.254 |
| (microclim, herbaceous) \| {abiotic.PC1, abiotic.PC2, fundiv, LES, RES, AGB, root} | herbaceous~microclim+abiotic.PC1+abiotic.PC2 +fundiv+LES+RES +AGB+root+(1\|location) | microclim = 0 | 0.462 |

**Supplementary Table 8. Nematode families identified, their trophic guild, and their mean individual fresh body mass.** The body mass values were retrieved from the Nemaplex database (http://nemaplex.ucdavis.edu).

| Nematode family | Trophic guild | Individual fresh body mass (in µg) |
|---|---|---|
| Tylenchidae | Herbivore | 0.152 |
| Criconematidae | Herbivore | 0.638 |
| Dolichodoridae | Herbivore | 0.458 |
| Pratylenchidae | Herbivore | 2.732 |
| Hoplolaimidae | Herbivore | 0.738 |
| Paratylenchidae | Herbivore | 0.063 |
| Psilenchidae | Herbivore | 0.489 |
| Cephalobidae | Bacterivore | 0.435 |
| Plectidae | Bacterivore | 0.889 |
| Prismatolaimidae | Bacterivore | 0.487 |
| Metateratocephalidae | Bacterivore | 0.889 |
| Teratocephalidae | Bacterivore | 0.128 |
| Rhabditidae | Bacterivore | 5.393 |
| Panagrolaimidae | Bacterivore | 4.077 |
| Alaimidae | Bacterivore | 0.756 |
| Bunonematidae | Bacterivore | 0.122 |
| Monhysteridae | Bacterivore | 0.423 |
| Bastianiidae | Bacterivore | 0.174 |
| Diplogasteridae | Bacterivore | 1.375 |
| Aphelenchoididae | Fungivore | 0.205 |
| Diphtherophoridae | Fungivore | 0.627 |
| Anguinidae | Fungivore | 11.054 |
| Aphelenchidae | Fungivore | 0.313 |
| Leptonchidae | Fungivore | 0.856 |
| Dorylaimidae | Omnivore | 8.834 |
| Tripylidae | Carnivore | 2.476 |
| Mononchidae | Carnivore | 5.329 |

**Supplementary Table 9. Collembola species identified, their trophic guild and their mean individual fresh and dry body mass.** For each species, the mean dry body mass (*M*, in µg) was calculated from the mean body length (*L*, in mm) retrieved from the BETSI database (https://portail.betsi.cnrs.fr/) using the allometric equation: $M = aL^b$, where $a$ is the normalisation coefficient and $b$ is the exponent. Abdomen length of Symphypleona was used in the original equations and was assumed to be 0.83 of the total body length. Two sets of coefficients coming from two independent studies[22,23] were used for each species ($a_1$, $b_1$ and $a_2$, $b_2$) and the two estimates of dry body mass were averaged[24]. Fresh body mass was calculated from the resulting average by dividing it by the proportion of the dry weight[23,24].

| Family | Species | Trophic guild | Fresh body mass (M, in µg) | Dry body mass (M, in µg) | Fresh body length (in mm) | $a_1$ | $b_1$ | $a_2$ | $b_2$ | Dry weight proportion |
|---|---|---|---|---|---|---|---|---|---|---|
| Entomobryidae | *Entomobrya corticalis* | Epedaphic | 125.64 | 37.69 | 1.50 | 11.75 | 2.52 | 14.26 | 2.71 | 0.30 |
| Entomobryidae | *Entomobrya nicoleti* | Epedaphic | 148.85 | 44.65 | 1.60 | 11.75 | 2.52 | 14.26 | 2.71 | 0.30 |
| Entomobryidae | *Entomobrya sp.* | Epedaphic | 136.94 | 41.08 | 1.55 | 11.75 | 2.52 | 14.26 | 2.71 | 0.30 |
| Entomobryidae | *Heteromurus nitidus* | Epedaphic | 481.19 | 144.36 | 2.50 | 11.75 | 2.52 | 14.26 | 2.71 | 0.30 |
| Entomobryidae | *Heteromurus sp.* | Epedaphic | 481.19 | 144.36 | 2.50 | 11.75 | 2.52 | 14.26 | 2.71 | 0.30 |
| Entomobryidae | *Lepidocyrtus lignorum* | Epedaphic | 202.84 | 60.85 | 1.80 | 11.75 | 2.52 | 14.26 | 2.71 | 0.30 |
| Entomobryidae | *Lepidocyrtus violaceus* | Epedaphic | 136.94 | 41.08 | 1.55 | 11.75 | 2.52 | 14.26 | 2.71 | 0.30 |
| Entomobryidae | *Orchesella bifasciata* | Epedaphic | 267.57 | 80.27 | 2.00 | 11.75 | 2.52 | 14.26 | 2.71 | 0.30 |
| Entomobryidae | *Orchesella cincta* | Epedaphic | 1,166.77 | 350.03 | 3.50 | 11.75 | 2.52 | 14.26 | 2.71 | 0.30 |
| Entomobryidae | *Orchesella flavescens* | Epedaphic | 2,986.80 | 896.04 | 5.00 | 11.75 | 2.52 | 14.26 | 2.71 | 0.30 |
| Entomobryidae | *Orchesella quinquefasciata* | Epedaphic | 2,986.80 | 896.04 | 5.00 | 11.75 | 2.52 | 14.26 | 2.71 | 0.30 |
| Entomobryidae | *Orchesella sp.* | Epedaphic | 1,166.77 | 350.03 | 3.50 | 11.75 | 2.52 | 14.26 | 2.71 | 0.30 |
| Entomobryidae | *Orchesella villosa* | Epedaphic | 3,396.90 | 1,019.07 | 5.25 | 11.75 | 2.52 | 14.26 | 2.71 | 0.30 |
| Entomobryidae | *Pseudosinella sp.* | Hemiedaphic | 43.34 | 13.00 | 1.00 | 11.75 | 2.52 | 14.26 | 2.71 | 0.30 |
| Isotomidae | *Cryptopygus bipunctatus* | Hemiedaphic | 6.13 | 2.21 | 0.70 | 5.62 | 3.28 | 8.43 | 3.22 | 0.36 |
| Isotomidae | *Desoria grijs* | Hemiedaphic | 131.51 | 47.34 | 1.80 | 5.62 | 3.28 | 8.43 | 3.22 | 0.36 |
| Isotomidae | *Desoria neglecta* | Hemiedaphic | 131.51 | 47.34 | 1.80 | 5.62 | 3.28 | 8.43 | 3.22 | 0.36 |
| Isotomidae | *Desoria paars* | Hemiedaphic | 131.51 | 47.34 | 1.80 | 5.62 | 3.28 | 8.43 | 3.22 | 0.36 |
| Isotomidae | *Desoria sp.* | Hemiedaphic | 131.51 | 47.34 | 1.80 | 5.62 | 3.28 | 8.43 | 3.22 | 0.36 |
| Isotomidae | *Folsomia quadrioculata* | Hemiedaphic | 58.17 | 20.94 | 1.40 | 5.62 | 3.28 | 8.43 | 3.22 | 0.36 |
| Isotomidae | *Folsomia sp.* | Hemiedaphic | 58.17 | 20.94 | 1.40 | 5.62 | 3.28 | 8.43 | 3.22 | 0.36 |
| Isotomidae | *Folsomides angularis* | Hemiedaphic | 11.39 | 4.10 | 0.88 | 6.46 | 2.99 | 5.62 | 2.80 | 0.36 |
| Isotomidae | *Folsomides parvulus* | Hemiedaphic | 12.36 | 4.45 | 0.90 | 6.46 | 2.99 | 5.62 | 2.80 | 0.36 |
| Isotomidae | *Isotoma anglicana* | Epedaphic | 1,757.29 | 632.63 | 4.00 | 5.62 | 3.28 | 8.43 | 3.22 | 0.36 |
| Isotomidae | *Isotoma caerulea* | Epedaphic | 1,757.29 | 632.63 | 4.00 | 5.62 | 3.28 | 8.43 | 3.22 | 0.36 |
| Isotomidae | *Isotoma sp.* | Epedaphic | 1,757.29 | 632.63 | 4.00 | 5.62 | 3.28 | 8.43 | 3.22 | 0.36 |
| Isotomidae | *Isotomiella minor* | Euedaphic | 26.59 | 9.57 | 1.10 | 5.62 | 3.28 | 8.43 | 3.22 | 0.36 |
| Isotomidae | *Isotomurus italicus* | Epedaphic | 382.05 | 137.54 | 2.50 | 5.62 | 3.28 | 8.43 | 3.22 | 0.36 |
| Isotomidae | *Isotomurus sp.* | Epedaphic | 382.05 | 137.54 | 2.50 | 5.62 | 3.28 | 8.43 | 3.22 | 0.36 |
| Isotomidae | *Parisotoma notabilis* | Hemiedaphic | 19.51 | 7.03 | 1.00 | 5.62 | 3.28 | 8.43 | 3.22 | 0.36 |
| Isotomidae | *Proisotoma sp.* | Epedaphic | 22.86 | 8.23 | 1.05 | 5.62 | 3.28 | 8.43 | 3.22 | 0.36 |
| Isotomidae | *Tetracanthella sp.* | Hemiedaphic | 35.27 | 12.70 | 1.20 | 5.62 | 3.28 | 8.43 | 3.22 | 0.36 |
| Isotomidae | *Vertagopus sp.* | Epedaphic | 120.02 | 43.21 | 1.75 | 5.62 | 3.28 | 8.43 | 3.22 | 0.36 |
| Onychiuridae | *Onychiuridae sp.* | Euedaphic | 96.71 | 29.01 | 1.90 | 4.27 | 2.75 | 5.60 | 2.77 | 0.30 |

| Family | Species | Trophic guild | Fresh body mass (M, in µg) | Dry body mass (M, in µg) | Fresh body length (in mm) | a1 | b1 | a2 | b2 | Dry weight proportion |
|---|---|---|---|---|---|---|---|---|---|---|
| Onychiuridae | *Onychiurus sp.* | Euedaphic | 96.71 | 29.01 | 1.90 | 4.27 | 2.75 | 5.60 | 2.77 | 0.30 |
| Onychiuridae | *Protaphorura sp.* | Epedaphic | 111.43 | 33.43 | 2.00 | 4.27 | 2.75 | 5.60 | 2.77 | 0.30 |
| Tomoceridae | *Pogonognathellus flavescens* | Epedaphic | 6,163.09 | 1,540.77 | 6.00 | 9.20 | 2.74 | 14.26 | 2.71 | 0.25 |
| Tomoceridae | *Pogonognathellus longicornis* | Epedaphic | 3,751.59 | 937.90 | 5.00 | 9.20 | 2.74 | 14.26 | 2.71 | 0.25 |
| Tomoceridae | *Tomocerina minuta* | Hemiedaphic | 331.15 | 82.79 | 2.05 | 9.20 | 2.74 | 14.26 | 2.71 | 0.25 |
| Tomoceridae | *Tomocerus vulgaris* | Epedaphic | 2,043.47 | 510.87 | 4.00 | 9.20 | 2.74 | 14.26 | 2.71 | 0.25 |
| Neelidae | *Megalothorax minimus* | Euedaphic | 19.23 | 5.77 | 0.40 | 39.99 | 2.11 | 39.99 | 2.11 | 0.30 |
| Hypogastruridae | *Ceratophysella denticulata* | Hemiedaphic | 120.41 | 36.12 | 1.80 | 9.77 | 2.55 | 5.60 | 2.77 | 0.30 |
| Hypogastruridae | *Ceratophysella sp.* | Hemiedaphic | 120.41 | 36.12 | 1.80 | 9.77 | 2.55 | 5.60 | 2.77 | 0.30 |
| Hypogastruridae | *Choreutinula inermis* | Hemiedaphic | 25.62 | 7.69 | 1.00 | 9.77 | 2.55 | 5.60 | 2.77 | 0.30 |
| Hypogastruridae | *Choreutinula sp.* | Hemiedaphic | 25.62 | 7.69 | 1.00 | 9.77 | 2.55 | 5.60 | 2.77 | 0.30 |
| Hypogastruridae | *Schoetella sp.* | Hemiedaphic | 25.62 | 7.69 | 1.00 | 9.77 | 2.55 | 5.60 | 2.77 | 0.30 |
| Hypogastruridae | *Schoetella ununguiculata* | Hemiedaphic | 25.62 | 7.69 | 1.00 | 9.77 | 2.55 | 5.60 | 2.77 | 0.30 |
| Hypogastruridae | *Willemia denisi* | Euedaphic | 11.00 | 3.30 | 0.73 | 9.77 | 2.55 | 5.60 | 2.77 | 0.30 |
| Hypogastruridae | *Willemia sp.* | Euedaphic | 11.00 | 3.30 | 0.73 | 9.77 | 2.55 | 5.60 | 2.77 | 0.30 |
| Isotomidae | *Anurophorus laricis* | Hemiedaphic | 62.10 | 18.63 | 1.40 | 9.77 | 2.55 | 5.60 | 2.77 | 0.30 |
| Isotomidae | *Anurophorus septentrionalis* | Hemiedaphic | 32.92 | 9.87 | 1.10 | 9.77 | 2.55 | 5.60 | 2.77 | 0.30 |
| Neanuridae | *Anurida grey* | Predators | 25.62 | 7.69 | 1.00 | 9.77 | 2.55 | 5.60 | 2.77 | 0.30 |
| Neanuridae | *Anurida sp.* | Predators | 25.62 | 7.69 | 1.00 | 9.77 | 2.55 | 5.60 | 2.77 | 0.30 |
| Neanuridae | *Friesea mirabilis* | Predators | 68.11 | 20.43 | 1.45 | 9.77 | 2.55 | 5.60 | 2.77 | 0.30 |
| Neanuridae | *Friesea subterranea* | Predators | 12.03 | 3.61 | 0.75 | 9.77 | 2.55 | 5.60 | 2.77 | 0.30 |
| Neanuridae | *Frisea sp.* | Predators | 16.71 | 5.01 | 0.85 | 9.77 | 2.55 | 5.60 | 2.77 | 0.30 |
| Neanuridae | *Lathriopyga longiseta* | Predators | 286.46 | 85.94 | 2.50 | 9.77 | 2.55 | 5.60 | 2.77 | 0.30 |
| Neanuridae | *Micranurida forslundi* | Predators | 19.42 | 5.83 | 0.90 | 9.77 | 2.55 | 5.60 | 2.77 | 0.30 |
| Neanuridae | *Micranurida granulata* | Predators | 4.15 | 1.24 | 0.50 | 9.77 | 2.55 | 5.60 | 2.77 | 0.30 |
| Neanuridae | *Micranurida pygmaea* | Predators | 4.15 | 1.24 | 0.50 | 9.77 | 2.55 | 5.60 | 2.77 | 0.30 |
| Neanuridae | *Micranurida sp.* | Predators | 4.15 | 1.24 | 0.50 | 9.77 | 2.55 | 5.60 | 2.77 | 0.30 |
| Neanuridae | *Monobella sp.* | Predators | 74.47 | 22.34 | 1.50 | 9.77 | 2.55 | 5.60 | 2.77 | 0.30 |
| Neanuridae | *Neanura alba* | Predators | 120.41 | 36.12 | 1.80 | 9.77 | 2.55 | 5.60 | 2.77 | 0.30 |
| Neanuridae | *Neanura muscorum* | Predators | 696.89 | 209.07 | 3.50 | 9.77 | 2.55 | 5.60 | 2.77 | 0.30 |
| Neanuridae | *Pseudachorutes dubius* | Predators | 463.67 | 139.10 | 3.00 | 9.77 | 2.55 | 5.60 | 2.77 | 0.30 |
| Odontellidae | *Odontella sp.* | Hemiedaphic | 25.62 | 7.69 | 1.00 | 9.77 | 2.55 | 5.60 | 2.77 | 0.30 |
| Odontellidae | *Superodontella lamellifera* | Hemiedaphic | 68.11 | 20.43 | 1.45 | 9.77 | 2.55 | 5.60 | 2.77 | 0.30 |
| Onychiuridae | *Kalaphorura burmeisteri* | Euedaphic | 14.25 | 4.27 | 0.80 | 9.77 | 2.55 | 5.60 | 2.77 | 0.30 |
| Onychiuridae | *Micraphorura absoloni* | Euedaphic | 14.25 | 4.27 | 0.80 | 9.77 | 2.55 | 5.60 | 2.77 | 0.30 |
| Onychiuridae | *Micraphorura sp.* | Euedaphic | 14.25 | 4.27 | 0.80 | 9.77 | 2.55 | 5.60 | 2.77 | 0.30 |
| Tullbergiidae | *Tullbergiidae* | Euedaphic | 6.69 | 2.01 | 0.60 | 9.77 | 2.55 | 5.60 | 2.77 | 0.30 |
| Arrhopalitidae | *Arrhopalites principalis* | Hemiedaphic | 277.32 | 58.24 | 1.00 | 190.55 | 3.63 | 39.63 | 3.80 | 0.21 |
| Arrhopalitidae | *Arrhopalites sp.* | Hemiedaphic | 548.03 | 115.09 | 1.00 | 190.55 | 3.63 | 39.63 | 3.80 | 0.21 |

| Family | Species | Trophic guild | Fresh body mass (M, in µg) | Dry body mass (M, in µg) | Fresh body length (in mm) | a1 | b1 | a2 | b2 | Dry weight proportion |
|---|---|---|---|---|---|---|---|---|---|---|
| Bourletiellidae | *Deuterosminthurus bicinctus* | Epedaphic | 242.40 | 50.90 | 0.80 | 190.55 | 3.63 | 39.63 | 3.80 | 0.21 |
| Dicyrtomidae | *Dicyrtoma fusca* | Epedaphic | 6,915.72 | 1,452.30 | 2.00 | 190.55 | 3.63 | 39.63 | 3.80 | 0.21 |
| Dicyrtomidae | *Dicyrtomidae sp.* | Epedaphic | 6,915.72 | 1,452.30 | 2.00 | 190.55 | 3.63 | 39.63 | 3.80 | 0.21 |
| Dicyrtomidae | *Dicyrtomina ornata* | Epedaphic | 15,648.81 | 3,286.25 | 2.50 | 190.55 | 3.63 | 39.63 | 3.80 | 0.21 |
| Dicyrtomidae | *Dicyrtomina sp.* | Epedaphic | 15,648.81 | 3,286.25 | 2.50 | 190.55 | 3.63 | 39.63 | 3.80 | 0.21 |
| Katiannidae | *Sminthurinus aureus* | Epedaphic | 372.84 | 78.30 | 0.90 | 190.55 | 3.63 | 39.63 | 3.80 | 0.21 |
| Katiannidae | *Sminthurinus sp.* | Epedaphic | 271.26 | 56.96 | 0.83 | 190.55 | 3.63 | 39.63 | 3.80 | 0.21 |
| Katiannidae | *Sminturinus bimaculatus* | Epedaphic | 148.79 | 31.25 | 0.70 | 190.55 | 3.63 | 39.63 | 3.80 | 0.21 |
| Katiannidae | *Sminturinus elegans* | Epedaphic | 148.79 | 31.25 | 0.70 | 190.55 | 3.63 | 39.63 | 3.80 | 0.21 |
| Katiannidae | *Sminturinus signatus* | Epedaphic | 271.26 | 56.96 | 0.83 | 190.55 | 3.63 | 39.63 | 3.80 | 0.21 |
| Sminthuridae | *Allacma fusca* | Epedaphic | 56,491.17 | 11,863.14 | 3.55 | 190.55 | 3.63 | 39.63 | 3.80 | 0.21 |
| Sminthuridae | *Caprainea marginata* | Epedaphic | 1875.71 | 393.90 | 1.40 | 190.55 | 3.63 | 39.63 | 3.80 | 0.21 |
| Sminthuridae | *Caprainea sp.* | Epedaphic | 1875.71 | 393.90 | 1.40 | 190.55 | 3.63 | 39.63 | 3.80 | 0.21 |
| Sminthuridae | *Lipothrix lubbocki* | Epedaphic | 6915.72 | 1452.30 | 2.00 | 190.55 | 3.63 | 39.63 | 3.80 | 0.21 |
| Sminthuridae | *Lipothrix sp.* | Epedaphic | 6915.72 | 1452.30 | 2.00 | 190.55 | 3.63 | 39.63 | 3.80 | 0.21 |
| Sminthuridae | *Spatulosminthurus flaviceps* | Epedaphic | 3,056.95 | 641.96 | 1.60 | 190.55 | 3.63 | 39.63 | 3.80 | 0.21 |
| Sminthurididae | *Sminthuridae sp.* | Epedaphic | 548.03 | 115.09 | 1.00 | 190.55 | 3.63 | 39.63 | 3.80 | 0.21 |
| Sminthurididae | *Sminthurides sp.* | Epedaphic | 43.51 | 9.14 | 0.50 | 190.55 | 3.63 | 39.63 | 3.80 | 0.21 |
| Sminthurididae | *Sphaeridia pumilis* | Epedaphic | 19.26 | 4.04 | 0.40 | 190.55 | 3.63 | 39.63 | 3.80 | 0.21 |
| Sminthurididae | *Sphaeridia sp.* | Epedaphic | 19.26 | 4.04 | 0.40 | 190.55 | 3.63 | 39.63 | 3.80 | 0.21 |
| Symphypleona | Symphypleona | Epedaphic | 1,703.59 | 357.75 | 1.36 | 190.55 | 3.63 | 39.63 | 3.80 | 0.21 |

**Supplementary Table 10. Earthworm species identified, and their ecological groups.** Earthworm species were classified based on the DriloBASE database (http://taxo.drilobase.org).

| Species | Ecological group |
|---|---|
| *Allolobophora carpathica* | Endogeic |
| *Allolobophora sturanyi dacidoides* | Endogeic |
| *Aporrectodea caliginosa* | Endogeic |
| *Aporrectodea rosea* | Endogeic |
| *Dendrobaena attemsi* | Epigeic |
| *Dendrobaena octaedra* | Epigeic |
| *Dendrobaena platyura* | Anecic |
| *Dendrobaena sp.* | Epigeic |
| *Lumbricus rubellus* | Epigeic |
| *Lumbricus sp.* | Epigeic |
| *Lumbricus terrestris* | Anecic |
| *Octodrillus complatus* | Endogeic |
| *Octolasium lacteum* | Endogeic |

**Supplementary Table 11. Regression parameters used to calculate individual metabolic rates for faunal consumers.** This was based on this equation: $\ln I = \ln i_0 + \mathrm{a} \ln M - E(1/kT)$, where $I$ is the metabolic rate, $i_0$ is a normalisation factor, $a$ is the allometric exponent, $E$ is the activation energy, $k$ is the Boltzmann constant, and $T$ is the temperature in Kelvin. The taxa-specific parameters were used if available, and general parameters were used otherwise. These parameter values are from Ehnes et al. (2011)[100] and unpublished data (Roswitha Ehnes).

| Regression model | Phylogenetic contrast | Taxa | ln $i_0$ | $a$ | $E$ |
|---|---|---|---|---|---|
| Linear | General | Nematoda, Gasteropoda | 23.055 | 0.695 | 0.686 |
| Phylogenetic | Arachnida | Araneae, Opiliones | 24.581 | 0.565 | 0.709 |
| Phylogenetic | Chilopoda | Chilopoda | 28.253 | 0.558 | 0.803 |
| Phylogenetic | Clitellata | Clitellata | 12.442 | 0.801 | 0.443 |
| Phylogenetic | Coleoptera | Coleoptera | 21.418 | 0.738 | 0.639 |
| Phylogenetic | Insecta | Blattodea, Collembola, Dermaptera, Diptera, Formicidae, Lepidoptera | 21.972 | 0.759 | 0.657 |
| Phylogenetic | Isopoda | Isopoda | 23.169 | 0.554 | 0.687 |
| Phylogenetic | Mesostigmata | Mesostigmata | 9.674 | 0.690 | 0.379 |
| Phylogenetic | Oribatida | Oribatida | 22.023 | 0.679 | 0.706 |
| Phylogenetic | Progoneata | Diplopoda | 22.347 | 0.571 | 0.670 |
| Phylogenetic | Prostigmata | Astigmata, Prostigmata | 10.281 | 0.660 | 0.413 |

**Supplementary Table 12. Diet-specific assimilation efficiency values averaged across all plots for faunal consumers.** These values were calculated from Jochum et al. (2017)[36] based on food N content, and corrected for temperature based on Lang et al. (2017)[37]. The growing season soil temperature averaged across all plots was used for temperature correction. Note that the assimilation efficiency of living plant fine roots by herbivores is much lower than the assimilation efficiency of plant leaves (0.545) based on Lang et al. (2017)[37] but this is consistent with the assimilation efficiency of plant roots (~0.180) based on Gan & Wickings (2020)[101].

| Food | Food N content | Assimilation efficiency uncorrected for temperature | Assimilation efficiency corrected for temperature |
|---|---|---|---|
| Living plant fine roots | 0.014 | 0.191 | 0.168 |
| Dead leaves | 0.011 | 0.170 | 0.148 |
| Dead roots | 0.014 | 0.191 | 0.168 |
| Dead wood | 0.003 | 0.124 | 0.102 |
| Soil organic matter | 0.002 | 0.121 | 0.099 |
| Bacteria: Gram- | 0.077 | 0.820 | 0.803 |
| Bacteria: Gram+ | 0.077 | 0.820 | 0.803 |
| Mycorrhizal fungi | 0.034 | 0.379 | 0.363 |
| General saprotroph fungi | 0.040 | 0.447 | 0.430 |
| Wood saprotroph fungi | 0.034 | 0.379 | 0.363 |
| Plant pathogenic fungi | 0.040 | 0.447 | 0.430 |
| Nematoda: Plant feeders | 0.053 | 0.599 | 0.582 |
| Nematoda: Bacterivores | 0.053 | 0.599 | 0.582 |
| Nematoda: Fungivores | 0.053 | 0.599 | 0.582 |
| Nematoda: Omnivores | 0.053 | 0.599 | 0.582 |
| Nematoda: Predators | 0.053 | 0.599 | 0.582 |
| Oribatida | 0.089 | 0.890 | 0.874 |
| Mesostigmata | 0.105 | 0.945 | 0.929 |
| Prostigmata & Astigmata | 0.101 | 0.935 | 0.918 |
| Collembola: Epedaphic | 0.109 | 0.954 | 0.938 |
| Collembola: Hemiedaphic | 0.109 | 0.954 | 0.938 |
| Collembola: Euedaphic | 0.109 | 0.954 | 0.938 |
| Collembola: Predators | 0.109 | 0.954 | 0.938 |
| Araneae: Small | 0.111 | 0.958 | 0.941 |
| Araneae: Large | 0.111 | 0.958 | 0.941 |
| Opiliones | 0.120 | 0.972 | 0.956 |
| Diplopoda | 0.054 | 0.610 | 0.593 |
| Chilopoda: Geophilomorpha | 0.117 | 0.968 | 0.951 |
| Chilopoda: Scolopendromorpha | 0.117 | 0.968 | 0.951 |
| Chilopoda: Lithobiomorpha | 0.117 | 0.968 | 0.951 |
| Isopoda | 0.066 | 0.733 | 0.717 |
| Dermaptera | 0.092 | 0.903 | 0.887 |
| Blattodea | 0.111 | 0.958 | 0.941 |
| Formicidae | 0.113 | 0.962 | 0.945 |
| Coleoptera: Carabidae | 0.100 | 0.932 | 0.915 |
| Coleoptera: Staphylinidae | 0.100 | 0.932 | 0.915 |
| Coleoptera: Elateridae | 0.090 | 0.895 | 0.878 |
| Coleoptera: Scarabaeidae/Geotrupidae | 0.090 | 0.895 | 0.878 |
| Coleoptera: Curculionidae | 0.090 | 0.895 | 0.878 |
| Lepidoptera | 0.082 | 0.854 | 0.837 |
| Diptera | 0.096 | 0.919 | 0.902 |
| Lumbricina: Epigeic | 0.122 | 0.975 | 0.958 |
| Lumbricina: Anecic | 0.122 | 0.975 | 0.958 |
| Lumbricina: Endogeic | 0.122 | 0.975 | 0.958 |
| Gastropoda: Snails | 0.042 | 0.470 | 0.454 |
| Gastropoda: Slugs | 0.094 | 0.911 | 0.894 |

**Supplementary Table 13. Regression parameters used to calculate the temperature correction for assimilation efficiency.** $\acute{\varepsilon}_0$ is the normalization constant of the assimilation efficiency, $E_\varepsilon$ is the activation energy for assimilation efficiency (eV), and $\varepsilon_0$ is the assimilation efficiency at 20°C. The parameter values are from Lang et al. (2017)[37].

| Consumer type | Consumed food type | $\acute{\varepsilon}_0$ | $E_\varepsilon$ | $\varepsilon_0$ |
|---|---|---|---|---|
| Herbivores | Living plant fine roots | -1.670 | | 0.545 |
| Detritivores | Plant litter and soil organic matter | 0.179 | 0.164 | 0.158 |
| Carnivores | Fauna and microbes | 2.266 | | 0.906 |

**Supplementary Table 14. Results of Bayesian multi-level random slope models**. Both the leaf and fine-root economics spectra (LES & RES) variables representing tree functional composition ranged from slow/conservative to fast/acquisitive attributes (Extended Data Fig. 3a). % var, proportion of total variance. $\beta_{st}$, slope regression coefficients standardised by standard deviation. CI 95%, 95% credible intervals. BF, Bayes factor (Methods). Significant effects ($p < 0.05$ or BF > 1) are reported in bold. †, $p < 0.10$; *, $p < 0.050$; **, $p < 0.010$; ***, $p < 0.001$. SOM, soil organic matter. $\Delta_i$, difference in LOO information criterion values between the random slope model and the random intercept model. Negative $\Delta_i$ indicate better goodness-of-fit for the random slope model than the random intercept model. When $|\Delta_i| < 6$, difference in goodness-of-fit is considered small, *i.e.* both models have comparable support in the data[102].

| | Taxonomic approach for tree community effects | | | | | | Functional approach for tree community effects | | | | | | | | |
|---|---|---|---|---|---|---|---|---|---|---|---|---|---|---|---|
| | Species richness | | | Species composition | | $\Delta_i$ | Functional diversity | | | Functional composition | | | | | $\Delta_i$ |
| | | | | | | | | | | | Leaf economics spectrum | | Fine-root economics spectrum | | |
| | % var | $\beta_{st}$ [CI 95%] | *p*-value | % var | BF | | % var | $\beta_{st}$ [CI 95%] | *p*-value | % var | $\beta_{st}$ [CI 95%] | *p*-value | $\beta_{st}$ [CI 95%] | *p*-value | |
| **Soil food web functioning** | | | | | | | | | | | | | | | |
| Carnivory | 3.75 | 0.00 [-0.38;0.40] | 0.997 | 7.19 | 0.601 | -1.70 | 1.20 | 0.03 [-0.21;0.28] | 0.837 | 32.15 | 0.35 [0.00;0.71] | **0.049*** | 0.48 [-0.01;0.96] | 0.054† | +1.09 |
| Microbivory | 3.94 | -0.04 [-0.43;0.34] | 0.781 | 13.62 | 0.87 | +3.19 | 0.68 | -0.03 [-0.25;0.19] | 0.775 | 37.54 | 0.32 [-0.02;0.61] | 0.061† | 0.78 [0.26;1.20] | **0.014*** | +1.11 |
| *Fungivory* | 3.34 | -0.01 [-0.38;0.35] | 0.942 | 8.46 | 0.52 | +2.81 | 0.63 | 0.01 [-0.23;0.25] | 0.916 | 26.52 | 0.27 [-0.12;0.60] | 0.133 | 0.73 [0.19;1.19] | **0.013*** | +1.82 |
| *Bacterivory* | 3.38 | -0.10 [-0.41;0.22] | 0.478 | 4.50 | 0.62 | -1.28 | 1.92 | -0.11 [-0.26;0.06] | 0.194 | 24.01 | 0.20 [-0.12;0.51] | 0.164 | 0.40 [-0.12;0.78] | 0.066† | +0.67 |
| Detritivory | 5.25 | -0.14 [-0.50;0.24] | 0.422 | 8.28 | 0.46 | -1.35 | 2.76 | -0.15 [-0.42;0.14] | 0.277 | 16.24 | 0.20 [-0.21;0.59] | 0.263 | 0.33 [-0.27;0.91] | 0.214 | +0.59 |
| *Soil engineering* | 6.06 | -0.18 [-0.51;0.13] | 0.223 | 5.95 | 0.43 | 0.07 | 2.61 | -0.16 [-0.44;0.11] | 0.220 | 14.62 | 0.34 [-0.19;0.86] | 0.161 | 0.07 [-0.64;0.74] | 0.796 | -3.74 |
| *SOM decomposition* | 9.50 | -0.25 [-0.58;0.09] | 0.130 | 24.24 | **3.52** | -0.89 | 4.75 | -0.22 [-0.50;-0.08] | 0.121 | 9.94 | 0.19 [-0.28;0.66] | 0.343 | 0.02 [-0.61;0.69] | 0.947 | +0.33 |
| *Litter engineering* | 4.94 | 0.11 [-0.28;0.49] | 0.543 | 13.70 | **1.06** | +1.95 | 2.57 | 0.15 [-0.12;0.41] | 0.263 | 21.15 | 0.29 [-0.22;0.70] | 0.176 | 0.36 [-0.32;0.89] | 0.238 | +2.27 |
| *Litter decomposition* | 5.72 | -0.05 [-0.52;0.41] | 0.819 | 3.54 | 0.39 | -4.85 | 1.53 | -0.06 [-0.37;0.23] | 0.667 | 21.28 | 0.14 [-0.33;0.57] | 0.452 | 0.48 [-0.12;1.08] | 0.102 | +0.49 |
| Herbivory | 6.77 | -0.15 [-0.56;0.28] | 0.427 | 11.31 | 0.94 | +2.69 | 1.54 | -0.12 [-0.41;0.21] | 0.389 | 18.07 | 0.43 [-0.12;0.95] | 0.105 | 0.38 [-0.25;0.98] | 0.203 | -4.02 |
| *Rhizophagy* | 5.27 | -0.13 [-0.51;0.24] | 0.422 | 8.28 | 0.46 | +5.02 | 1.58 | -0.15 [-0.43;0.15] | 0.266 | 21.49 | 0.25 [-0.37;0.86] | 0.354 | 0.66 [-0.03;1.47] | 0.066† | -3.27 |
| *Root pathogenicity* | 3.32 | -0.02 [-0.39;0.34] | 0.880 | 26.71 | **4.98** | +0.61 | 1.04 | 0.03 [-0.24;0.33] | 0.827 | 19.31 | 0.45 [0.05;0.93] | 0.067† | 0.20 [-0.55;0.87] | 0.457 | -0.97 |
| Plant C allocation to soil by roots | 4.32 | -0.08 [-0.43;0.31] | 0.601 | 8.95 | 0.75 | +0.27 | 1.65 | -0.08 [-0.37;0.23] | 0.549 | 10.83 | 0.07 [-0.42;0.61] | 0.799 | 0.16 [-0.59;0.98] | 0.707 | +3.89 |
| Soil food web multifunctionality | 8.27 | -0.16 [-0.62;0.32] | 0.435 | 20.51 | **2.07** | -0.21 | 1.67 | -0.12 [-0.39;0.17] | 0.368 | 31.97 | 0.47 [0.03;0.88] | **0.037*** | 0.60 [0.01;1.13] | **0.046*** | -0.80 |

**Supplementary Table 15. Adjacency matrix of feeding preferences averaged across all plots.** The values indicate the trophic interaction strengths [0-1] between a consumer and its resources. The sum of all resource values of a given consumer is always equal to one. Consumers and food resources are respectively in columns and rows. See Supplementary Table 2 for abbreviation meanings.

| | BaGn | BaGp | FuM | FuS | FuW | FuP | NePl | NeBa | NeFu | NeOmn | NePr | AcOri | AcMeso | AcOther | ColEp | ColHe | ColEu | ColPr | ArSm | ArLa | Opi | Dpod | ChiGe | ChiSco | ChiLi | Iso | Der | Bla | Ant | CptCar | CptSta | CptOmn | CptHD | CptH | Lepi | Dipt | EwEp | EwAn | EwEn | GaSn | GaSl |
|---|---|---|---|---|---|---|---|---|---|---|---|---|---|---|---|---|---|---|---|---|---|---|---|---|---|---|---|---|---|---|---|---|---|---|---|---|---|---|---|---|---|
| LivRoot | | | | | | 1.00 | 0.71 | | 0.14 | | | | | 0.02 | | | 0.05 | | | | | 0.02 | | | | 0.01 | 0.01 | 0.01 | 0.36 | | | 0.07 | 0.08 | 0.65 | 0.71 | | 0.01 | | | | |
| PR | 1.00 | | 1.00 | 0.04 | | | | | | | | | | | | | | | | | | | | | | | | | | | | | | | | | | | | | |
| LeafLit | | 0.30 | | 0.63 | | | | | | | | 0.48 | | 0.10 | 0.24 | 0.48 | 0.18 | | | | | 0.86 | | | | 0.58 | 0.38 | 0.83 | | | | 0.23 | 0.55 | | | 0.32 | 0.70 | 0.13 | | 0.39 | 0.38 |
| RootLit | | 0.02 | | 0.04 | | | | | | | | 0.05 | | 0.01 | 0.01 | 0.05 | 0.02 | | | | | 0.04 | | | | 0.01 | 0.01 | 0.02 | | | | 0.03 | 0.03 | | | 0.01 | 0.03 | 0.02 | 0.01 | 0.01 | 0.02 |
| WoodLit | | | | | 1.00 | | | | | | | | | | | | | | | | | 0.01 | | | | | | 0.01 | | | | 0.01 | 0.02 | 0.02 | | | 0.01 | | | | |
| SOM | | 0.69 | | 0.29 | | | | | | | | | | | | | | | | | | | | | | | | | | | | | | | | | | 0.65 | 0.72 | | |
| BaGn | | | | | | | | 0.25 | | 0.13 | 0.07 | | | | | | 0.01 | | | | | | | | | 0.01 | 0.01 | | | | | | | | | | 0.01 | 0.01 | 0.02 | 0.03 | 0.03 |
| BaGp | | | | | | | | 0.75 | | 0.37 | 0.21 | 0.01 | | 0.01 | | 0.01 | 0.02 | | | | | | | | | 0.02 | 0.02 | | | | | | 0.01 | | | | 0.04 | 0.03 | 0.05 | 0.09 | 0.09 |
| FuM | | | | | | | | | 0.46 | | | 0.22 | | 0.18 | 0.22 | 0.22 | 0.35 | | | | | | | | | 0.11 | | | | | | 0.09 | 0.14 | | | 0.06 | | | | 0.16 | 0.18 |
| FuS | | | | | | | 0.27 | | 0.39 | | | 0.18 | 0.17 | 0.15 | 0.53 | 0.18 | 0.29 | 0.17 | | | | 0.08 | | | | 0.27 | 0.07 | 0.07 | 0.14 | | 0.17 | 0.07 | 0.18 | 0.31 | 0.27 | 0.11 | 0.13 | 0.10 | 0.14 | 0.27 | 0.26 |
| FuW | | | | | | | | | | | | | | | | | | | | | | | | | | | | | | | | | | 0.01 | | | | | | | |
| FuP | | | | | | | 0.01 | | | | | | | | | | | | | | | | | | | | | | 0.01 | | | | | 0.01 | 0.01 | | | | | | |
| NePl | | | | | | | | | | 0.07 | 0.14 | 0.01 | 0.08 | 0.15 | | | 0.01 | 0.01 | | | | | | | | | | | | | | | | | | | 0.03 | 0.02 | 0.03 | | |
| NeBa | | | | | | | | | | 0.13 | 0.20 | 0.02 | 0.16 | 0.16 | | 0.01 | 0.02 | 0.03 | 0.01 | | | | | | | | | | 0.01 | | 0.01 | | | | | | | 0.02 | 0.02 | | |
| NeFu | | | | | | | | | | 0.01 | 0.02 | | 0.01 | 0.02 | | | | | | | | | | | | | | | | | | | | | | | | | | | |
| NeOmn | | | | | | | | | | 0.14 | 0.18 | 0.01 | 0.22 | 0.10 | | 0.02 | 0.03 | 0.15 | 0.08 | 0.01 | | | 0.03 | 0.03 | 0.02 | | | | 0.03 | | 0.05 | 0.02 | | | | | 0.02 | 0.01 | 0.02 | | |
| NePr | | | | | | | | | | 0.01 | 0.01 | | 0.02 | 0.01 | | | | 0.01 | | | | | | | | | | | | | | | | | | | | | | | |
| AcOri | | | | | | | | | | 0.04 | 0.04 | | 0.09 | 0.02 | | 0.01 | 0.01 | 0.09 | 0.04 | | | | 0.01 | 0.01 | 0.01 | | | | 0.02 | | 0.01 | | | | | | | | | | |
| AcMeso | | | | | | | | | | 0.03 | 0.04 | | 0.06 | 0.02 | | | 0.01 | 0.05 | 0.04 | 0.01 | | | 0.01 | 0.01 | 0.01 | | | | 0.01 | | 0.01 | 0.01 | | | | | | | | | |
| AcOther | | | | | | | | | | | 0.01 | | 0.01 | 0.01 | | | | | | | | | | | | | | | | | | | | | | | | | | | |
| ColEp | | | | | | | | | | 0.01 | 0.01 | | 0.02 | | | | | 0.16 | 0.31 | 0.19 | 0.58 | | 0.08 | 0.21 | 0.23 | | 0.05 | 0.02 | 0.16 | 0.20 | 0.18 | 0.04 | | | | 0.05 | | | | | |
| ColHe | | | | | | | | | | 0.04 | 0.05 | | 0.11 | 0.01 | | 0.01 | 0.01 | 0.18 | 0.14 | 0.02 | | | 0.03 | 0.04 | 0.03 | | | | 0.05 | | 0.05 | 0.02 | | | | | | | | | |
| ColEu | | | | | | | | | | 0.01 | 0.01 | | 0.01 | | | | | 0.02 | 0.01 | | | | | | | | | | | | | | | | | | | | | | |
| ColPr | | | | | | | | | | | | | | | | | | 0.01 | 0.02 | | 0.01 | | | 0.01 | 0.01 | | | | 0.01 | | 0.01 | | | | | | | | | | |
| ArSm | | | | | | | | | | | | | | | | | | | 0.01 | | | | | | | | | | | | | | | | | | | | | | |
| ArLa | | | | | | | | | | | | | | | | | | 0.01 | 0.02 | 0.03 | 0.03 | | 0.01 | 0.02 | 0.03 | | 0.01 | | 0.01 | 0.04 | 0.02 | 0.01 | | | | 0.01 | | | | | |
| Opi | | | | | | | | | | | | | | | | | | | | | 0.01 | | | | | | | | | 0.01 | | | | | | | | | | | |
| Dpod | | | | | | | | | | | | | | | | | | | 0.01 | 0.04 | 0.02 | | 0.04 | 0.04 | 0.03 | | 0.03 | | 0.01 | 0.05 | 0.02 | 0.02 | | | | 0.03 | | | | | |
| ChiGe | | | | | | | | | | | | | | | | | | | 0.01 | 0.01 | | | 0.06 | 0.03 | 0.03 | | 0.01 | | 0.01 | | 0.02 | 0.02 | | | | 0.01 | | | | | |
| ChiSco | | | | | | | | | | | | | | | | | | | | | | | | | | | | | | | | | | | | | | | | | |
| ChiLi | | | | | | | | | | | | | | | | | | | 0.02 | 0.02 | 0.02 | | 0.03 | 0.04 | 0.04 | | 0.01 | | 0.01 | 0.02 | 0.02 | 0.01 | | | | 0.01 | | | | | |
| Iso | | | | | | | | | | | | | | | | | | 0.01 | 0.01 | 0.03 | 0.02 | | 0.01 | 0.02 | 0.02 | | 0.01 | | 0.01 | 0.03 | 0.01 | | | | | 0.01 | | | | | |
| Der | | | | | | | | | | | | | | | | | | | 0.01 | 0.01 | 0.01 | | 0.01 | 0.01 | 0.01 | | 0.01 | | | 0.01 | 0.01 | | | | | 0.01 | | | | | |
| Bla | | | | | | | | | | | | | | | | | | | | | | | | | | | | | | | | | | | | | | | | | |
| Ant | | | | | | | | | | | | | | | | | | | 0.01 | | 0.01 | | | | | | | | | | | | | | | | | | | | |
| CptCar | | | | | | | | | | | | | | | | | | | | 0.01 | 0.01 | | | | 0.01 | | | | | 0.02 | | | | | | | | | | | |
| CptSta | | | | | | | | | | | | | | | | | | | | | | | | | | | | | | | | | | | | | | | | | |
| CptOmn | | | | | | | | | | | | | | | | | | 0.01 | 0.01 | 0.01 | | | 0.04 | 0.03 | 0.02 | | 0.01 | | 0.01 | | 0.02 | 0.02 | | | | 0.01 | | | | | |
| CptHD | | | | | | | | | | | | | 0.01 | | | | | 0.01 | 0.05 | 0.17 | 0.06 | | 0.17 | 0.12 | 0.11 | | 0.09 | 0.01 | 0.04 | 0.17 | 0.11 | 0.09 | | | | 0.09 | | | | 0.01 | 0.01 |
| CptH | | | | | | | | | | | | | | | | | | | 0.01 | 0.01 | | | 0.03 | 0.02 | 0.01 | | 0.01 | | | | 0.01 | 0.01 | | | | 0.01 | | | | | |
| Lepi | | | | | | | | | | | | | | | | | | 0.01 | 0.01 | 0.02 | 0.02 | | 0.02 | 0.02 | 0.02 | | 0.01 | | 0.01 | 0.03 | 0.01 | 0.01 | | | | 0.01 | | | | | |
| Dipt | | | | | | | | | | | | | 0.01 | | | | | 0.03 | 0.08 | 0.15 | 0.11 | | 0.10 | 0.12 | 0.14 | | 0.09 | 0.01 | 0.04 | 0.18 | 0.08 | 0.05 | | | | 0.09 | | | | 0.01 | 0.01 |
| EwEp | | | | | | | | | | | | | | | | | | | 0.02 | 0.12 | 0.02 | | 0.08 | 0.08 | 0.10 | | 0.08 | 0.01 | 0.01 | 0.10 | 0.04 | 0.04 | | | | 0.08 | | | | 0.01 | 0.01 |
| EwAn | | | | | | | | | | | | | | | | | | | | 0.01 | | | 0.03 | 0.01 | 0.01 | | 0.01 | | | | 0.01 | 0.02 | | | | 0.01 | | | | | |
| EwEn | | | | | | | | | | | | | | | | | | | | | | | 0.15 | 0.05 | | | | | 0.01 | | 0.05 | 0.08 | | | | | | | | | |
| GaSn | | | | | | | | | | | | | 0.01 | | | | | 0.02 | 0.05 | 0.08 | 0.06 | | 0.06 | 0.07 | 0.08 | | 0.05 | 0.01 | 0.02 | 0.10 | 0.05 | 0.02 | | | | 0.05 | | | | | |
| GaSl | | | | | | | | | | | | | | | | | | | | 0.01 | | | 0.01 | 0.01 | 0.01 | | 0.01 | | | 0.01 | 0.01 | | | | | 0.01 | | | | | |

**Supplementary references**

1. Frostegard, A. & Bååth, E. The use of phospholipid fatty acid analysis to estimate bacterial and fungal biomass in soil. *Biol. Fertil. Soils* **22**, 59–65 (1996).
2. Prada-Salcedo, L. D., Wambsganss, J., Bauhus, J., Buscot, F. & Goldmann, K. Low root functional dispersion enhances functionality of plant growth by influencing bacterial activities in European forest soils. *Environ. Microbiol.* **23**, 1889–1906 (2021).
3. Bligh, E. G. & Dyer, W. J. A rapid method of total lipid extraction and purification. *Can. J. Biochem. Physiol.* **37**, 911–917 (1959).
4. Joergensen, R. G. Phospholipid fatty acids in soil—drawbacks and future prospects. *Biol. Fertil. Soils* **58**, 1–6 (2022).
5. Olsson, P. A., Bååth, E., Jakobsen, I. & Söderström, B. The use of phospholipid and neutral lipid fatty acids to estimate biomass of arbuscular mycorrhizal fungi in soil. *Mycol. Res.* **99**, 623–629 (1995).
6. Klamer, M. & Bååth, E. Estimation of conversion factors for fungal biomass determination in compost using ergosterol and PLFA 18:2ω6,9. *Soil Biol. Biochem.* **36**, 57–65 (2004).
7. Potapov, A. M. *et al.* Feeding habits and multifunctional classification of soil-associated consumers from protists to vertebrates. *Biol. Rev.* **97**, 1057–1117 (2022).
8. Waring, B. G., Averill, C. & Hawkes, C. V. Differences in fungal and bacterial physiology alter soil carbon and nitrogen cycling: insights from meta-analysis and theoretical models. *Ecol. Lett.* **16**, 887–894 (2013).
9. Fierer, N., Strickland, M. S., Liptzin, D., Bradford, M. A. & Cleveland, C. C. Global patterns in belowground communities. *Ecol. Lett.* **12**, 1238–1249 (2009).
10. Tedersoo, L. *et al.* Best practices in metabarcoding of fungi: From experimental design to results. *Mol. Ecol.* **31**, 2769–2795 (2022).
11. Prada-Salcedo, L. D. *et al.* Fungal guilds and soil functionality respond to tree community traits rather than to tree diversity in European forests. *Mol. Ecol.* **30**, 572–591 (2021).
12. Nilsson, R. H. *et al.* The UNITE database for molecular identification of fungi: handling dark taxa and parallel taxonomic classifications. *Nucleic Acids Res.* **47**, D259–D264 (2019).
13. Nguyen, N. H. *et al.* FUNGuild: An open annotation tool for parsing fungal community datasets by ecological guild. *Fungal Ecol.* **20**, 241–248 (2016).
14. Collado, E. *et al.* Divergent above- and below-ground responses of fungal functional groups to forest thinning. *Soil Biol. Biochem.* **150**, 108010 (2020).
15. Tedersoo, L. *et al.* Tree diversity and species identity effects on soil fungi, protists and animals are context dependent. *ISME J.* **10**, 346–362 (2016).
16. Hagenbo, A. *et al.* Variations in biomass of fungal guilds are primarily driven by factors related to soil conditions in Mediterranean Pinus pinaster forests. *Biol. Fertil. Soils* **58**, 487–501 (2022).
17. Cheeke, T. E., Phillips, R. P., Kuhn, A., Rosling, A. & Fransson, P. Variation in hyphal production rather than turnover regulates standing fungal biomass in temperate hardwood forests. *Ecology* **102**, e03260 (2021).
18. Ehnes, R. B., Rall, B. C. & Brose, U. Phylogenetic grouping, curvature and metabolic scaling in terrestrial invertebrates. *Ecol. Lett.* **14**, 993–1000 (2011).
19. Andrassy, I. Die rauminhalts-und gewichtsbestimmung der fadenwürmer (Nematoden). *Acta Zool. Hung.* **2**, 1–5 (1956).
20. Yeates, G. W. Soil nematodes in terrestrial ecosystems. *J. Nematol.* **11**, 213–229 (1979).
21. Yeates, G. W., Bongers, T., Degoede, R. G. M., Freckman, D. W. & Georgieva, S. S. Feeding Habits in Soil Nematode Families and Genera—An Outline for Soil Ecologists. *J. Nematol.* **25**, 315–331 (1993).
22. Tanaka, M. Ecological studies on communities of soil Collembola in Mt. Sobo, southwest Japan. *Jpn. J. Ecol.* **20**, 102–110 (1970).

23. Petersen, H. Estimation of dry weight, fresh weight, and calorific content of various Collembolan species. *Pedobiologia* **15**, 222–243 (1975).
24. Potapov, A. M. *et al.* Globally invariant metabolism but density-diversity mismatch in springtails. *Nat. Commun.* **14**, 674 (2023).
25. Petersen, H. & Luxton, M. A comparative analysis of soil fauna populations and their role in decomposition processes. *Oikos* **39**, 287–388 (1982).
26. Edwards, C. A. Relationship between weights, volumes and numbers of soil animals. in *Progress in Soil Biology* 585–591 (North-Holland Publishing Company, New York, 1967).
27. Ganault, P. *et al.* Relative importance of tree species richness, tree functional type, and microenvironment for soil macrofauna communities in European forests. *Oecologia* **196**, 455–468 (2021).
28. Mercer, R. D., Gabriel, A. G. A., Barendse, J., Marshall, D. J. & Chown, S. L. Invertebrate body sizes from Marion Island. *Antarct. Sci.* **13**, 135–143 (2001).
29. Gillespie, L. M. *et al.* Tree species mixing affects soil microbial functioning indirectly via root and litter traits and soil parameters in European forests. *Funct. Ecol.* **35**, 2190–2204 (2021).
30. Brown, J. H., Gillooly, J. F., Allen, A. P., Savage, V. M. & West, G. B. Toward a metabolic theory of ecology. *Ecology* **85**, 1771–1789 (2004).
31. Anderson, J. P. E. & Domsch, K. H. A physiological method for the quantitative measurement of microbial biomass in soils. *Soil Biol. Biochem.* **10**, 215–221 (1978).
32. Xu, X. *et al.* Global Pattern and Controls of Soil Microbial Metabolic Quotient. *Ecol. Monogr.* **87**, 429–441 (2017).
33. Sakamoto, K. & Oba, Y. Effect of fungal to bacterial biomass ratio on the relationship between CO2 evolution and total soil microbial biomass. *Biol. Fertil. Soils* **17**, 39–44 (1994).
34. Bååth, E. & Anderson, T. H. Comparison of soil fungal/bacterial ratios in a pH gradient using physiological and PLFA-based techniques. *Soil Biol. Biochem.* **35**, 955–963 (2003).
35. Sandler, S. I. & Orbey, H. On the thermodynamics of microbial growth processes. *Biotechnol. Bioeng.* **38**, 697–718 (1991).
36. Jochum, M. *et al.* Decreasing Stoichiometric Resource Quality Drives Compensatory Feeding across Trophic Levels in Tropical Litter Invertebrate Communities. *Am. Nat.* **190**, 131–143 (2017).
37. Lang, B., Ehnes, R. B., Brose, U. & Rall, B. C. Temperature and consumer type dependencies of energy flows in natural communities. *Oikos* **126**, 1717–1725 (2017).
38. Wambsganss, J., Beyer, F., Freschet, G. T., Scherer-Lorenzen, M. & Bauhus, J. Tree species mixing reduces biomass but increases length of absorptive fine roots in European forests. *J. Ecol.* **109**, 2678–2691 (2021).
39. Wambsganss, J. *et al.* Tree species mixing causes a shift in fine-root soil exploitation strategies across European forests. *Funct. Ecol.* **35**, 1886–1902 (2021).
40. McCormack, M. L. *et al.* Redefining fine roots improves understanding of below-ground contributions to terrestrial biosphere processes. *New Phytol.* **207**, 505–518 (2015).
41. Yuan, Z. Y., Chen, H. Y. H. & Reich, P. B. Global-scale latitudinal patterns of plant fine-root nitrogen and phosphorus. *Nat. Commun.* **2**, 344 (2011).
42. Dawud, S. M. *et al.* Is Tree Species Diversity or Species Identity the More Important Driver of Soil Carbon Stocks, C/N Ratio, and pH? *Ecosystems* **19**, 645–660 (2016).
43. Potapov, A. M. Multifunctionality of belowground food webs: resource, size and spatial energy channels. *Biol. Rev.* **97**, 1691–1711 (2022).
44. Barnes, A. D. *et al.* Consequences of tropical land use for multitrophic biodiversity and ecosystem functioning. *Nat. Commun.* **5**, 5351 (2014).
45. Yao, H., Chapman, S. J., Thornton, B. & Paterson, E. 13C PLFAs: a key to open the soil microbial black box? *Plant Soil* **392**, 3–15 (2015).

46. Högberg, P. *et al.* High temporal resolution tracing of photosynthate carbon from the tree canopy to forest soil microorganisms. *New Phytol.* **177**, 220–228 (2008).
47. Kaiser, C. *et al.* Exploring the transfer of recent plant photosynthates to soil microbes: mycorrhizal pathway vs direct root exudation. *New Phytol.* **205**, 1537–1551 (2015).
48. Kramer, C. & Gleixner, G. Soil organic matter in soil depth profiles: Distinct carbon preferences of microbial groups during carbon transformation. *Soil Biol. Biochem.* **40**, 425–433 (2008).
49. Treonis, A. M. *et al.* Identification of groups of metabolically-active rhizosphere microorganisms by stable isotope probing of PLFAs. *Soil Biol. Biochem.* **36**, 533–537 (2004).
50. Ling, N., Wang, T. & Kuzyakov, Y. Rhizosphere bacteriome structure and functions. *Nat. Commun.* **13**, 836 (2022).
51. Lu, W. *et al.* Impact of vegetation community on litter decomposition: Evidence from a reciprocal transplant study with 13C labeled plant litter. *Soil Biol. Biochem.* **112**, 248–257 (2017).
52. Lyu, M. *et al.* Simulated leaf litter addition causes opposite priming effects on natural forest and plantation soils. *Biol. Fertil. Soils* **54**, 925–934 (2018).
53. Shahzad, T. *et al.* Contribution of exudates, arbuscular mycorrhizal fungi and litter depositions to the rhizosphere priming effect induced by grassland species. *Soil Biol. Biochem.* **80**, 146–155 (2015).
54. Paterson, E., Sim, A., Osborne, S. M. & Murray, P. J. Long-term exclusion of plant-inputs to soil reduces the functional capacity of microbial communities to mineralise recalcitrant root-derived carbon sources. *Soil Biol. Biochem.* **43**, 1873–1880 (2011).
55. Brose, U. *et al.* Predator traits determine food-web architecture across ecosystems. *Nat. Ecol. Evol.* **3**, 919–927 (2019).
56. Barnes, A. D. *et al.* Energy Flux: The Link between Multitrophic Biodiversity and Ecosystem Functioning. *Trends Ecol. Evol.* **33**, 186–197 (2018).
57. Gauzens, B. *et al.* fluxweb: An R package to easily estimate energy fluxes in food webs. *Methods Ecol. Evol.* **10**, 270–279 (2019).
58. Byrnes, J. E. K. *et al.* Investigating the relationship between biodiversity and ecosystem multifunctionality: challenges and solutions. *Methods Ecol. Evol.* **5**, 111–124 (2014).
59. Byrnes, J. multifunc: Analysis of ecological drivers on ecosystem multifunctionality R package version 0.6. 2. *R Found. Stat. Comput. Vienna* (2014).
60. Pérez-Harguindeguy, N. *et al.* New handbook for standardised measurement of plant functional traits worldwide. *Aust. J. Bot.* **61**, 167–234 (2013).
61. Kattge, J. *et al.* TRY plant trait database – enhanced coverage and open access. *Glob. Change Biol.* **26**, 119–188 (2020).
62. Laliberté, E. & Legendre, P. A distance-based framework for measuring functional diversity from multiple traits. *Ecology* **91**, 299–305 (2010).
63. Garnier, E. *et al.* Plant functional markers capture ecosystem properties during secondary succession. *Ecology* **85**, 2630–2637 (2004).
64. Revelle, W. *Psych: Procedures for Psychological, Psychometric, and Personality Research*. (2020).
65. Weigelt, A. *et al.* An integrated framework of plant form and function: the belowground perspective. *New Phytol.* **232**, 42–59 (2021).
66. Jucker, T., Bouriaud, O., Avacaritei, D. & Coomes, D. A. Stabilizing effects of diversity on aboveground wood production in forest ecosystems: linking patterns and processes. *Ecol. Lett.* **17**, 1560–1569 (2014).
67. Pollastrini, M. *et al.* Taxonomic and ecological relevance of the chlorophyll a fluorescence signature of tree species in mixed European forests. *New Phytol.* **212**, 51–65 (2016).

68. Joly, F.-X. *et al.* Tree species diversity affects decomposition through modified micro-environmental conditions across European forests. *New Phytol.* **214**, 1281–1293 (2017).
69. Ampoorter, E. *et al.* Driving mechanisms of overstorey–understorey diversity relationships in European forests. *Perspect. Plant Ecol. Evol. Syst.* **19**, 21–29 (2016).
70. Fick, S. E. & Hijmans, R. J. WorldClim 2: new 1-km spatial resolution climate surfaces for global land areas. *Int. J. Climatol.* **37**, 4302–4315 (2017).
71. Yang, X. *et al.* Determination of Soil Texture by Laser Diffraction Method. *Soil Sci. Soc. Am. J.* **79**, 1556–1566 (2015).
72. Wild, J. *et al.* Climate at ecologically relevant scales: A new temperature and soil moisture logger for long-term microclimate measurement. *Agric. For. Meteorol.* **268**, 40–47 (2019).
73. Josse, J., Chavent, M., Liquet, B. & Husson, F. Handling missing values with regularized iterative multiple correspondence analysis. *J. Classif.* **29**, 91–116 (2012).
74. Husson, F. & Josse, J. missMDA: Handling missing values with/in multivariate data analysis (principal component methods). (2014).
75. Van Soest, P. J. & Wine, R. H. Use of detergents in the analysis of fibrous feeds. IV. Determination of plant cell-wall constituents. *J. Assoc. Off. Anal. Chem.* **50**, 50–55 (1967).
76. Gessner, M. O. & Steiner, D. Acid Butanol Assay for Proanthocyanidins (Condensed Tannins). in *Methods to Study Litter Decomposition : A Practical Guide* (eds. Graça, M. A. S., Bärlocher, F. & Gessner, M. O.) 107–114 (Springer, Netherlands, 2005).
77. Cornelissen, J. H. C. *et al.* Foliar pH as a new plant trait: can it explain variation in foliar chemistry and carbon cycling processes among subarctic plant species and types? *Oecologia* **147**, 315–326 (2006).
78. Freschet, G. T., Aerts, R. & Cornelissen, J. H. C. A plant economics spectrum of litter decomposability. *Funct. Ecol.* **26**, 56–65 (2012).
79. Joly, F.-X., Scherer-Lorenzen, M. & Hättenschwiler, S. Resolving the intricate role of climate in litter decomposition. *Nat. Ecol. Evol.* **7**, 214–223 (2023).
80. Goodrich, B., Gabry, J., Ali, I. & Brilleman, S. rstanarm: Bayesian applied regression modeling via Stan. R package version 2.21.4. (2023).
81. Gelman, A. *et al. Bayesian Data Analysis*. (CRC Press, 2013).
82. Zuur, A. F., Ieno, E. N. & Elphick, C. S. A protocol for data exploration to avoid common statistical problems. *Methods Ecol. Evol.* **1**, 3–14 (2010).
83. Lüdecke, D., Ben-Shachar, M. S., Patil, I., Waggoner, P. & Makowski, D. performance: An R package for assessment, comparison and testing of statistical models. *J. Open Source Softw.* **6**, (2021).
84. Ives, A. R. For testing the significance of regression coefficients, go ahead and log-transform count data. *Methods Ecol. Evol.* **6**, 828–835 (2015).
85. Barr, D. J., Levy, R., Scheepers, C. & Tily, H. J. Random effects structure for confirmatory hypothesis testing: Keep it maximal. *J. Mem. Lang.* **68**, 255–278 (2013).
86. Schielzeth, H. & Forstmeier, W. Conclusions beyond support: overconfident estimates in mixed models. *Behav. Ecol.* **20**, 416–420 (2009).
87. Matuschek, H., Kliegl, R., Vasishth, S., Baayen, H. & Bates, D. Balancing Type I error and power in linear mixed models. *J. Mem. Lang.* **94**, 305–315 (2017).
88. Nakagawa, S. & Cuthill, I. C. Effect size, confidence interval and statistical significance: a practical guide for biologists. *Biol. Rev.* **82**, 591–605 (2007).
89. Gronau, Q. F., Singmann, H. & Wagenmakers, E.-J. bridgesampling: An R package for estimating normalizing constants. *ArXiv Prepr. ArXiv171008162* (2017).
90. Vehtari, A., Gelman, A. & Gabry, J. Practical Bayesian model evaluation using leave-one-out cross-validation and WAIC. *Stat. Comput.* **27**, 1413–1432 (2017).
91. Vehtari, A. *et al.* LOO: Efficient leave-one-out cross-validation and WAIC for Bayesian models (2019). *R Package Version* **2**, (2022).

92. Shi, H. & Yin, G. Reconnecting p-Value and Posterior Probability Under One- and Two-Sided Tests. *Am. Stat.* **75**, 265–275 (2021).
93. Shipley, B. Confirmatory path analysis in a generalized multilevel context. *Ecology* **90**, 363–368 (2009).
94. Lefcheck, J. S. piecewiseSEM: Piecewise structural equation modeling in R for ecology, evolution, and systematics. *Methods Ecol. Evol.* **7**, 573–579 (2016).
95. Joly, F.-X. *et al.* Detritivore conversion of litter into faeces accelerates organic matter turnover. *Commun. Biol.* **3**, 660 (2020).
96. Ponge, J. F. Humus forms in terrestrial ecosystems: a framework to biodiversity. *Soil Biol. Biochem.* **35**, 935–945 (2003).
97. Zanella, A. *et al.* A European morpho-functional classification of humus forms. *Geoderma* **164**, 138–145 (2011).
98. Schwarz, B. *et al.* Warming alters energetic structure and function but not resilience of soil food webs. *Nat. Clim. Change* **7**, 895–900 (2017).
99. Wan, B. *et al.* Energy flux across multitrophic levels drives ecosystem multifunctionality: Evidence from nematode food webs. *Soil Biol. Biochem.* **169**, 108656 (2022).
100. Ehnes, R. B., Rall, B. C. & Brose, U. Phylogenetic grouping, curvature and metabolic scaling in terrestrial invertebrates. *Ecol. Lett.* **14**, 993–1000 (2011).
101. Gan, H. & Wickings, K. Root herbivory and soil carbon cycling: Shedding “green” light onto a “brown” world. *Soil Biol. Biochem.* **150**, 107972 (2020).
102. Harrison, X. A. *et al.* A brief introduction to mixed effects modelling and multi-model inference in ecology. *PeerJ* **6**, e4794 (2018).
